# Fully addressable, designer superstructures assembled from a single modular DNA origami

**DOI:** 10.1101/2023.09.14.557688

**Authors:** Johann M. Weck, Amelie Heuer-Jungemann

## Abstract

Intricate self-organization is essential in many biological processes, underpinning vital functions and interactions. In an effort to mimic such processes, synthetic biology aims to engineer dynamic structures with controllable functions using nanotechnological tools. A key requirement of engineered building blocks is the ability to assemble and disassemble hierarchically with precision. Using the DNA origami technique, we here present the moDON, a modular DNA origami nanostructure, which is capable of assembling into almost 20 000 diverse monomers, forming complex and controlled superstructures in three dimensions. While shape and addressability of DNA origami are nearly arbitrary, its overall size is limited by the scaffold size. Previous methods of extending the size of DNA origami (e.g. hierarchical assembly, modified scaffolds, etc.), either led to loss over control of shape and addressability beyond monomers or to proportionally increased cost and design effort. With the moDON we were able to overcome both issues. The modular design combines xy- and z-plane assembly methods, enabling the construction of finite and periodic structures beyond 1 GDa. We demonstrate xy-z orthogonality, by enabling controlled selective or parallel assembly and disassembly via distinct orthogonal triggers. The kinetic profile of assembly and disassembly aligns with biological time scales, paving the way for applications in dynamic nanomachinery and advanced biomaterials. Finally, we showcase the conjugation of gold nanoparticles to specific positions within superstructures, underscoring the efficacy of this approach for creating intricate and orthogonal nanoscale architectures with preserved site-specific addressability. The moDON thus offers an efficient, cost-effective solution for constructing large, precisely organized, and fully addressable structures with vast potential in synthetic cellular systems design.

## INTRODUCTION

Nature equipped cellular systems with a plurality of functions for homeostasis, reproduction, and interaction with its environment. All of these rely upon the capacity of biological matter to self-organize into higher-order structures. Protein complexes fulfill these functional and organizational roles, highly selectively assembling and disassembling into dimers (*e*.*g*. reverse transcriptase), small multimers (*e*.*g*. the apoptosome), or large multimers made up of many subunits, such as microtubules and actin filaments making up the cytoskeleton^1,2^. Their size ranges from a few hundred nanometers (nm) to several micrometers (µm), spanning the relevant distances for fulfilling their function, despite being assembled from well-defined subunits of only a few nm in size^3^. As synthetic biology progresses, researchers aim to mimic said functions, both on a molecular and on a cellular level. Therefore the need for building blocks capable of controllably assembling and disassembling into large multimeric structures on biologically relevant time scales has become more urgent.

Structural DNA nanotechnology and especially the DNA origami^4^ technique, where a long, circular scaffold strand, extracted from the M13mp18 bacteriophage, is folded into any desired 2D or 3D shape using short synthetic staple strands, presents an indispensable tool for nanoscale engineering. DNA origami nanostructures (DONs) ranging from just a few nm to over a hundred nm can be easily designed and synthesized, with freely available CAD software such as caDNAno.^5^ To date a large variety of DONs have been published, ranging from the simplest 2D structures^4^ to functional DNA origami motors^6^. A very attractive feature of DONs is their complete addressability for guest molecule placement with base pair (bp) precision. This feature has been employed to position various functional moieties ranging from fluorophores to nanoparticles or proteins, enabling the formation of fluorescent nanorulers,^7^ plasmonic nanosensors,^8,9^ nm-sized force sensors,^10,11^ or to study complex biological questions.^12-14^

Despite all of these advantages, a bottleneck in DON design is the overall final size limit, determined by the scaffold strand (the M13mp18 genome is made up of 7249 nt), which is the common factor in almost all DNA origami structures. Although different insertions have led to scaffold sizes of close to 9000 bases, the overall size increase in 3D DONs is marginal. Therefore, a variety of different strategies have been reported to increase DON sizes. These include the use of a lambda/M13 hybrid phage as a scaffold source^15^, the development of orthogonal scaffold strands^16^, the use of unscaffolded structures^17-19^, origami slats^20,21^, or hierarchical assembly into finite^22,23^ or periodic^23,24^ structures. Each of these methods has at least one major drawback: With the lambda/M13 hybrid phage, synthesis of scaffolds larger than 50 000 bases^15^, equivalent to ∼six regular origami was achieved. A similar size was achieved with superstructures from orthogonal scaffolds, reaching sizes equivalent to five regular origami. However, both approaches have similar bottlenecks: Firstly, the effort for scaffold production is increased tremendously and secondly, the unique scaffold sequence requires the same amount of unique staples to be designed and synthesized individually, increasing effort and monetary cost proportionally to superstructure size. The latter problem also affects unscaffolded nanostructures, employing single-stranded tile assembly, where every ssDNA tile requires individual design and sythesis. Nonetheless, using 10 000 unique ssDNA tiles, superstructures in the Gigadalton (GDa) scale have been constructed^19^. The largest superstructures reported to date were constructed using so-called DNA origami slats^21^, through connection of several long 6 or 12 helix bundles (HBs) via well positioned, complementary ssDNA handles. Although these structures reached several GDa in size and their synthesis is comparably cost effective, assembly has only thus far only been shown in 2D. Almost similar sizes can be reached through hierarchical assembly, where a multitude of either the same or different DON monomers are assembled into a large superstructure. This approach has led to structures measuring 0.5 µm^2^ (2D), depicting a μm-sized image of the Mona Lisa^25^, while 3D structures could reach up to the GDa scale^22^. The high structural control and potential size make it the most feasible approach for the formation of superstructures. Nevertheless, design and construction are faced with different challenges.

For the design of large hierarchically assembled superstructures, connectivity always faces one of two problems: On the one hand, assemblies of self-complementary DONs lead to repetitive, homomultimeric finite^22^ or periodic^23,24^ superstructures. This approach is simple and cost-effective, as in principle only one type of monomer is required. However, it has the great disadvantage of an overall loss of structural control of the superstructure beyond the repetitive subunit, as only form and size of the subunit is controlled, yet superstructure formation occurs without control over the final structure. Consequently, it is impossible to controllably retain the site-specific addressability of the structure beyond the single monomer or subunit. On the other hand, assemblies made up of many different DONs, each complementary to one another, but not themselves, yield finite, heteromultimeric superstructures^6,16^. With this approach the structural control over the superstructure is high and the site-specific addressability is retained. However, the downsides are the requirement for a multitude of different DONs, leading to high design effort and increasing costs, proportional to superstructure size, and the need to be assembled in several steps, resulting in a tremendous decrease in overall yield.

Therefore, currently, a concept, combining both approaches, *i*.*e*. using only one type of DON that can be modularly modified (low cost) and controllably assembled into different superstructures using specific connections (high structural control and retained controlled addressability) is missing. Such a concept could alleviate the issues associated with traditional hierarchical assembly and allow for the controlled assembly (and disassembly) of large superstructures, or a multitude of orthogonal smaller structures, with retained site-specific addressability across the entire superstructure. A simple and cost-effective approach to constructing large, rigid structures with high orthogonality to other connections, would also be a much-needed addition to the toolkit of synthetic cellular systems design. In response to this challenge, we here introduce the moDON, a single modular DNA origami formed from one set of staples, able to controllably assemble and disassemble into large superstructures in the xy- as well as in the z plane. Modularity in xy is introduced through a design method of modular scaffold routing, changing the scaffolded structure of the origami with exchange of only a few staples, giving rise to hundreds of individual monomers. Additionally, a modular three-strand connection system in the z-direction allowed us to form ∼ 20 000 unique moDON monomers. Using this approach, we demonstrate the construction of arbitrary finite superstructures in three dimensions, as well as periodic structures with monomeric and multimeric repetitive subunits, all in one-step assemblies. Further, we show that xy- and z-connections are fully orthogonal allowing for parallel and selective assembly and disassembly of each individual connection using different orthogonal triggers (see Figure S1 for overview). We also give insights into the underlying dynamics, showing quick assembly and disassembly on biologically relevant time scales. Finally, we show that the site-specific addressability of each monomer in the superstructure is fully retained by placing gold nanoparticles (Au NPs) at specific positions in the superstructures, demonstrating a highly controllable, efficient and cost-effective strategy for the formation of large and orthogonal superstructures.

## RESULTS AND DISCUSSION

### Design & Characterization of the moDON

The basic design of the moDON is a 78 HB, based on the honeycomb lattice design. As can be seen in **Scheme 1**, the moDON monomer consists of one large core section (indicated in white), containing six extended helices on the left and right side of the structure (indicated in turquois), as well as a modular shell (indicated in yellow). The scaffold in the core part was layered evenly to facilitate folding (see Figure S2). To counteract any residual twist in the structure^22,26^, a deletion was introduced every 126 bp, thus adjusting the twist to 34.56°/bp over the whole structure (see Figure S3). The unused 116 nt long scaffold loop was positioned in the middle of the structure, pointing into the 6 HB cavities, to prevent any undesired interactions.

The different approaches to modularity and assembly connectivity of the moDON in both the xy- and the z-direction are illustrated in **Scheme 1**. Connection in the xy-direction was achieved by accurately fitting modularly changeable protrusions and indentations of helix duplexes (see yellow helices in **Scheme 1b**), while the connection in the z-direction was achieved by employing a three-strand system, including a connector-strand, continuing the ssDNA scaffold route in the design (see turquois helices in **Scheme 1c**). In the following sections both connection strategies will be introduced in detail.

### Modularity and Assembly in xy-Direction

Modularity in the xy-direction was introduced by re-routing the scaffold on the outer shell of the moDON (yellow helices in **Scheme 1**), without changing the core structure (white helices). To minimize the number of nt in the core structure involved in re-routing of the shell, certain helices were chosen as so-called hinge helices (HH) between the stable core and the modular shell (see Figures S2, S4, S5). The modular shell section of the moDON consists of helices extending out from the HHs. To each side of one HH, two additional helices (four in total), extend out, forming patterns of protrusions or indentations that make up an xy-connection site as depicted in yellow in **Scheme 1a** (and S2c). For the design of these connection sites, rules outlined in refs. ^23,27^ were adapted (*i*.*e*. a minimum length of 21 nt, as well as no staple/scaffold cross-overs close to end positions). Special attention was paid to avoid point symmetries in the design of the connection sites, which could lead to specific, but 180° turned connections. The twelve different active xy-connection sites that can possibly be formed were designated as α-ζ and α*-ζ* for the respective complementary sites. Each of those could also be selectively deactivated/passivated. For ease of illustration, two different sets of configurations are depicted in **Scheme 1a**, with configuration 1 displaying connection sites α, β and γ as well as their complements α*-γ*, while configuration 2 displays connection sites δ, ε and ζ as well as their complements. It is important to note that a moDON can have a combination of connection sites from both configurations. However, α of the first configuration and δ of the second configuration share the same position in the moDON, thus not allowing them to be present on the same structure. The same holds true for β and ε, as well as for γ and ζ and all respective complements (an overview is given in Figure S6). Mutual exclusivity was ensured by checking all possible permutations of each connection site in all configurations. In order to obtain the different configurations, the scaffold routing in the HH and the adjacent helix loops needed to be adjusted from traditional DNA origami scaffold routing designs (see **Scheme 1a**, and S4).

As the scaffold is circular by default, and in order for only the shell, but not the core, to be modular, a complex scaffold routing approach was devised. In traditional DNA origami design, the scaffold cross-overs, connecting two adjacent helices, are generally placed at alternating positions between the ends of the helices and towards the middle of the helices (*cf*. Figure S2a). Constraints from the design of the modular shell and the requirement for simplicity in scaffold layering required a deviation from this traditional routing. To allow for modularity, the shell was designed from closed scaffold loops, adjacent to and protruding from each HH, forming two short, parallel helices (illustrated in Figures S2b, S2c, and S4). The scaffold cross-overs between these two helices in the loop were placed at the ends of the helices, while the cross-over to the HH was placed towards the middle (Figure S2b). Connection of the HH to the core structure therefore again was required to be at the helical ends, interrupting the alternating middle-end-middle scaffold cross-overs from helix 0 (H0) to H22 (for helix numeration see Figure S5a). By design, the cross-over between H22 and H23 was required to be placed in the middle of the structure, as H22 connects to HH75 at the helix ends. This was adjusted for all hinge motifs, allowing free modularity for all connections in the xy-direction, through addition of the appropriate staple subset (see **Scheme 1a** and S6). In order to correct for rounding errors when determining helical cross-over positions in these modular parts, insertions were added into the design as required (Figure S3, and S5b).

Connection sites were designed to be orthogonal across both configurations. Since connection sites are complementary to the respective opposite connection site on the moDON, the changeable sections are positioned not point-symmetrically, but mirror-symmetrically (see Figure S6). Consequently, this leads to an uneven distribution of the number of configurations across the HHs, as HH65 only contains one configuration, whereas HH50 contains four configurations, of all permutations between the adjacent connection sites γ* or ζ* and α or δ. A total of 13 connection site mixes is therefore required to form all scaffold configurations (see Table S1). oxDNA simulations of moDONs in configurations 1 and 2 were used to predict correct folding and stability (Figure S7 and S8), which was further confirmed experimentally by agarose gel electrophoresis (AGE) and transmission electron microscopy (TEM) (Figure S9).

**Scheme 1:**
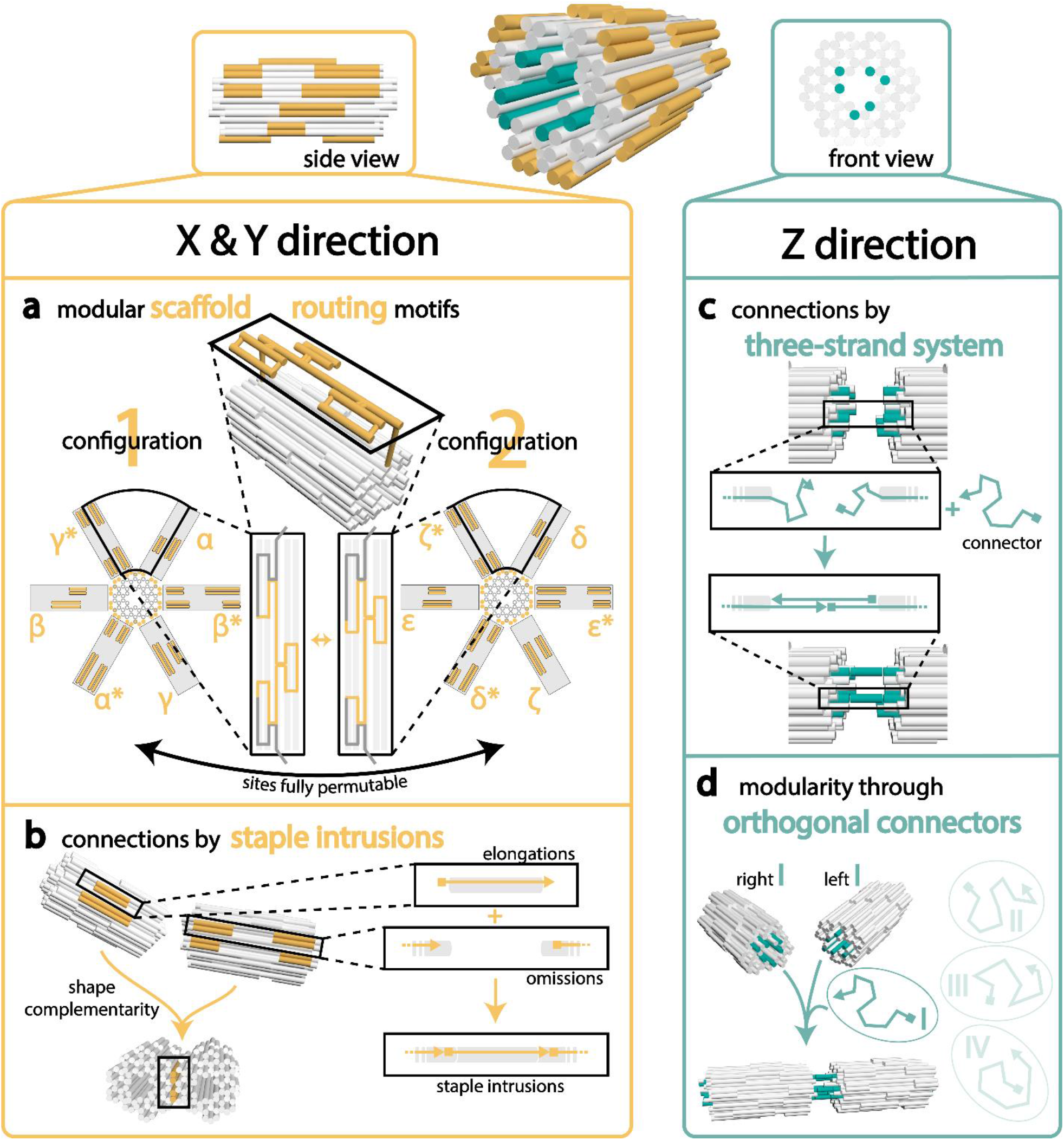
Connectivity and modularity of the moDON is achieved by employing two orthogonal connection strategies, each with a unique modularity. (**a**) The xy-connections contain modular scaffold motifs, secluding the modular shell from the stable core structure: The scaffold performs a cross-over from the core structure to the HH (yellow) on the helix ends. Whereas the scaffold cross-over from the HH to the connection sites (yellow) loop out from the middle of the helix. The ends remain stable (grey), and the middle part of the connection site is configurable with just a few staples. (**b**) In the xy-direction the connection is achieved by shape complementary sites and stabilized by short staple intrusions yielding specificity. (**c**) The connection in z-direction (turquois) is achieved via a three-strand-system, with ssDNA handles protruding from the left and right ends of each monomer, each complementary to one half of an external connector strand. Handles on the right side of the moDON are elongated at the 3’ end, and handles at the left side are elongated at the 5’ end, making the connection directional. (**d**) A set of four orthogonal connectors and handles on the moDON achieves modularity in the z-direction.

To ensure even more control over xy-connected assemblies, in addition to shape complementarity in the connection sites, we also introduced short staple intrusions (see **Scheme 1b** and S3). Furthermore, connection sites could also be deactivated/passivated by inclusion of 5 Thymine (T) staple overhangs (Figure S3). Together, this would allow us to mix a manifold of different moDON monomers, with specific connections sites, which could assemble into distinct superstructures (concept see **Figure 1a**). To experimentally test the modularity and connectivity in the xy-direction, we initially formed dimers and trimers from all possible complementary monomers (α-ζ and α*-ζ*). Analysis by AGE revealed a clear shift in electrophoretic mobility for dimers and trimers, compared to monomers, proportional to superstructure size, suggesting successful assembly (Figures S10, S11). This was further confirmed by TEM imaging, which showed well-assembled dimers and trimers as illustrated in **Figure 1d**. Impressively, the assembly yield, as determined from the gel band intensities (Figures S10-11) was found to be near quantitative, demonstrating the effectiveness of the approach. After the successful assembly of dimers and trimers, we next sought to form larger structures and therefore constructed tetramers and hexamers. AGE analysis of both structures again successfully showed the expected reduction in electrophoretic mobility dependent on superstructure size as well as near quantitative assembly yields for both structures, (**Figures 1b** and S12-13). Further demonstrating the versatility of the moDONs xy-connection approach, we next constructed a multitude of arbitrary structures up to heptamers of different configurations. As can be seen in **Figure 1d** (and Figures S14-17) all structures assembled exceptionally well, showcasing the versatility of our approach. We also constructed periodic, homo-multimeric xy-lattices with a unit cell of 2 moDONs, resulting in 2 dimensional structures spanning more than 36 000 nm^2^ (Figure S18) made up of 46 monomers. We summarize, that one-step connections of a total of seven unique moDON monomers (from six unique connections) at the same time are possible (see Figure S17). Modularity allows for the doubling of connections in each direction, and thus increases the configuration space for superstructures 8-fold.

**Figure 1:**
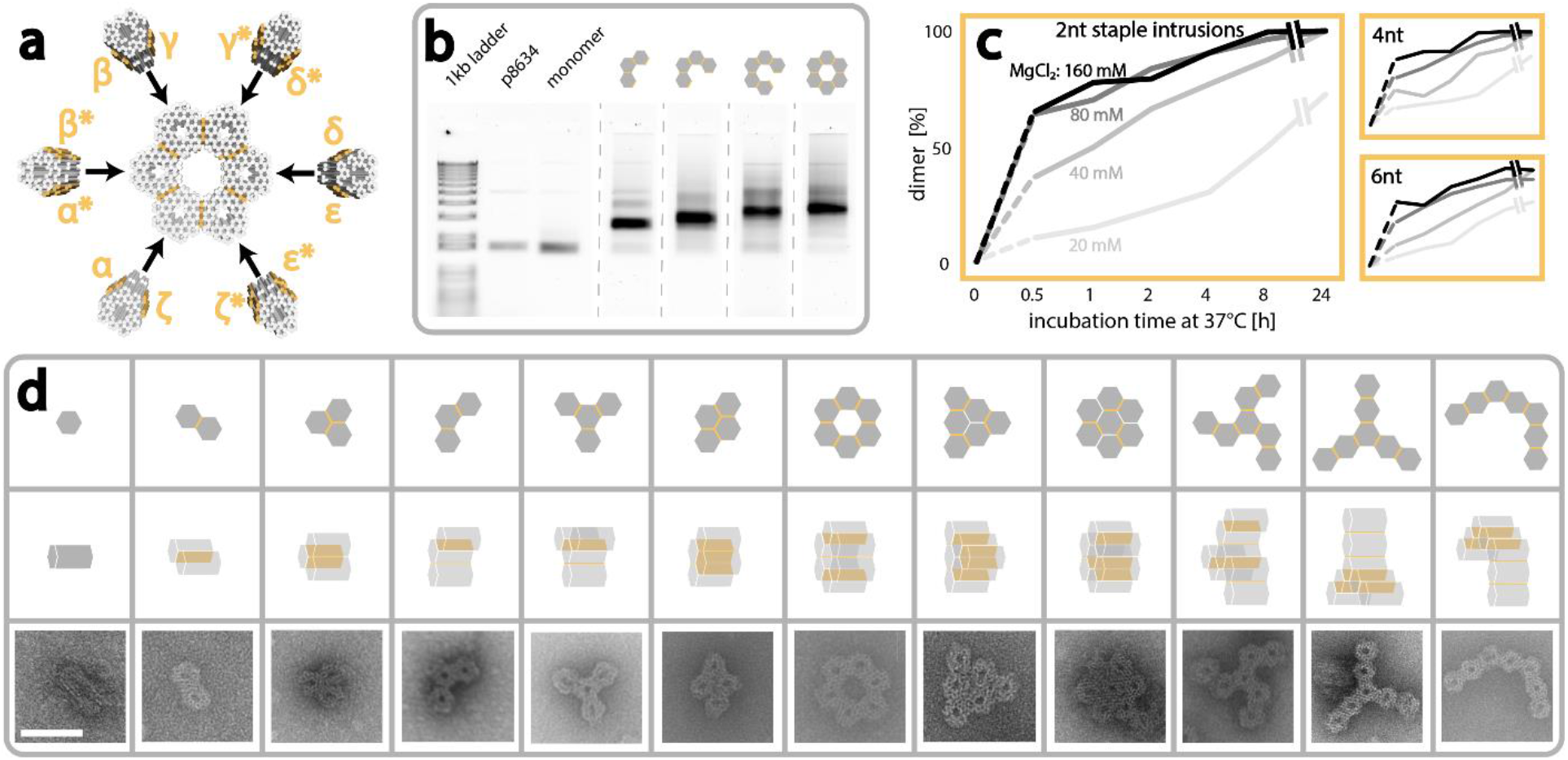
Superstructures in xy-direction are constructed from a multitude of moDON monomers. Active connection sites are indicated in yellow. (**a**) Exemplary construction of a hexameric ring superstructure from 6 different moDON configurations (βγ, γ*δ*, δε, ε*ζ*, ζα, α*β*) in a one-step-reaction. moDONs γ*δ* and ζα are cross configuration structures. (**b**) Gel image of hexameric ring construction and intermediate multimers. The monomer displays a higher electrophoretic mobility compared to the scaffold, p8634. Higher order multimeric structures display slower electrophoretic mobilities proportional to superstructure size. (**c**) moDON dimerization yield at different MgCl_2_ concentrations over 24 h showing that full dimerization is only achieved at 40 mM or higher MgCl_2_ concentration. (**d**) Exemplary xy-direction superstructures made from various moDONs in one-step-reactions. Pictograms indicate designed structure. Scale bar is 50 nm and holds for all TEM micrographs.

After successful assembly of a great variety of different higher order structures, we next sought to investigate the influence of different experimental parameters and design choices on superstructure assembly in the xy-direction in more detail. In brief, we examined a parameter tensor of (i) incubation times from 0.5 to 24 h, (ii) MgCl_2_ concentrations from 20 to 160 mM, and (iii) the length of staple intrusions of 2, 4, and 6 nt respectively. Analysis by AGE revealed that both the incubation time as well as the MgCl_2_ concentration in the buffer had a large influence on dimerization ratio for all lengths of staple intrusions (see Figures S19-20). Analyses of the corresponding band intensities (**Figure 1c**) revealed that longer incubation times as well as higher concentrations of MgCl_2_, resulted in an increase in the yield of dimerized moDONs. At 20 mM MgCl_2_ the dimerization yield was only ∼10 % after 0.5 h and ultimately only rose to 70 % after 24 h. It was thus deemed to not be sufficient to connect moDONs within a reasonable amount of time. Dimerization yields were improved by increasing the MgCl_2_ concentration to 40 mM, reaching full dimerization after 24 h of incubation. It should be noted that a MgCl_2_ concentration of 160 mM did not further increase the speed of dimerization, but led to increased smearing of the bands in the gel (Figures S19-20). We also note, that even this high concentration of MgCl_2_ did not lead to undesired multimerization, which we attribute to the compact design and thorough passivation. As can be seen from **Figure 1c**, the abovementioned observations hold true for all lengths of staple intrusions, suggesting that their length does not play a vital role in the assembly process. Summarizing, we can identify a MgCl_2_ concentration of 40 mM or higher as the main trigger for assembly in the xy-direction.

### Modularity and Assembly in z-Direction

After successful assembly of moDON superstructures in the xy-direction, we next tested the connection strategy for the z-direction. Connections in the z-direction were designed with a three-strand system, orthogonal to the xy-connectivity. For this, left and right sides of the moDON were functionalized independently, resulting in the ability to connect to two different monomers at a time as illustrated in **Scheme 1c,d**. In order to allow for directional assembly, six helices were designated as connection sites, depicted in turquois in **Scheme 1**. Staples of connector helices on the left side of the structure were elongated by an additional 10 nt at their 5’ end and staples of connector helices on the right side of the structure were elongated by 11 nt at their 3’ end. Upon the addition of a 21 nt long, complementary connector strand, a connection would be formed resulting in two full helical turns between monomers, continuing the helical pacing in the respective moDON monomers. To ensure stability and rigidity, each connection site was designed to consist of six handles of the same sequence. We specifically chose short lengths of extending staples of only 10/11 nt in order to prevent unintentional passivation of the connection site, enabling for incompletely paired connectors to detach from the handle. To further minimize undesired hybridization, a three-letter alphabet of Cytosin, Adenin, and Thymine was used in the connector design. In total, we designed four different connectors: I, II, III, and IV (see **Scheme 1d** and Table S15). Their mutual orthogonality, as well as the orthogonality towards all moDON connection sites were verified *in silico* by NUPACK analysis (Figure S21).

Similar to the assembly in the xy-direction, we initially experimentally investigated the formation of dimers with all possible permutations. AGE revealed successful dimer formation only upon addition of the correct connector strand, confirming mutual orthogonality of the connectors (Figure S22). Subsequent analysis of structures by TEM (Figure S23) and AGE (Figure S24a) revealed a near quantitative dimer yield. Interestingly, when testing various connector handle to connector ratios, we found that full dimerization was achieved even with a 1:1 ratio as determined by AGE (see Figure S24a). A more detailed analysis of these structures by TEM, however, revealed that the connection between single moDONs appeared to be bent (Figure S24b). Contrastingly, when the ratio was increased to 1:4 or higher, dimers were observed to be linear (Figure S25b). We attribute this to the decreased probability of successfully forming all of the six possible connections if connector concentrations are too low, which in return leads to the z-connections acting as hinge-regions. We therefore used ratios of at least 1:4 (handle : connector) for all assemblies.

To further showcase the orthogonality and efficacy of the three-strand-connecting system, we next constructed a multitude of different structures ranging from dimers to pentamers with one specific connector strand for each connection. As can be seen from the AGE analysis in **Figure 2c** (and Figure S22), all structures formed successfully and with high yield, from one set of orthogonal moDONs (i.e. five different monomers). Higher order structures displayed proportionally slower electrophoretic mobilities, as desired. N.B.: as the mixture contained five different types of monomers, the addition of only connector I results in the formation of dimers of only two of the different monomers, with the other three types of monomers, non-complementary to connector I, remaining in the solution, resulting in two bands in the gel. The same holds true for trimer and tetramers. Only the addition of all five different connectors results in full assembly of all monomers into a pentamer with near quantitative yield (**Figures 2d** and S25, S26).

**Figure 2:**
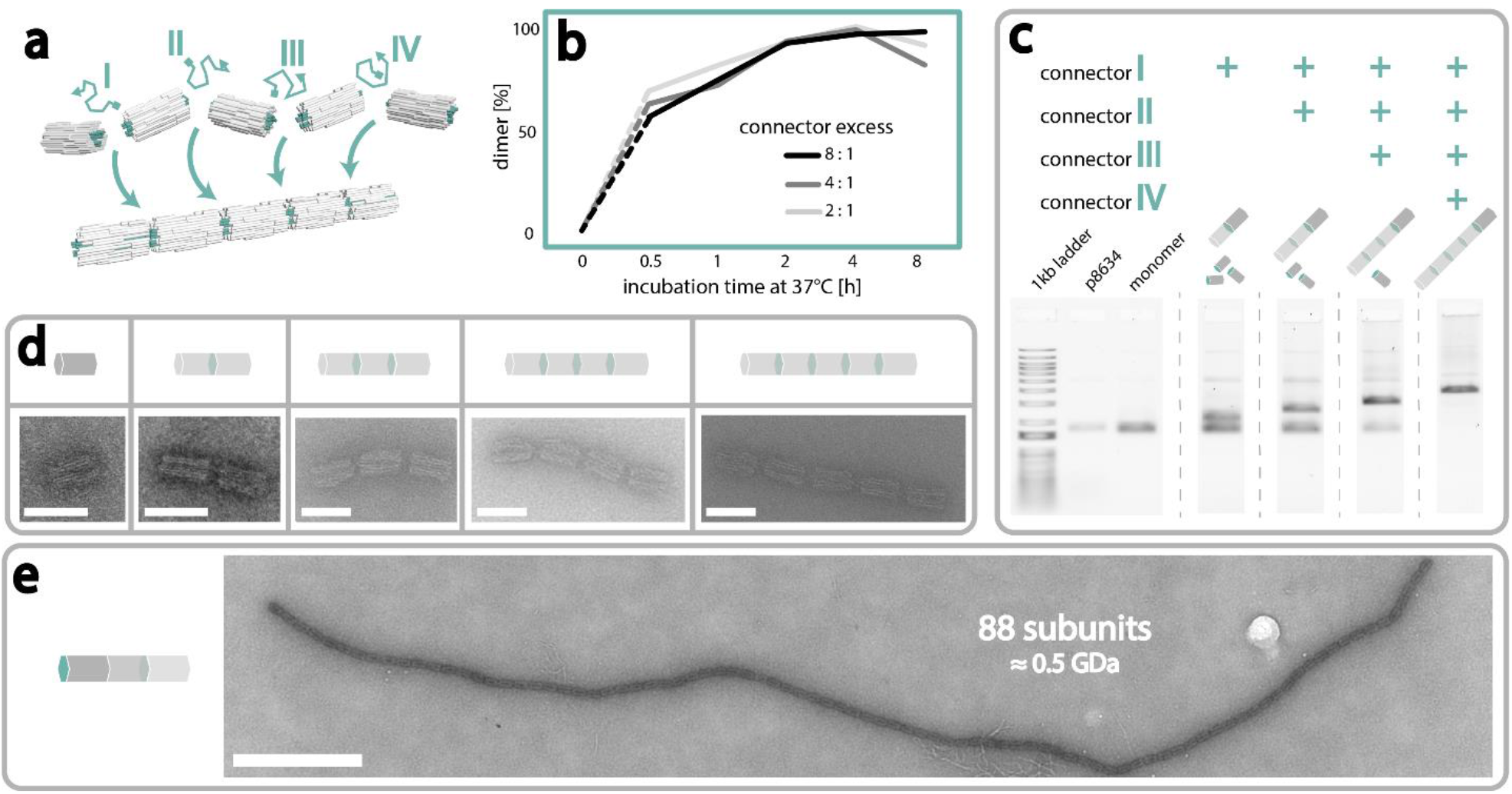
Superstructures in z-direction are constructed by connecting moDON monomers with a connector strand. (**a**) Construction of a pentamer from moDON monomers. The respective connector strand (I, II, III, or IV) hybridizes to specific ssDNA handles on a moDON. (**b**) Dimerization yield dependent on connector excess over time. In each case more than 50 % of moDONs were dimerized after 0.5 h. Increased connector excess did not yield faster dimerization. (**c**) Gel image of z-directional connections forming from the same moDON mix of five different monomers, with stepwise addition of the respective connectors. Electrophoretic mobility is reduced proportional to the degree of multimerization. Shown below the formed structures are the orthogonal monomers not involved in the connection. (**d**) Exemplary z-direction superstructures made from various moDONs in a one-step-reaction. Pictogram indicates designed structure; turquois indicates connection sites. (**e**) Assembly of a periodic tube of moDON in z-direction with a monomeric subunit. Scale bars in (**d**) are 50 nm, scale bar in (**e**) is 500 nm

After having successfully assembled finite structures, we consequently used our approach to construct periodic tubes from monomeric subunits. As can be seen in **Figures 2e** and S27, these tubes reached lengths of up to 88 monomers measuring > 4 µm in length, corresponding to a molecular weight of ∼0.5 GDa. With this we were further able to highlight the efficiency and orthogonality of our assembly approach in the z-direction.

We next sought to investigate the interrelation of connection speed with connector and MgCl_2_ concentration. We thus constructed a matrix of (i) incubation times, from 0.5 to 8 h, and (ii) connector to handle ratios from 2 to 8, or (iii) MgCl_2_ concentrations from 5 to 40 mM. Analysis by AGE revealed the dimerization ratio to be very fast, reaching complete dimerization already after 2 h (**Figure 2b**, S28a) independent of the connector excess. Interestingly, compared to the assembly kinetics in the xy-direction, connection in the z-direction was nearly twice as fast. We attribute this to the increased accessibility of the connection sites, at the ends of the moDON, compared to the slightly obstructed staple intrusions. Further, we suspect the repetitiveness of the connection sites to play a major role, as each connector has six potential, equal connection handles on the moDON. We additionally found that higher MgCl_2_ concentrations (40 mM) resulted in faster dimerization kinetics, reaching full dimerization after only 1 h. However, after 8 h, all MgCl_2_ concentrations of 10 mM and higher resulted in full dimer formation (see Figure S28b,c). We thus conclude, that, as could be expected, the assembly process is dominantly triggered by the presence of the connector strand, while perhaps surprisingly, the concentration of MgCl_2_ in the buffer had no significant influence (above 10 mM). Importantly, these findings confirm that the assembly trigger for z-directional connections (connector strands) is fully orthogonal to that of xy-directional assembly (high MgCl_2_ concentrations).

### Selective Assembly and Disassembly

We thus far showed that higher order structures can be reliably formed in both xy- and z-directions, with high yields. However, control of size and structure of xy- and/or z-directional superstructures constitute only the first feature of the moDON. In the following we show assembly of structures combining xy- and z-connections, and the selective disassembly thereof, as well as investigating the underlying dynamics.

As all xy-configurations and z-connections are orthogonal to one another, we hypothesized that we would be able to assemble different structures in parallel, at the same time, in the same tube. To demonstrate parallel assembly, we formed simultaneously (i) a trimer from moDONs with active α, β, and γ sides, (ii) a tetramer from moDONs with active δ, ε, and ζ sides and (iii) a pentamer in z-direction, in a one-step reaction. After addition of the respective triggers (MgCl_2_ for xy-assembly and and connector strands for z-assembly) structures were analyzed by AGE and TEM. Encouragingly, species with three different electrophoretic mobilities could be observed in the gel, corresponding to trimers, tetramers and pentamers, as desired (see Figure S29b). This was further confirmed by TEM analysis, where many structures of all three species could be seen (see **Figures 3a** and S29). This not only illustrates the amount of unique connection sites and their respective orthogonality, but also powerfully demonstrates the trigger-dependent selectivity of the assembly process, which we now explore further.

**Figure 3:**
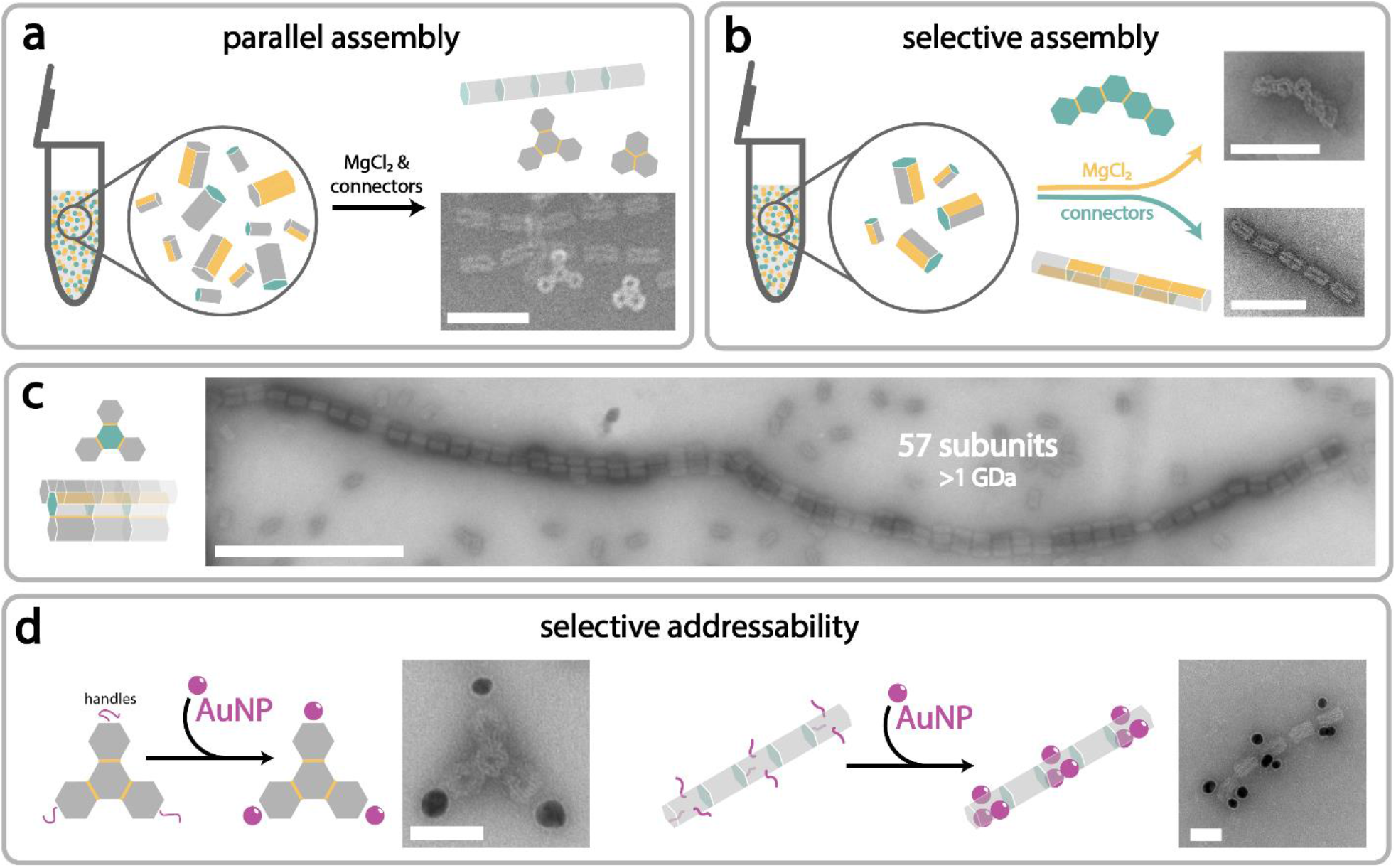
Parallel and selective assembly with retained monomer addressability: (**a**) The xy-z-orthogonality allows for parallel formation of several different structures in the same tube. xy-connection sites are indicated in yellow, z-connection sites in turquois. (**b**) From the same set of five moDON monomers, two different superstructures were selectively assembled: a distinct xy-pentamer or a z-pentamer, depending on the nature of the added assembly trigger (MgCl_2_ for xy-connections, connector strands for z-connections). (**c**) Large 3-dimensional structures are formed by combining xy- and z-connections. The tubes are constructed with a tetrameric subunit in xy-direction that can infinitively stack in the z-direction. The multimeric tubes yield sizes of up to 57 subunits, reaching over 1 GDa cumulative mass. (**d**) Addressability in moDON superstructures is retained beyond a single monomer. ssDNA handles (purple) are positioned exclusively on the vertices of the tetrameric (from xy-connections) or at the middle and outer monomers in a pentameric (from z-connections) structure, therefore controlling Au NP (purple) arrangement within the superstructure. Scale bars are 100 nm for (**a**) and (**b**), 500 nm for (**c**), and 50 nm for (d).

After having successfully established that our approach can yield several desired structures in parallel, we next sought to achieve selective assembly of different structures from the same moDONs. For this, we constructed five moDONs, each with active xy- and z-connection sites capable of forming a pentamer either in the xy- or the z-direction, depending on the respective trigger (MgCl_2_ or connectors) added. Encouragingly, we indeed found that almost exclusively one or the other trigger-specific pentamer was constructed from the same set of moDONs. As can be seen in **Figures 3b** (and S30) an increase in MgCl_2_ concentration resulted in xy-pentamers, whereas an addition of the connector strands resulted in z-pentamers. This further demonstrated the orthogonality and thus specificity of the connections, constituting the foundation for hierarchically assembling very large structures.

Until now we only showed the selective xy- or z-directional assembly of superstructures. However, due to the xy-z orthogonality, a combination of both assembly methods can be used to achieve even larger-sized superstructures. To demonstrate this, we aimed to assemble periodic structures in the z-direction with varying xy-directional subunits (trimers or tetramers) in a single assembly step. Note that only one of the moDONs in the trimeric or tetrameric subunit was designed to contain z-directional connection sites at both ends for periodic assembly (indicated in turquois in **Figure 3c**). Most previously published large hierarchically assembled superstructures were formed in multi-step assemblies, to achieve larger and larger structures. Here, trimeric or tetrameric subunits and subsequent infinite periodic structures thereof, were all successfully formed in a single step, evidenced by TEM analysis (**Figures 3c**, and S31, S32). The representative micrograph of periodic tetrameric subunit assemblies in **Figure 3c**, displays one of the largest structures we observed, which was made up of 57 tetrameric subunits, with lengths reaching almost 3 µm and a molecular weight of over 1 GDa. Although only assemblies from monomeric, trimeric and tetrameric subunits were shown, the approach can easily be extended to any other possible xy-subunit, making it a highly versatile method. Furthermore, this could also be extended to form networked structures, as found in the cytoskeleton of cellular systems.

In addition to construction of arbitrarily shaped superstructures, another desirable goal in hierarchical assembly is to retain the ability of modifying each point in the superstructure selectively. This is also often not possible in traditional hierarchical assembly approaches. Therefore, to demonstrate that using the moDON approach, targeted addressability of single moDONs within a superstructure can be retained, we selectively conjugated DNA-coated Au NPs to specific sections of the assembled structures. Specifically, we designed a tetramer (xy-connectivity) and a pentamer (z-connectivity) where only specific monomers displayed anti-handles complementary to the DNA-Au NPs. Schematic illustrations of superstructure and anti-handle placement are shown in **Figure 3d**. Analysis of the assembled structures by TEM clearly shows that Au NPs only attached to those monomers displaying the anti-handle, while all other moDON monomers in the superstructure remained bare (see **Figures 3d**, and S33, S34). We thus confirmed that indeed, using the moDON, site-specific addressability of each specific monomer in the superstructure is fully retained and can be used for the selective modification with functional guest molecules such as NPs or proteins. This could enable further functionality for synthetic cell design allowing e.g. the selective attachment and transport of cargo along an artificial cytoskeleton^28^. Although not demonstrated here, the modular design of the moDON could also allow for several different modifications at pre-designed positions within the same superstructure.

In nature functional protein complexes or microtubules are trigger-dependently assembled and disassembled. Thus, after demonstrating controlled and trigger-specific assembly of superstructures as well as their retained addressability, we lastly investigated, if such superstructures could analogously be selectively disassembled, mimicking cellular systems. We initially tested disassembly for the z-connection. Due to the connector strand-mediated assembly of moDONs in the z-direction, removal of the connector by toehold-mediated strand displacement poses an excellent tool. For this, the connectors were extended to display a connector specific, short ssDNA toehold region of 7 nt. The addition of fully complementary invader strands could then selectively remove the connector strand (see NUPACK simulations in Figures S35 – 36, and Table S15) and thus result in superstructure disassembly as schematically depicted in **Figure 4a**. To illustrate the specificity of the connector-invader pairs, as well as their orthogonality towards the other pairs, we initially investigated the selective partial disassembly of a z-pentamer. Depending on the sequence of the invaders added, pentamers should selectively disassemble into various ensembles ranging from tetramers to monomers. Encouragingly, analysis by AGE and TEM showed that the programmed disassembly process is indeed highly selective and effective, disassembling the pentamers into the desired substructures of tetramers, trimers, dimers, and monomers depending on the invader strand added (**Figures 4b** and S37 – S40). To further showcase the effectiveness of this methodology, we next demonstrated the selective disassembly of periodic tubes made up from either monomeric or dimeric subunits in the z-direction. Once more, AGE revealed the highly effective and selective disassembly into monomeric (Figure S41a,b) or dimeric subunits (Figure S41c,d) as desired. Further examining the influence of different experimental conditions on z-disassembly, we observed that neither the amount of invader strand added (2-, 4-, or 8-fold excess over connector strands), nor the concentration of MgCl_2_ (in a range of 5 – 20 mM) had any significant effect on disassembly kinetics or yield with almost all structures being fully disassembled after ∼2 h if the correct invader was added (see **Figure 4d** and S42). This confirms that the main trigger for z-disassembly is the type of invader strand added, while the concentration of MgCl_2_ in the buffer played no significant role.

**Figure 4:**
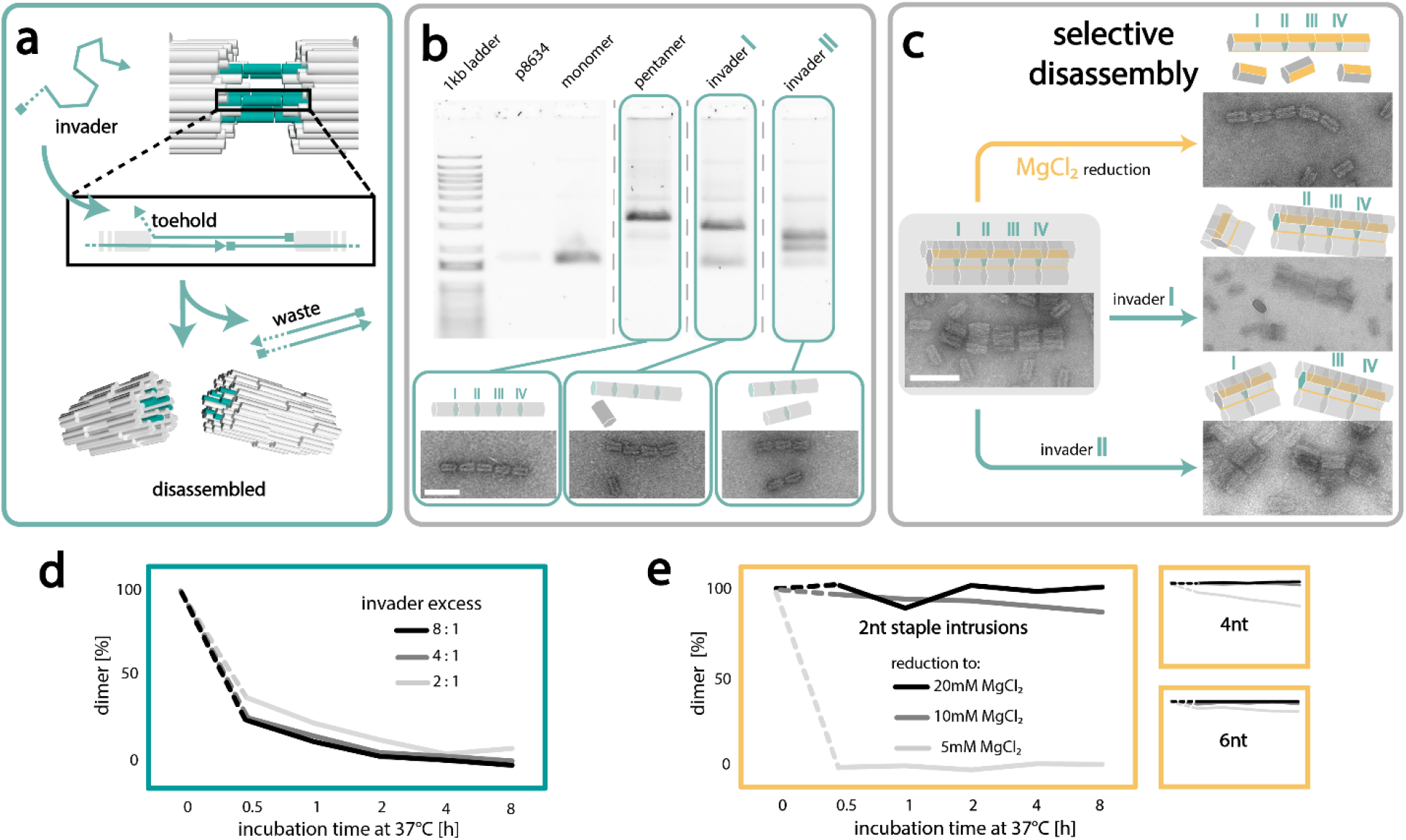
Disassembly of moDON superstructures. (**a**) Schematic of z-directional disassembly: Connector strands are elongated by a short ssDNA toehold. An invader strand, fully complementary to the elongated connector is able to hybridize first to the toehold, and then detach the connector from the ssDNA handles on the moDONs. This results in z-directional disassembly. (**b**) Selective disassembly of z-connections. The unique sequence of each connector allows for selective disassembly of single connections in the superstructure. Shown here is the invasion of connector I or connector II, by their respective invaders, resulting in disassembly into tetramers and monomers, or dimers and trimers respectively. (**c**) Selective disassembly of a tetrameric pentamer: Reduction of MgCl_2_ leads to disassembly of xy-connections, leaving only z-directional pentamer and monomers intact. Addition of invader strands I or II, leads to selective disassembly of connections I or II, leaving tetrameric monomers and tetramers or tetrameric dimers and trimers, respectively. (**d**) Disassembly yield over time at different levels of invader excess. (**e**) Dimer disassembly yield over time at different MgCl_2_ concentrations and staple intrusion lengths. xy-connections with only 2 nt staple intrusions dissociate fast, in low MgCl_2_ buffer. Longer staple intrusions of 4 nt or 6 nt stabilize dimers, proportional to the staple intrusion length. Scale bars are 100 nm.

While the three-strand connection system in the z-direction excellently lent itself for disassembly by toehold-mediated strand displacement, xy-assemblies could not easily be disassembled using the same method. We therefore investigated other parameters to achieve selective disassembly in the xy-direction. We hypothesized that if short lengths of the staple intrusions were utilized, disassembly could be mediated by reducing the MgCl_2_ concentration, as had been reported in the literature.^23^ To test this hypothesis, dimeric structures containing staple intrusions of 2, 4, or 6 nt were exposed to different MgCl_2_ concentrations for different reaction times. While for assembly kinetics, reaction time and MgCl_2_ concentration played an important role, as discussed earlier, for the disassembly kinetics the number of nt in the staple intrusions were found to be crucially important. As can be seen in **Figures 4e** (and S43), no dimer disassembly could be observed in 10 or 20 mM MgCl_2_ supplemented buffer for dimers formed with 4 and 6 nt staple intrusions. However, for the shortest system of only 2 nt staple intrusion length, a small number of monomers with higher electrophoretic mobility could be observed. Full disassembly of dimers into the respective monomers was only achieved for structures formed with 2 nt intrusions if the MgCl_2_ concentration in the buffer was reduced to 5 mM. As can be seen in **Figure 4e** (and S43), these dimers were fully disassembled after only 30 min, while all other dimers formed with longer staple intrusions did not fully disassemble even after 8 h under these conditions. This suggests that structural integrity of xy-connections is highly dependent on staple intrusion length, and MgCl_2_ concentration, which can be explained with the deeper free energy funnel from longer staple intrusions and decreased shielding of backbone repulsion from lower MgCl_2_ concentrations. Consequently, less stable xy-connections, with only 2 nt staple intrusions were used for selective disassembly.

We thus far showed that both xy- and z-directional assemblies can be selectively assembled and disassembled. We further showed that not only the triggers for inducing assembly (i.e. connector strand- and MgCl_2_-induced hybridization), but also the triggers of disassembly (i.e. addition of invader strands or reduction of MgCl_2_ concentration) are fully orthogonal. This should allow for selective disassembly of large superstructures into any desired subunit, breaking either xy- or z-connections. To test this, we designed a superstructure, consisting of five tetrameric subunits in the xy-direction, which further assembled into a pentamer of tetramers in the z-direction. The successful formation of the tetrameric pentamers was confirmed by TEM analysis (**Figures 4c** and S44). Subsequently, we exposed the assembled structure to the different disassembly triggers, *i*.*e*. low MgCl_2_ concentrations or invader strands in order to achieve either monomeric pentamers or tetramers of defined lengths (monomeric, dimeric etc.). Supporting our hypothesis, analysis by TEM revealed that depending on the trigger added, structures could selectively be disassembled into the desired subunits: When the MgCl_2_ concentration was reduced to 5 mM, only the xy-connections disassembled, resulting in a z-directional pentamer and a large number of monomers (**Figure 4c**, upper right panel and Figure S45). Conversely, addition of invader I resulted in tetrameric monomers and tetramers, (**Figure 4c**, middle right panel, and Figure S46), suggesting that only the first z-connection was released, as desired. Analogously, the addition of invader II resulted in disassembly of the second z-connection, keeping all other z-connections intact and thus the tetrameric pentamer was split into tetrameric dimers and trimers (**Figure 4c**, lower right panel, and Figure S47). With this we finally showed, that the moDON can be assembled and disassembled with complete xy-z orthogonality giving absolute and complete control over the process.

## MATERIALS AND METHODS

### DNA Origami Design, Synthesis and Purification

DNA origami structures were designed with caDNAno^5^ version 2.4.10 and simulated with oxDNA^29-32^ version 3.5.0 with 100 000 000 iterations at standard settings, after relaxation with 5000 CPU and 1 000 000 GPU iterations. DNA sequences for the three-strand-system were analyzed with NUPACK^33^ version 2.2.

Unmodified DNA oligos were purchased from IDT, desalted and dispersed in IDTE pH 8.0 at 100 µM. Scaffold DNA was produced as described previously^34^. DNA was stored at -20 °C until further use. Folding was performed on a Biometric TRIO thermocycler from Analytic Jena, or a MJ research PTC-200. The folding buffer consisted of 1× TAE (40 mM Tris, 20 mM acetic acid, 1 mM EDTA, pH 8.2) supplemented with 15 mM MgCl_2_. Scaffold and staples, in at least 5-fold excess, were added to the folding buffer,

denatured for 5 min at 65 °C, folded isothermally for 3 h at 50.5 °C and then held at 4 °C until further use. Purification of folded structures was performed with Amicon centrifugal filters (Merck Millipore, cat.no.: UFC5100) and TAE buffer with 5 mM MgCl_2_ in five washing steps at 8 000 rcf for 4 min. The purified structures were stored at -20 °C until further use.

### Superstructure Assembly

Connection of moDONs to superstructures were performed, if not further specified, by addition of MgCl_2_ to 40 mM for xy-connections or 5-fold excess of connectors over handles for z-connections to a moDON ensemble. This ensemble of moDONs was, if not further specified, equimolar with respect to the different monomers (usually 1 or 2 nM), in TAE (40 mM Tris, 20 mM acetic acid, 1 mM EDTA, pH 8.2) and incubated at 37 °C over-night.

### Agarose Gel Electrophoresis

AGE was performed, unless further specified, with 1 % agarose gels, TAE running buffer (40 mM Tris, 20 mM acetic acid, 1 mM EDTA, pH 8.2) supplemented with 11 mM MgCl_2_ and SYBR safe (invitrogen, cat.no.: S33102) at 70 V, for 90 min and on ice. Gels were imaged on a typhoon FLA 9000 laserscanner.

### TEM

Carbon-coated copper grids (Plano GmbH, cat.no.: S162-3) were treated for 30 s with oxygen plasma, 10 fmol of the respective sample was incubated on the grid for 5 min, the solution removed with filter paper, and subsequently stained with 5 µl of a 2 % uranylformate solution for 10 s, which was again removed with filter paper. The grid was air-dried before imaging. Imaging was performed on a Jeol-JEM-1230 transmission electron microscope, operating at an acceleration voltage of 80 kV. Micrographs were analyzed with Fiji^35^ version 2.9.0.

### Au NP Synthesis and moDON Modification

Au NP were synthesized as described previously^36^. In brief, 50 mL of 1 mM hydrogen tetrachloroaurate in milliQ water was heated in aqua-regia-cleaned glassware to a rolling simmer, stirring vigorously with a magnetic stir bar. Then 400 µl of a (also simmering) 2 % trisodium citrate solution was added. After a color change to dark red, the heat was turned off, and the solution stirred for an additional 15 min, and then put aside to cool to room temperature. The solution was then concentrated by centrifugation at 10 000 rcf for 10 min and re-dispersion of the pellet in milliQ water. The concentration was determined from absorption measurements taken with a Nanodrop 1000. Samples were stored at 4 °C until further use.

Au NPs were subsequently functionalized with thiolated DNA oligos (biomers.net, HPLC purified, resuspended in milliQ water). For this, thiolated DNA was added to the Au NPs in 12 500-fold excess and subsequently frozen at -20 °C. Functionalized Au NP were purified via four steps of centrifugation at 20 000 rcf for 10 min, subsequent supernatant removal and addition of the same amount of milliQ water. Concentrations were calculated from absorption measurements taken with a Nanodrop 1000. Samples were stored at 4 °C until further use.

moDON-Au NP constructs were formed by addition of moDON to DNA-coated-Au NPs while simultaneously vortexing at a low level, followed by 1 h of incubation at room temperature. DNA-Au NP conjugates were added with at least 5x excess over handles on the moDON. Constructs were optionally purified by AGE, with the above specified parameters.

## CONCLUSION

In summary, we introduced the moDON as a modular DNA origami structure that can assemble into diverse pre-designed superstructures in a controlled manner with fully retained site-specific addressability. Modularity is achieved through two distinct approaches: xy-directional assembly via modular scaffold strand routing and z-directional assembly via exchangeable connection handles in a three-strand system. With six positions in xy-direction, each being able to form one of two connection sites (α-ζ) or be passive, and two opposite positions in z-direction being able to form one of four connection sites or be passive, the total number of unique monomers that can be formed through the modularity of just one single moDON is:

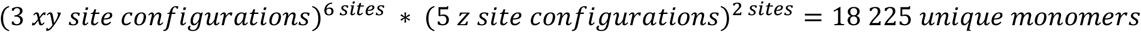

With 18,225 unique monomers possible from a single moDON set of staples, our method significantly reduces cost (total number of stapled needed are only ∼ 1,9× higher than those generally needed for a single structure (Table S18)), and design time compared to individual design of each DNA origami. To change the configuration of one connection site only ∼ 2.5 % of the staples need to be exchanged.

We demonstrated the versatility of our approach by assembling and disassembling various superstructures reaching µm and GDa scales, while retaining selective addressability of individual monomers. Notably, our approach allows assembly and disassembly on biologically relevant length- and timescales, opening possibilities for creating intricate artificial systems, including synthetic cell cytoskeletons. Compared to DNA nanotubes, mostly used to mimic the natural cytoskeleton in synthetic biology, the here constructed periodic tubes have a very large subunit, predisposing them for the incorporation of more complex approaches to multifunctionality.

The here presented methods of modularity will prove a powerful additional in the toolbox of DNA origami construction. We expect the xy-z orthogonal modularity and connections design to be adaptable to any other DNA origami structure, not only the moDON, potentially even allowing for further expansions of orthogonal connection sites. All in all, these features present the modular approach to DON superstructure construction as a promising new tool for the construction of large superstructures with exceptional control, suitable for synthetic biology applications.

## AUTHOR INFORMATION

### Author Contributions

The manuscript was written through contributions of both authors. The authors have given approval to the final version of the manuscript.

### Notes

The authors declare no competing financial interests.

## ACKNOWLEDGMENT

The authors are thankful for support with TEM imaging from M. Braun and U. Weber.

## REFERENCES

1 Murrell, M., Oakes, P. W., Lenz, M. & Gardel, M. L. Forcing cells into shape: the mechanics of actomyosin contractility. Nat Rev Mol Cell Bio 16, 486–498, doi:10.1038/nrm4012 (2015).

2 Brouhard, G. J. & Rice, L. M. Microtubule dynamics: an interplay of biochemistry and mechanics. Nat Rev Mol Cell Bio 19, 451–463, doi:10.1038/s41580-018-0009-y (2018).

3 Milo, R. & Phillips, R. Cell Biology by the Numbers. (Taylor & Francis, 2015).

4 Rothemund, P. W. Folding DNA to create nanoscale shapes and patterns. Nature 440, 297–302, doi:10.1038/nature04586 (2006).

5 Douglas, S. M. et al. Rapid prototyping of 3D DNA-origami shapes with caDNAno. Nucleic Acids Res 37, 5001–5006, doi:10.1093/nar/gkp436 (2009).

6 Pumm, A. K. et al. A DNA origami rotary ratchet motor. Nature 607, 492–498, doi:10.1038/s41586-022-04910-y (2022).

7 Raab, M. et al. Using DNA origami nanorulers as traceable distance measurement standards and nanoscopic benchmark structures. Sci Rep 8, 1780, doi:10.1038/s41598-018-19905-x (2018).

8 Kuzyk, A. et al. DNA-based self-assembly of chiral plasmonic nanostructures with tailored optical response. Nature 483, 311–314, doi:10.1038/nature10889 (2012).

9 Acuna, G. P. et al. Fluorescence enhancement at docking sites of DNA-directed self-assembled nanoantennas. Science 338, 506–510, doi:10.1126/science.1228638 (2012).

10 Nickels, P. C. et al. Molecular force spectroscopy with a DNA origami-based nanoscopic force clamp. Science 354, 305–307, doi:10.1126/science.aah5974 (2016).

11 Funke, J. J. et al. Uncovering the forces between nucleosomes using DNA origami. Sci Adv 2, e1600974, doi:10.1126/sciadv.1600974 (2016).

12 Berger, R. M. L. et al. Nanoscale FasL Organization on DNA Origami to Decipher Apoptosis Signal Activation in Cells. Small 17, e2101678, doi:10.1002/smll.202101678 (2021).

13 Hellmeier, J. et al. DNA origami demonstrate the unique stimulatory power of single pMHCs as T cell antigens. P Natl Acad Sci USA 118, doi:10.1073/pnas.2016857118 (2021).

14 Veneziano, R. et al. Role of nanoscale antigen organization on B-cell activation probed using DNA origami. Nat Nanotechnol 15, 716-+, doi:10.1038/s41565-020-0719-0 (2020).

15 Marchi, A. N., Saaem, I., Vogen, B. N., Brown, S. & LaBean, T. H. Toward larger DNA origami. Nano Lett 14, 5740–5747, doi:10.1021/nl502626s (2014).

16 Engelhardt, F. A. S. et al. Custom-Size, Functional, and Durable DNA Origami with Design-Specific Scaffolds. Acs Nano 13, 5015–5027, doi:10.1021/acsnano.9b01025 (2019).

17 Wei, B., Dai, M. J. & Yin, P. Complex shapes self-assembled from single-stranded DNA tiles. Nature 485, 623-+, doi:10.1038/nature11075 (2012).

18 Ke, Y., Ong, L. L., Shih, W. M. & Yin, P. Three-dimensional structures self-assembled from DNA bricks. Science 338, 1177–1183, doi:10.1126/science.1227268 (2012).

19 Ong, L. L. et al. Programmable self-assembly of three-dimensional nanostructures from 10,000 unique components. Nature 552, 72-+, doi:10.1038/nature24648 (2017).

20 Yao, G. et al. Meta-DNA structures. Nat Chem 12, 1067–1075, doi:10.1038/s41557-020-0539-8 (2020).

21 Wintersinger, C. M. et al. Multi-micron crisscross structures grown from DNA-origami slats. Nat Nanotechnol 18, 281–289, doi:10.1038/s41565-022-01283-1 (2023).

22 Wagenbauer, K. F., Sigl, C. & Dietz, H. Gigadalton-scale shape-programmable DNA assemblies. Nature 552, 78–83, doi:10.1038/nature24651 (2017).

23 Gerling, T., Wagenbauer, K. F., Neuner, A. M. & Dietz, H. Dynamic DNA devices and assemblies formed by shape-complementary, non-base pairing 3D components. Science 347, 1446–1452, doi:10.1126/science.aaa5372 (2015).

24 Tigges, T., Heuser, T., Tiwari, R. & Walther, A. 3D DNA Origami Cuboids as Monodisperse Patchy Nanoparticles for Switchable Hierarchical Self-Assembly. Nano Letters 16, 7870–7874, doi:10.1021/acs.nanolett.6b04146 (2016).

25 Tikhomirov, G., Petersen, P. & Qian, L. L. Fractal assembly of micrometre-scale DNA origami arrays with arbitrary patterns. Nature 552, 67–71, doi:10.1038/nature24655 (2017).

26 Ke, Y. et al. Multilayer DNA origami packed on a square lattice. J Am Chem Soc 131, 15903–15908, doi:10.1021/ja906381y (2009).

27 Wagenbauer, K. F. et al. How We Make DNA Origami. Chembiochem 18, 1873–1885, doi:10.1002/cbic.201700377 (2017).

28 Zhan, P., Jahnke, K., Liu, N. & Gopfrich, K. Functional DNA-based cytoskeletons for synthetic cells. Nat Chem 14, 958–963, doi:10.1038/s41557-022-00945-w (2022).

29 Snodin, B. E. et al. Introducing improved structural properties and salt dependence into a coarse-grained model of DNA. J Chem Phys 142, 234901, doi:10.1063/1.4921957 (2015).

30 Rovigatti, L., Sulc, P., Reguly, I. Z. & Romano, F. A comparison between parallelization approaches in molecular dynamics simulations on GPUs. J Comput Chem 36, 1–8, doi:10.1002/jcc.23763 (2015).

31 Sulc, P. et al. Sequence-dependent thermodynamics of a coarse-grained DNA model. J Chem Phys 137, 135101, doi:10.1063/1.4754132 (2012).

32 Ouldridge, T. E., Louis, A. A. & Doye, J. P. Structural, mechanical, and thermodynamic properties of a coarse-grained DNA model. J Chem Phys 134, 085101, doi:10.1063/1.3552946 (2011).

33 Zadeh, J. N. et al. NUPACK: Analysis and design of nucleic acid systems. J Comput Chem 32, 170–173, doi:10.1002/jcc.21596 (2011).

34 Russell, D. W. S. J. Molecular Cloning: A Laboratory Manual. Vol. 1 (Cold Spring Harbor Laboratory, Cold Spring Harbor: New York, NY, USA, 2001).

35 Schindelin, J. et al. Fiji: an open-source platform for biological-image analysis. Nat Methods 9, 676–682, doi:10.1038/nmeth.2019 (2012).

36 Enustun, B. V. & Turkevich, J. Coagulation of Colloidal Gold. J Am Chem Soc 85, 3317-+, doi:DOI 10.1021/ja00904a001 (1963).

