## Supplementary Information for "Fully addressable, designer superstructures assembled from a single modular DNA origami"

### Table of Figures

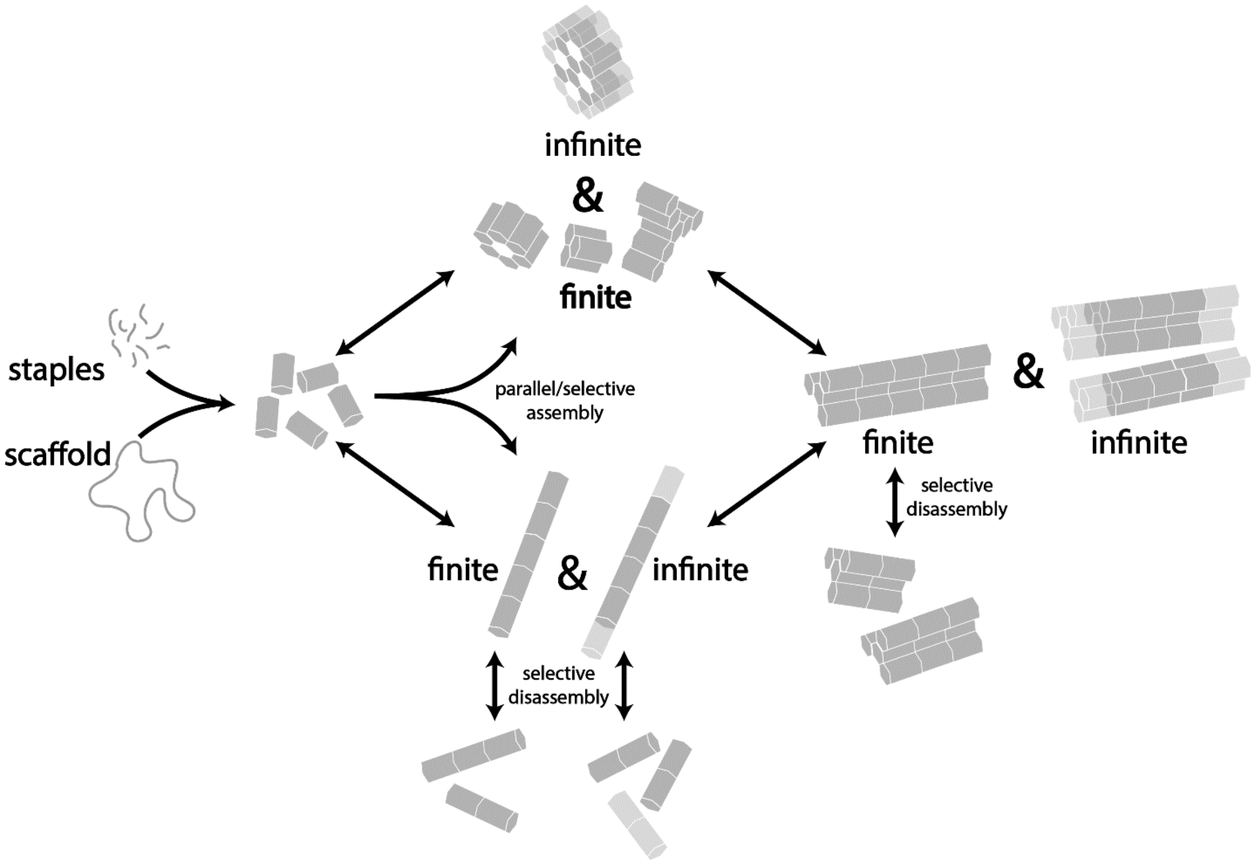

**Figure S1: connectivity overview**: different moDON monomers are folded separately. Specific superstructures are designed by a combination of moDONs with complimentary connection sites. In the xy-direction, perpendicular to the helical direction, finite and infinite structures are assembled by increasing the $MgCl_2$ concentration. In the z-direction, the helical direction, finite and infinite structures are formed by addition of connectors strands. Connections are orthogonal to each other, as well as to the connection strategies themselves. This enables *parallel* assembly of different xy-directional and z-directional structures in one reaction vessel. This also enables *selective* assembly of either xy- or z-structures, from moDONs carrying both connection sites for xy- and z- structures, depending on which trigger is added. Both approaches can be combined to form large structures in all directions. Conversely, a decrease in $MgCl_2$ concentration leads to disassembly of xy-connections. The addition of invader strands leads to selective disassembly of z-connections through toehold-mediated strand displacement. Both disassembly strategies are also orthogonal towards each other. In case of z-disassembly, the sequence specificity of the connectors and invaders also leads to orthogonality of the single connection sites towards each other.

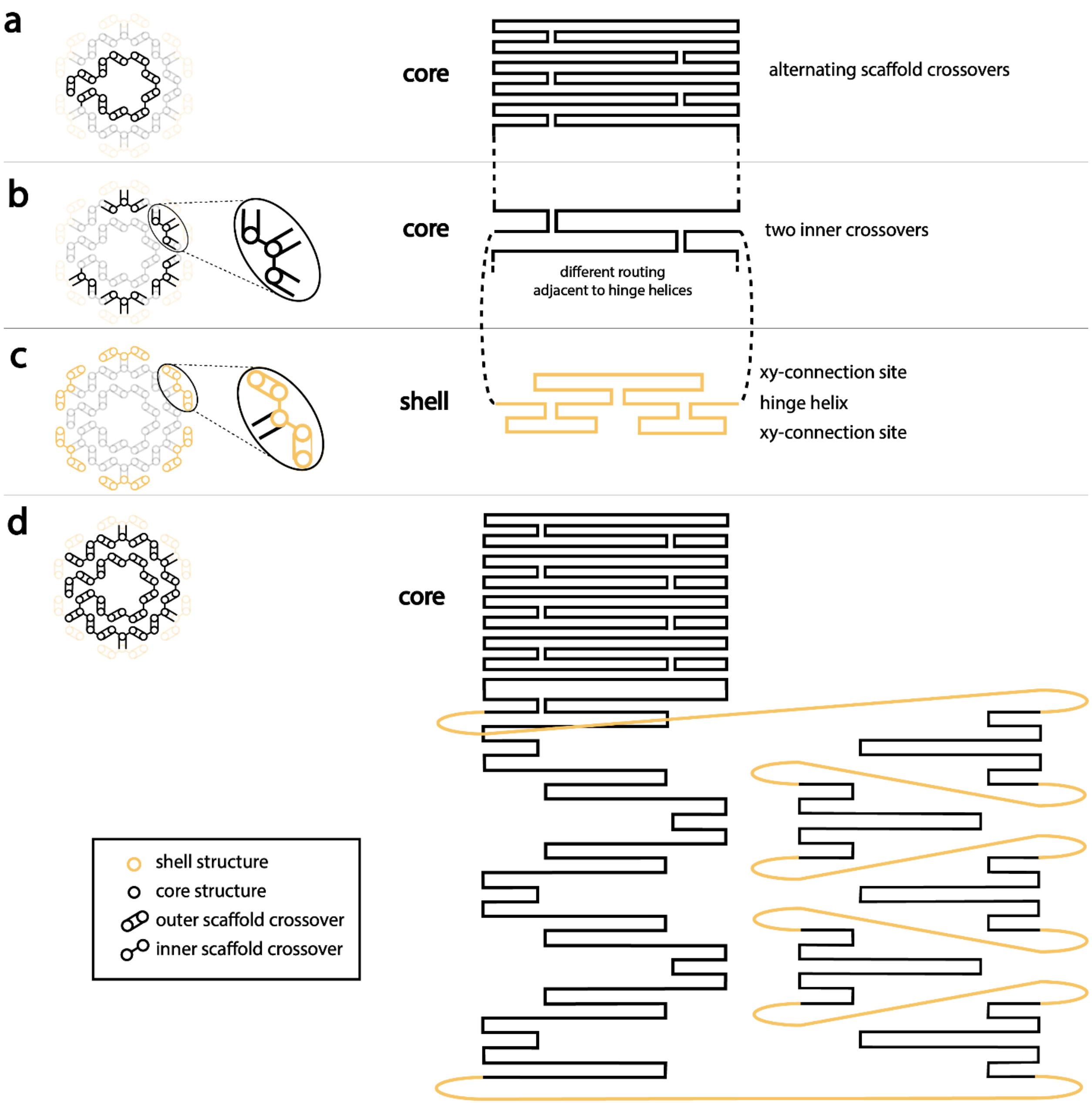

**Figure S2: Scaffold routing in core and shell:** (**a**) the inner core part (black) of the moDON is layered evenly, with alternating scaffold cross-overs at the helix ends and in the helix middle. This was done to ease the folding pathway. The positions of the scaffold cross-overs in the helix middle are varied, to avoid the introduction of an artificial breaking point. (**b**) The outer core part (black) of the moDON was not able to be layered with alternating scaffold cross-over positions. Every fifth helix has three cross-overs, to connect with a hinge helix (yellow) (**c**). From the hinge helix the xy-connection sites (yellow) loop out. The connection sites are modular and their routing is changed by exchange of a few staples. The hinge helix connects with scaffold crossovers at the helix ends back to the core structure. The connections from this adjacent helix to the second and third helix is then done with scaffold cross-overs in the middle of the helix. This leads to the scaffold routing pattern shown in (**d**), with the connection sites shown as curved yellow lines.

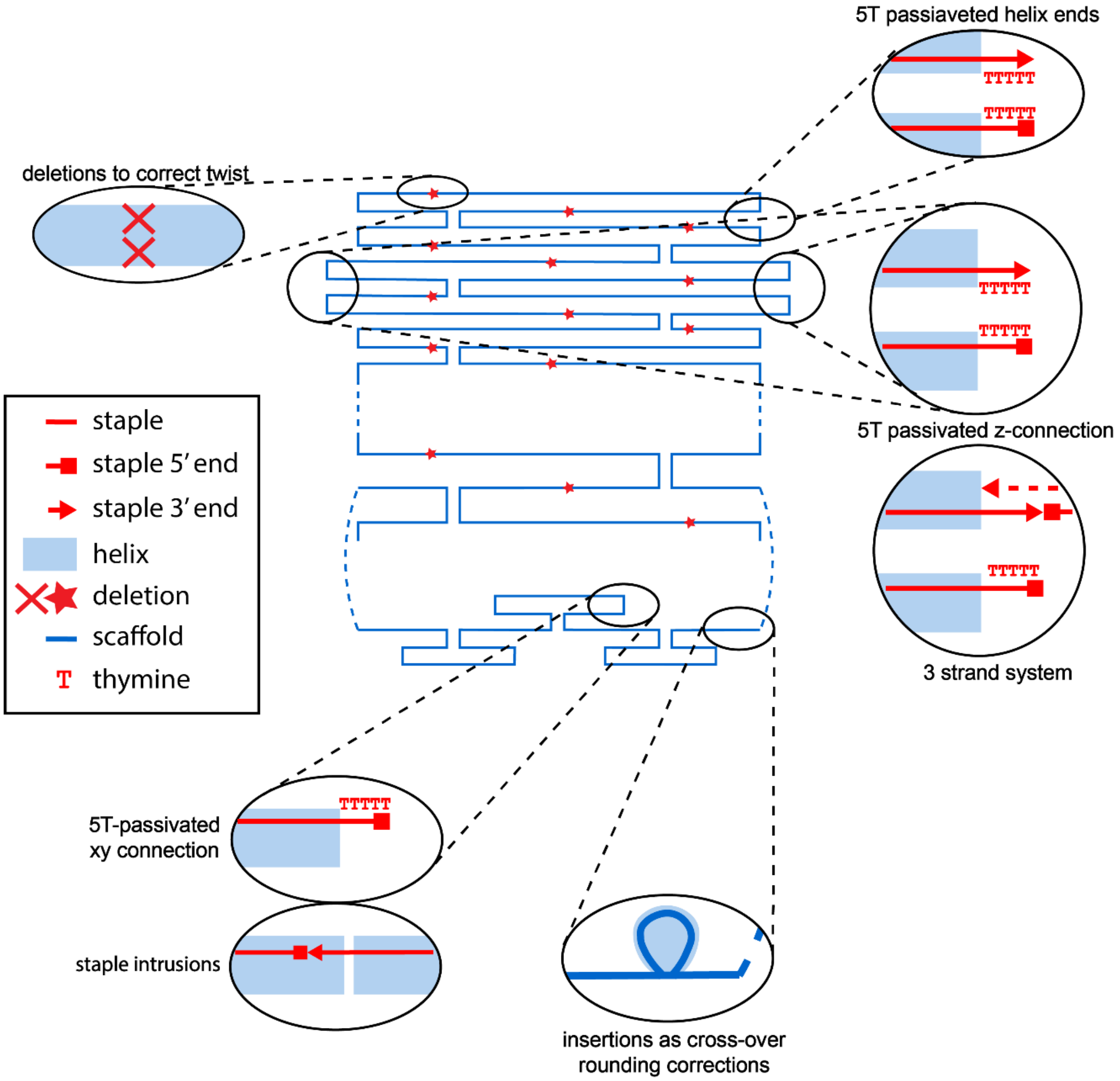

**Figure S3: Staple routing and positioning of corrections:** the residual twist of the honeycomb lattice was corrected with deletions every 126 bp and the crossover rounding errors were corrected with insertions at the connection site ends (see also Figure S5b), to keep the number of bp in the modular shell even. To prevent unwanted blunt-end interactions of the moDON, the helical ends were passivated with 5 Thymine (T) bases. The same strategy was used to passivate both connections sites of xy- and z-connections if desired. Connections in xy-direction were stabilized by staple intrusions, of elongated staples from one moDON to short staple omission in the complementary structure. Connections in z-direction were constructed with a three-strand-system. Here the 5' ends on the moDONs left side and the 3' ends on the moDONs right side were elongated by 10 nt or 11 nt, respectively, both constituting handles, complementary to each a half of a connector strand. Assigning 5' elongations to the left and 3' elongations to the right side of the moDON, achieved directionality of the z-connection. For staple sequences see Tables S2-S15.

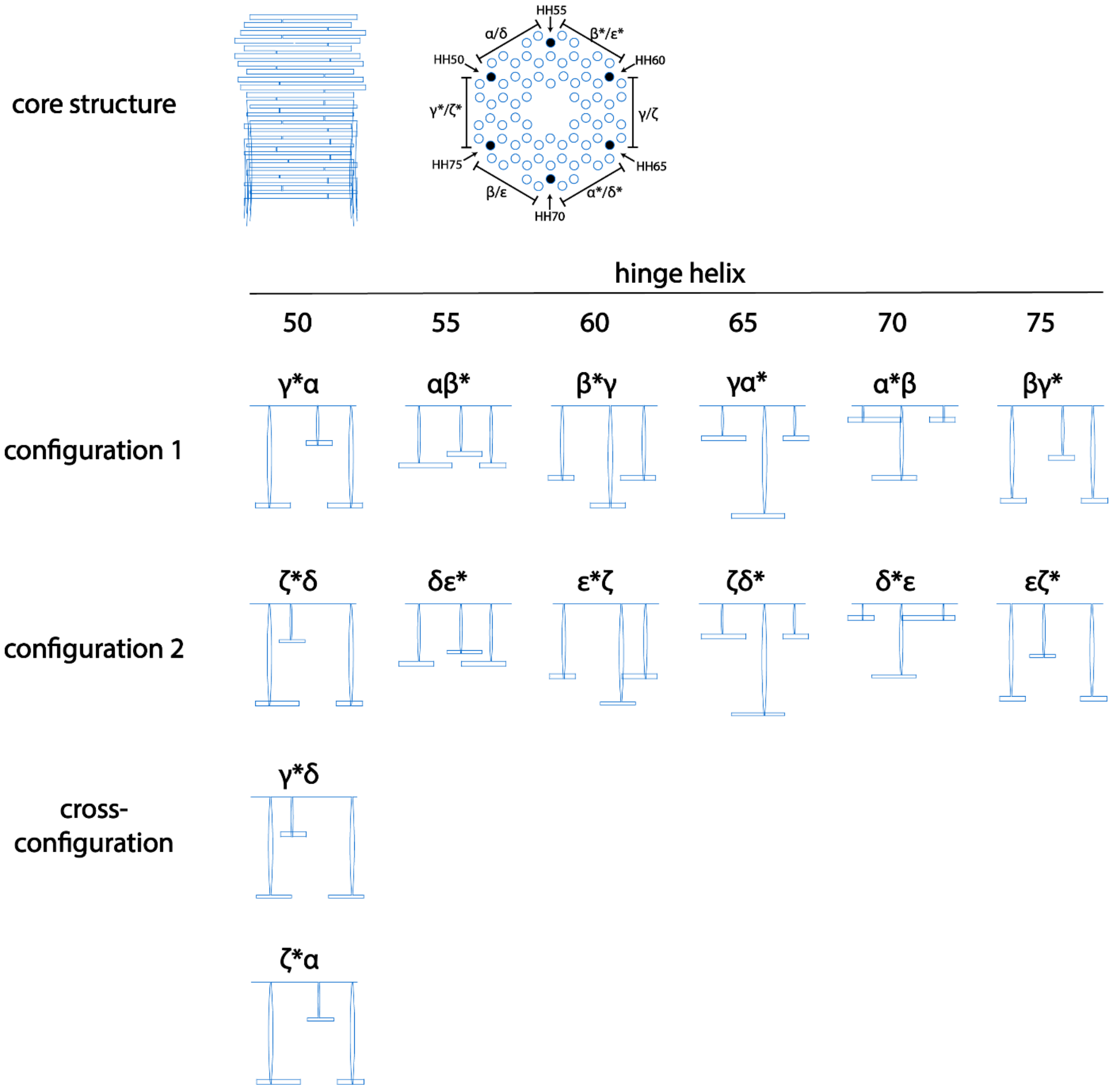

**Figure S4: caDNAno scaffold routing** paths of the different xy-connection sites. The core routing (top left image) always remains the same, while the shell is modular. All different configurations are arranged with respect to their configuration, and their hinge helix (HH). Note here, that HH50 has four possible configurations, while HH65 only has one. All other HH have two configurations. Also compare to Table S1. For staple sequences see Tables S2-S15.

a

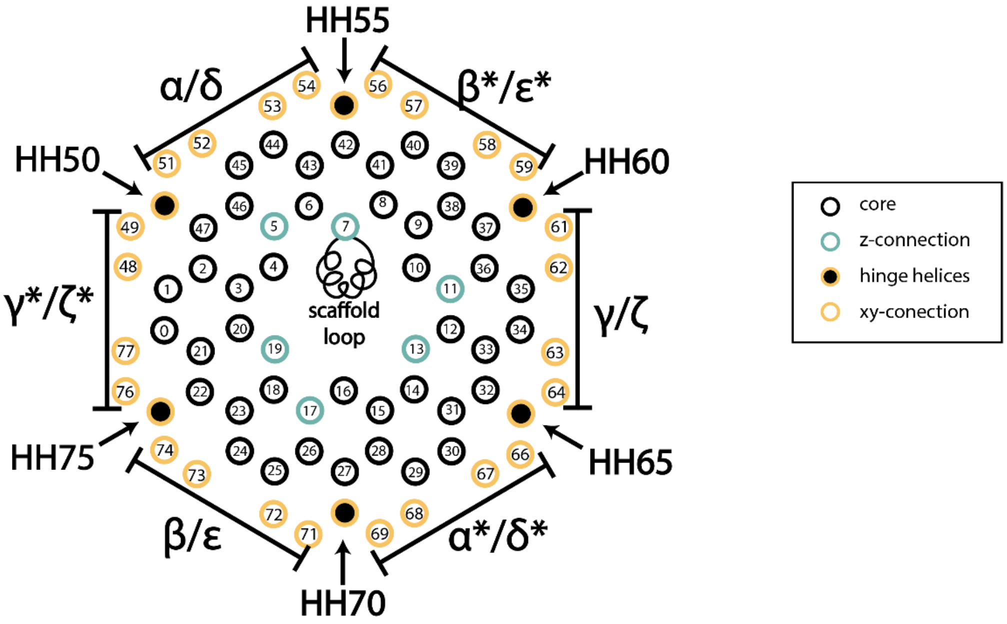

b

| hinge helix # | number of nt in scaffold loop αβγ | δεζ | γ*δ | ζ*α | final |
|---|---|---|---|---|---|
| 50 | 358 +2 | 356 +4 | 356 +4 | 358 +2 | 360 |
| 55 | 400 | 400 | 400 | 400 | 400 |
| 60 | 356 | 356 | 356 | 356 | 356 |
| 65 | 420 | 420 | 420 | 420 | 420 |
| 70 | 420 +2 | 422 | 422 | 422 | 422 |
| 75 | 314 | 314 | 314 | 314 | 314 |

**Figure S5: Helix numeration and scaffold loop corrections**: (**a**) Helix numeration of the moDON. Helices in yellow are part of the xy-connection sites, helices in turquois are z-connections sites, black helices are purely structural. Hinge helices are yellow with a black core. The scaffold loop was placed in the middle of helix 7, facing inwards into the moDON. (**b**) Since the literature value for one full helical turn in B-DNA configuration is not an integer, it is either rounded up or down in the caDNAno software to 11 or 10 bp. Changing the position and/or length of the modular parts from one configuration to the other leads to different amounts of nt in the scaffold of the modular parts. Those were corrected by small insertions. For staple sequences see Tables S2-S15.

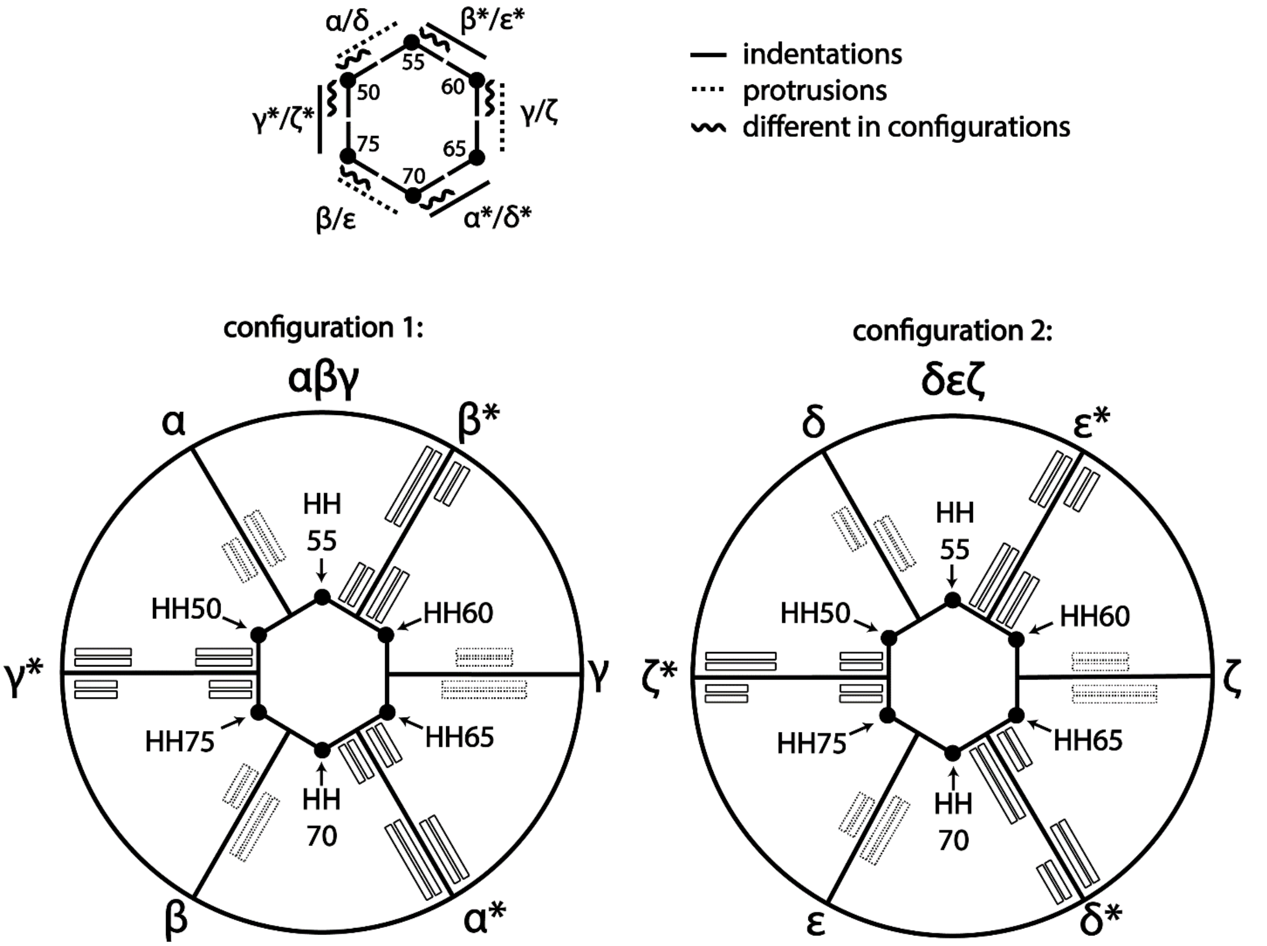

**Figure S6: Overview over connection sites and configurations**: All 12 connection sites are orthogonal to each other, across both configurations. Protrusions fit accurately into indentations, which are always denoted with the same letter and an asterisk. The position of connection sites cannot be changed in the moDON, but the connection site can be either from configuration 1 or configuration 2. Each connection site can be passivated. For staple sequences see Tables S2-S15.

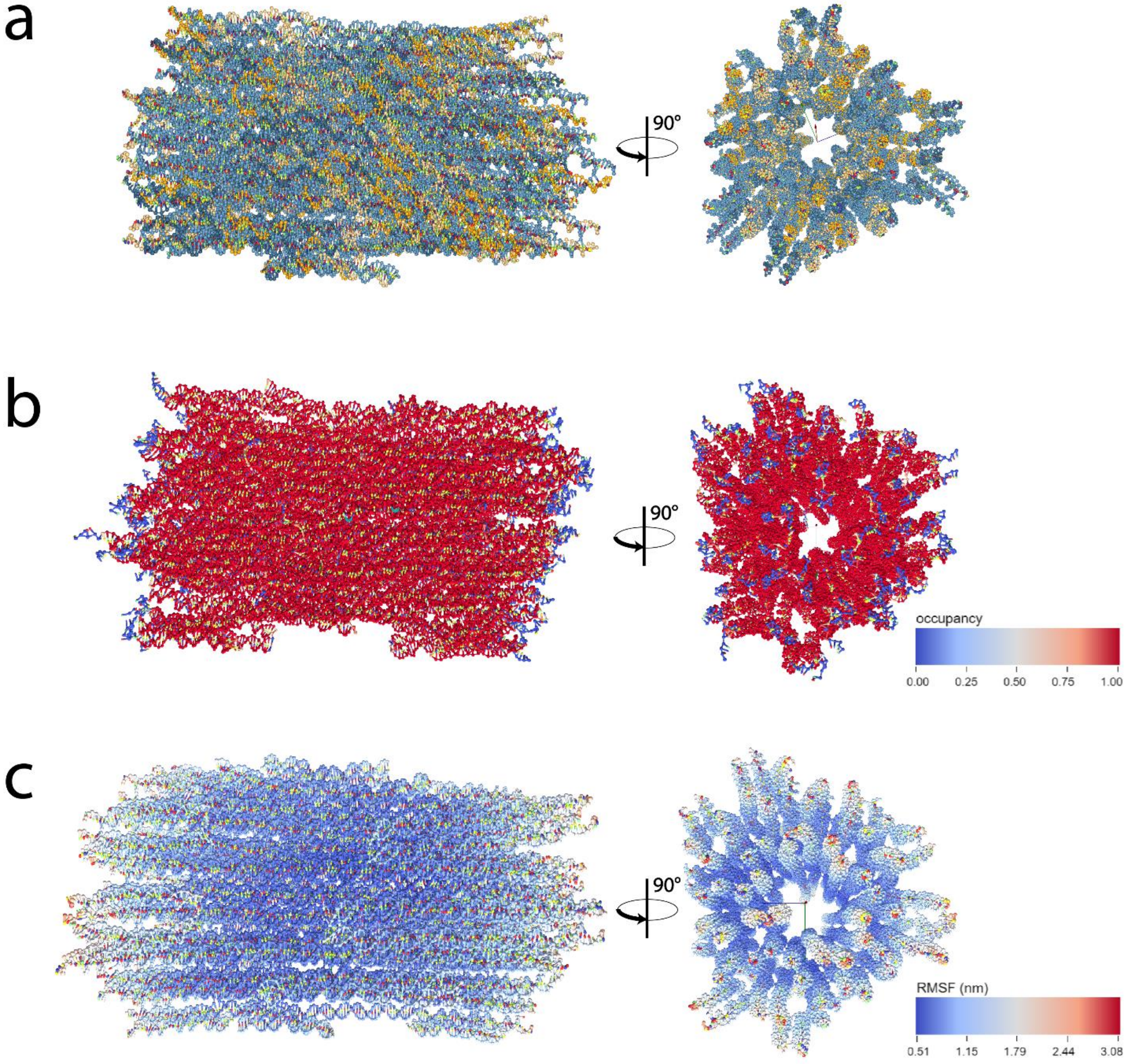

**Figure S7: oxDNA simulation of the moDON in configuration 1 (αβγ):** Side view to the left and top view on the right of (**a**) the averaged structure (**b**) the bond occupancy, indicating a high occupancy of the whole structure, except the 5T passivated helix ends (**c**) root mean squared fluctuation (RMSF) on the average structure, showing low flexibility across the sides, and marginally more towards the helix ends. Higher fluctuation is only seen for the 5T ssDNA passivation, explained by the lower persistence length of ssDNA compared to dsDNA. This oxDNA simulation, compared to the oxDNA simulation of the moDON in configuration 1 (Figure S8) suggests overall structural stability, even with changed shell structure.. The relaxation was performed with 5 000 CPU, and 1 000 000 GPU iterations, the simulation was performed with 100 000 000 iterations, all at 20°C, 1 M NaCl.

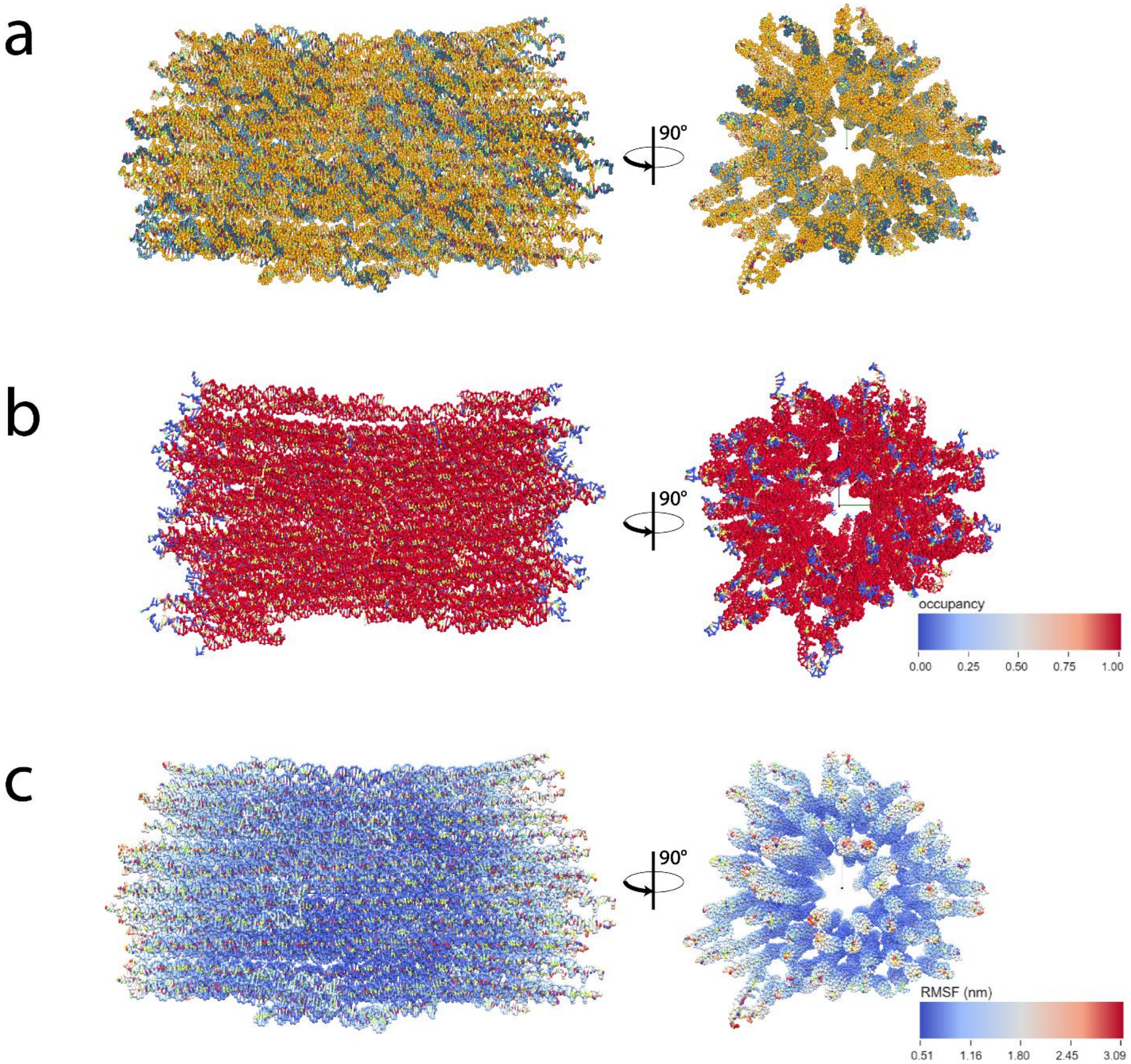

**Figure S8: oxDNA simulation of the moDON in configuration 2 (δεζ):** Side view to the left and top view on the right of (**a**) the averaged structure (**b**) the bond occupancy, indicating a high occupancy of the whole structure, except the 5T passivated helix ends (**c**) root mean squared fluctuation (RMSF) on the average structure, showing low flexibility across the sides, and marginally more towards the helix ends. Higher fluctuation is only seen for the 5T ssDNA passivation, explained by the lower persistence length of ssDNA compared to dsDNA. This oxDNA simulation, compared to the oxDNA simulation of the moDON in configuration 1 (Figure S7) suggests overall structural stability, even with changed shell structure. The relaxation was performed with 5 000 CPU, and 1 000 000 GPU iterations, the simulation was performed with 100 000 000 iterations, all at 20°C, 1 M NaCl.

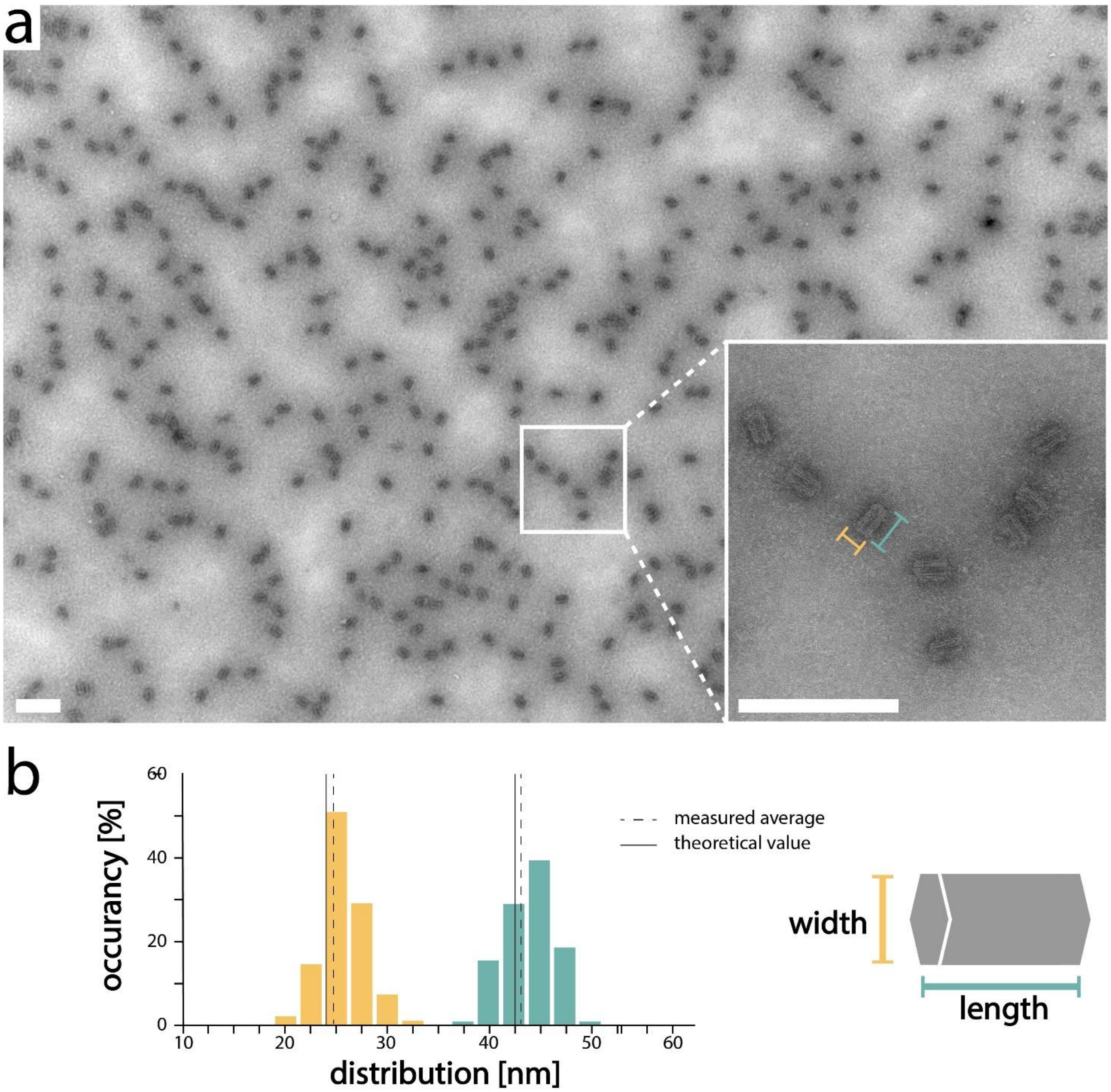

**Figure S9: Analysis of moDON monomers** (**a**) TEM micrographs of properly folded moDON monomers. (**b**) Size distribution of length (turquois) and width (yellow). Measured averages (length 42.98 nm, width 24.67 nm) is slightly larger than expected theoretical values (length 42.5 nm and width 24.0 nm) as calculated from number of bp in each helix of the core structure (125 bp) and number of helices at broadest point (12 helices). N > 100 individual monomers. Scale bars are 200 nm.

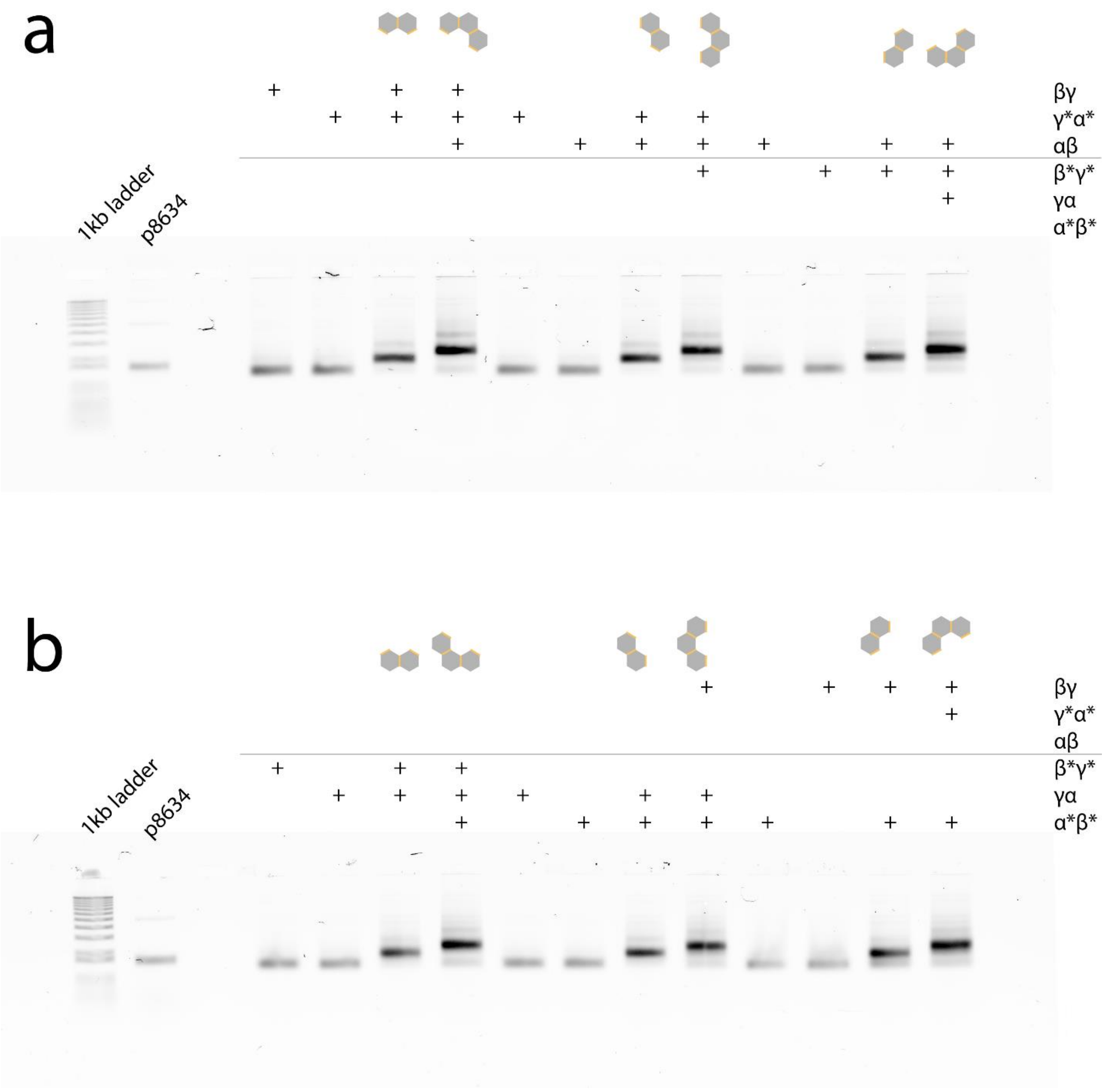

**Figure S10: AGE gel shift assay of dimer/trimer permutations 1:** moDONs in configuration 1. (**a**) and (**b**) show all permutations of dimers and trimers as constructed from the moDONs βγ, γ*α*, αβ, β*γ*, γα, and α*β*, as well as the respective monomers.

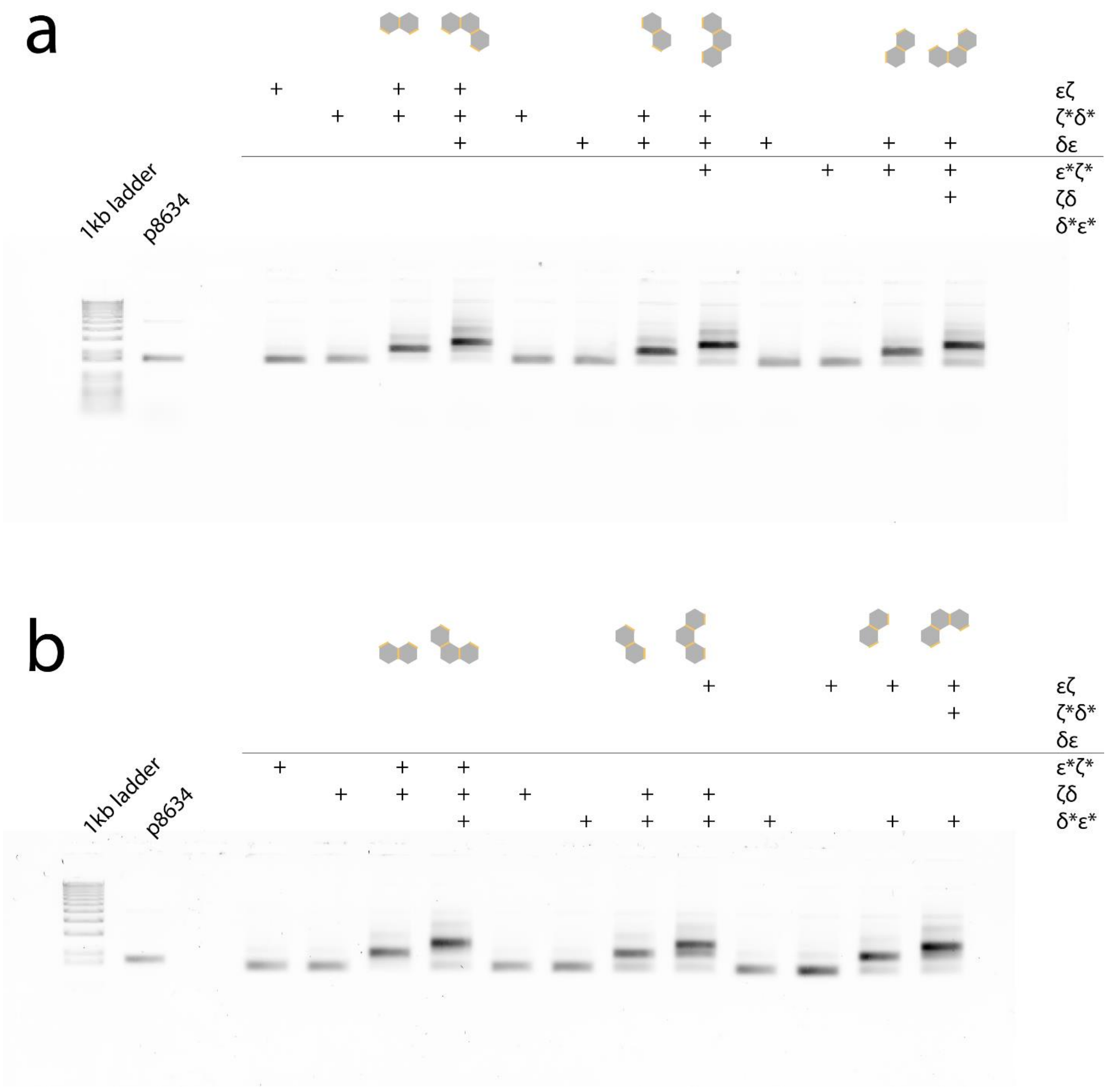

**Figure S11: AGE gel shift assay of dimer/trimer permutations 2:** moDONs in configuration 2. (**a**) and (**b**) show all permutations of dimers and trimers as constructed from the moDONs εζ, ζ*δ, δε, ε*ζ*, ζδ, and δ*ε*, as well as the respective monomers.

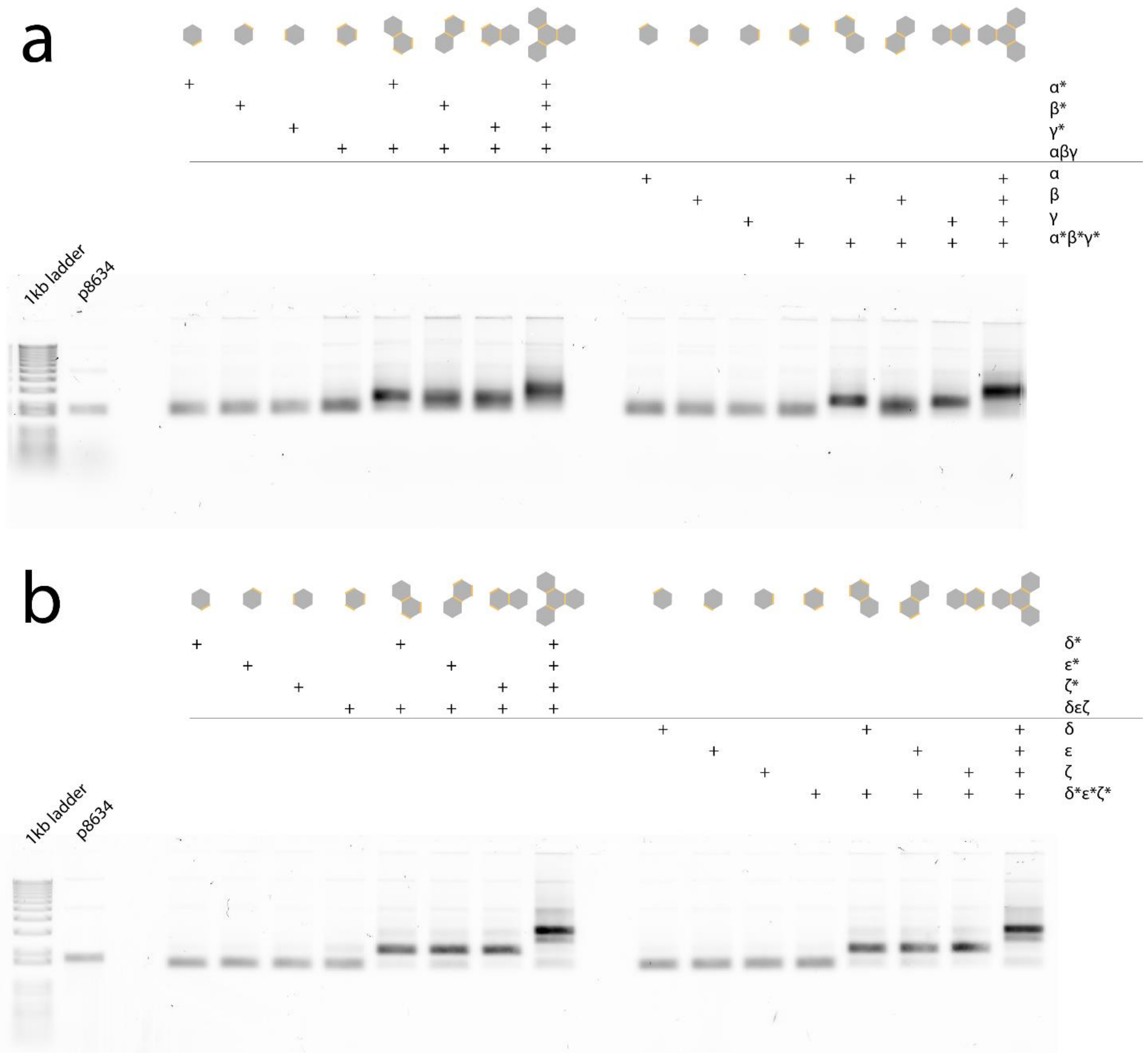

**Figure S12: AGE gel shift assay of tetramers** of moDONs in configurations 1 and 2. All monomers and dimers of the sub-structures, as well as the final tetramers are shown. (**a**) In configuration 1 α*, β*, γ*, and αβγ for the first tetramer and α, β, γ, and α*β*γ* for the second tetramer were used. (**b**) In configuration 2 δ*, ε*, ζ* and δεζ were used for the first tetramer and δ, ε, ζ, as well as δεζ were used for the second tetramer.

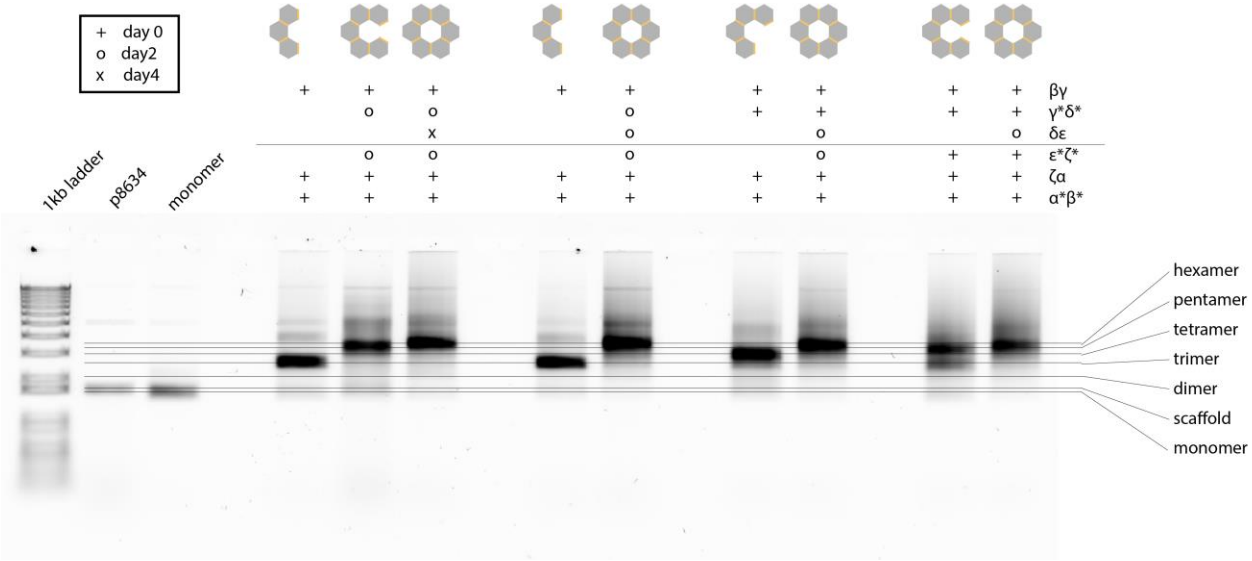

**Figure S13: AGE gel shift assay of hexamers** made from moDONs with connection sites from both configurations. Here, the moDONs α*β* and βγ only have connection sites from configuration 1; δε, and ε*ζ* only have connection sites from configuration 2. The connection sites of moDON monomers γ*δ* as well as ζα are a mixture from both configurations. moDONs were added subsequently to circumvent the formation of equimolar parts of mutually exclusive tetramers or pentamers, not able to form hexamers.

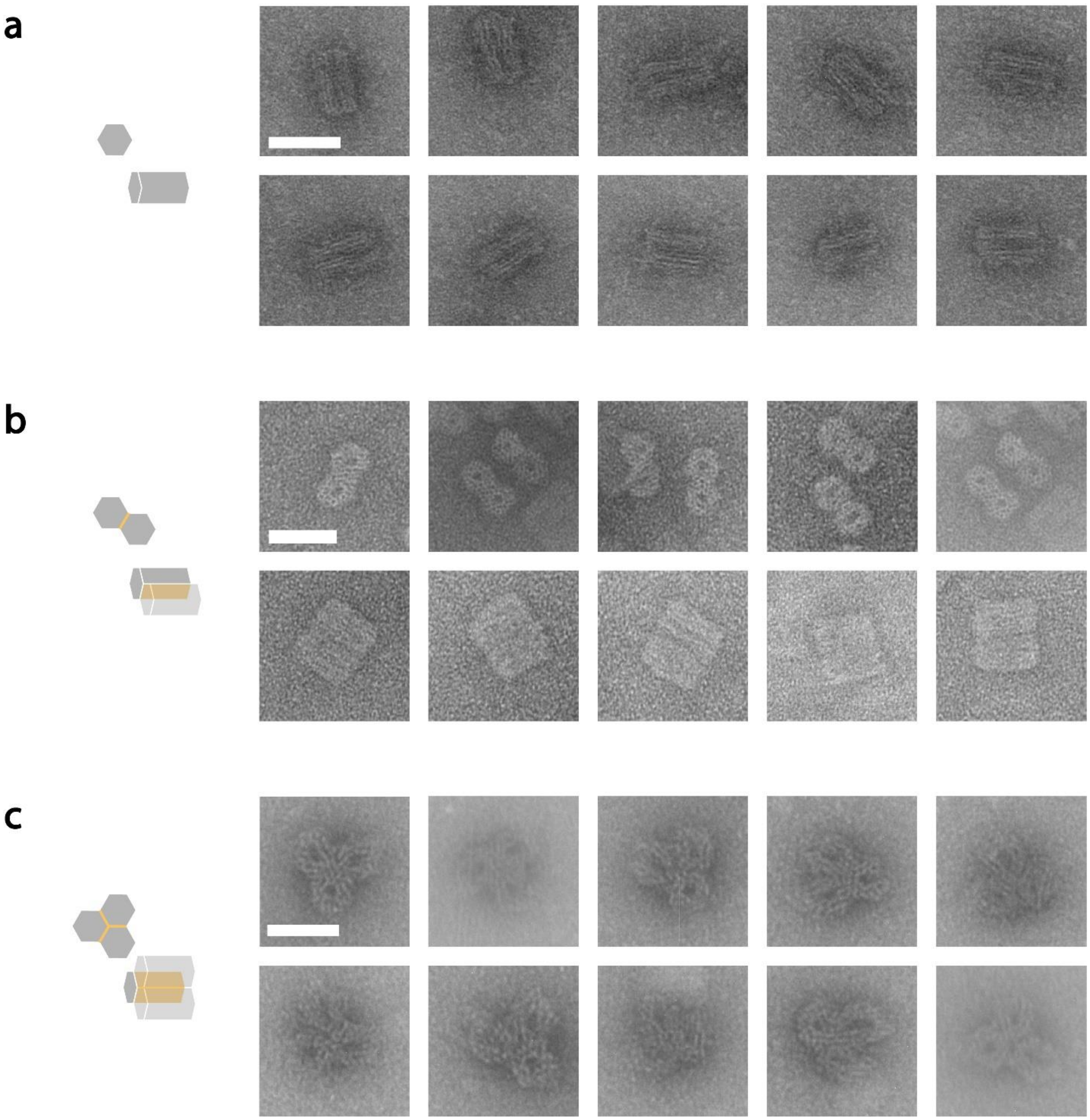

**Figure S14: Gallery of xy-structures 1:** Showing TEM micrographs of (**a**) monomers, (**b**) dimers, and (**c**) trimers. Scale bars are 50 nm and hold for all micrographs of the respective structure.

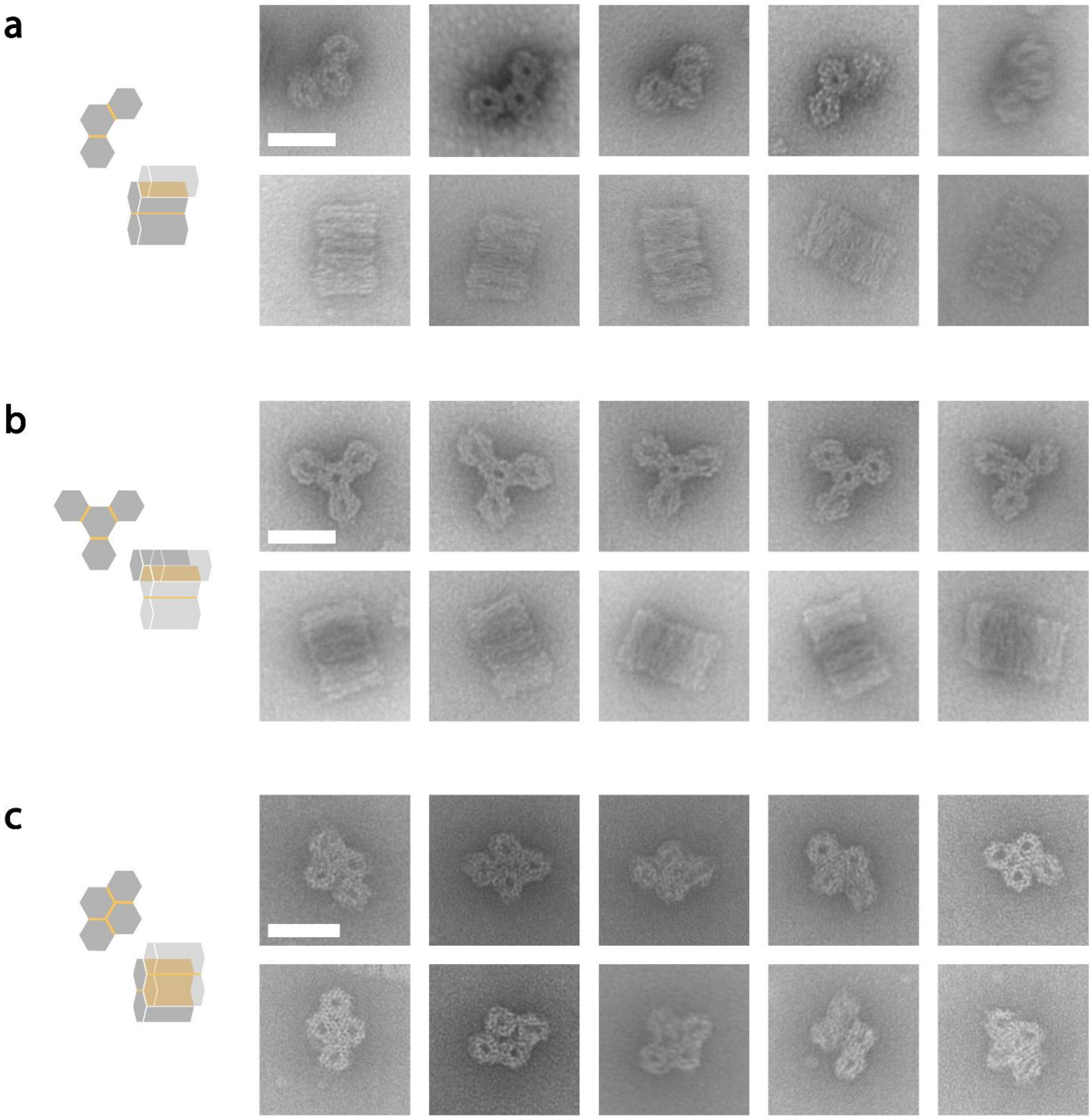

**Figure S15: Gallery of xy-structures 2**: Showing TEM micrographs of (**a**) trimers, (**b**), (**c**) different tetramers. Scale bars are 50 nm and hold for all micrographs of the respective structure.

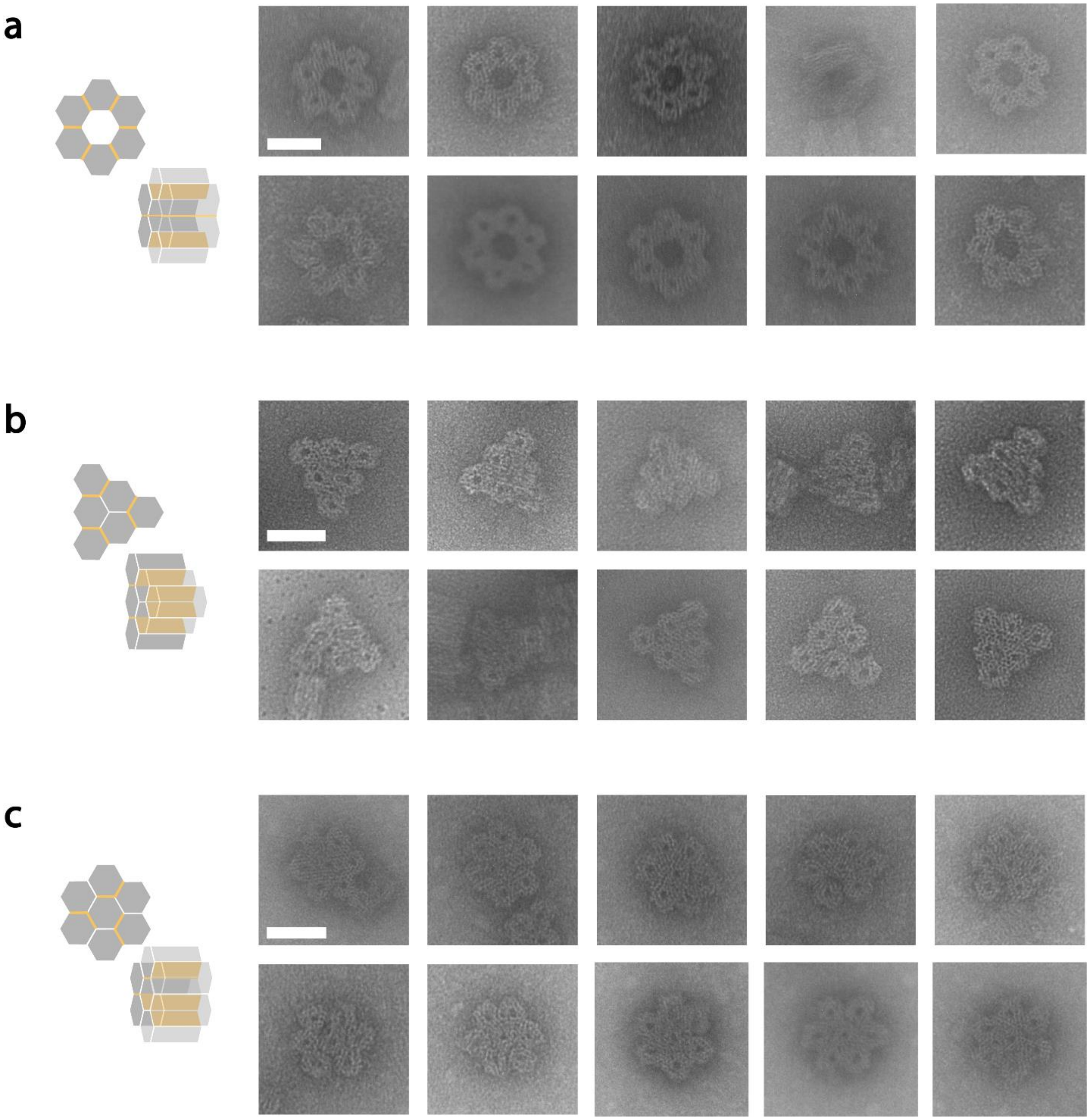

**Figure S16: Gallery of xy-structures 3:** Showing TEM micrographs of different (**a**), (**b**) hexamers, and (**c**) heptamers. Scale bars are 50 nm and hold for all micrographs of the respective structure.

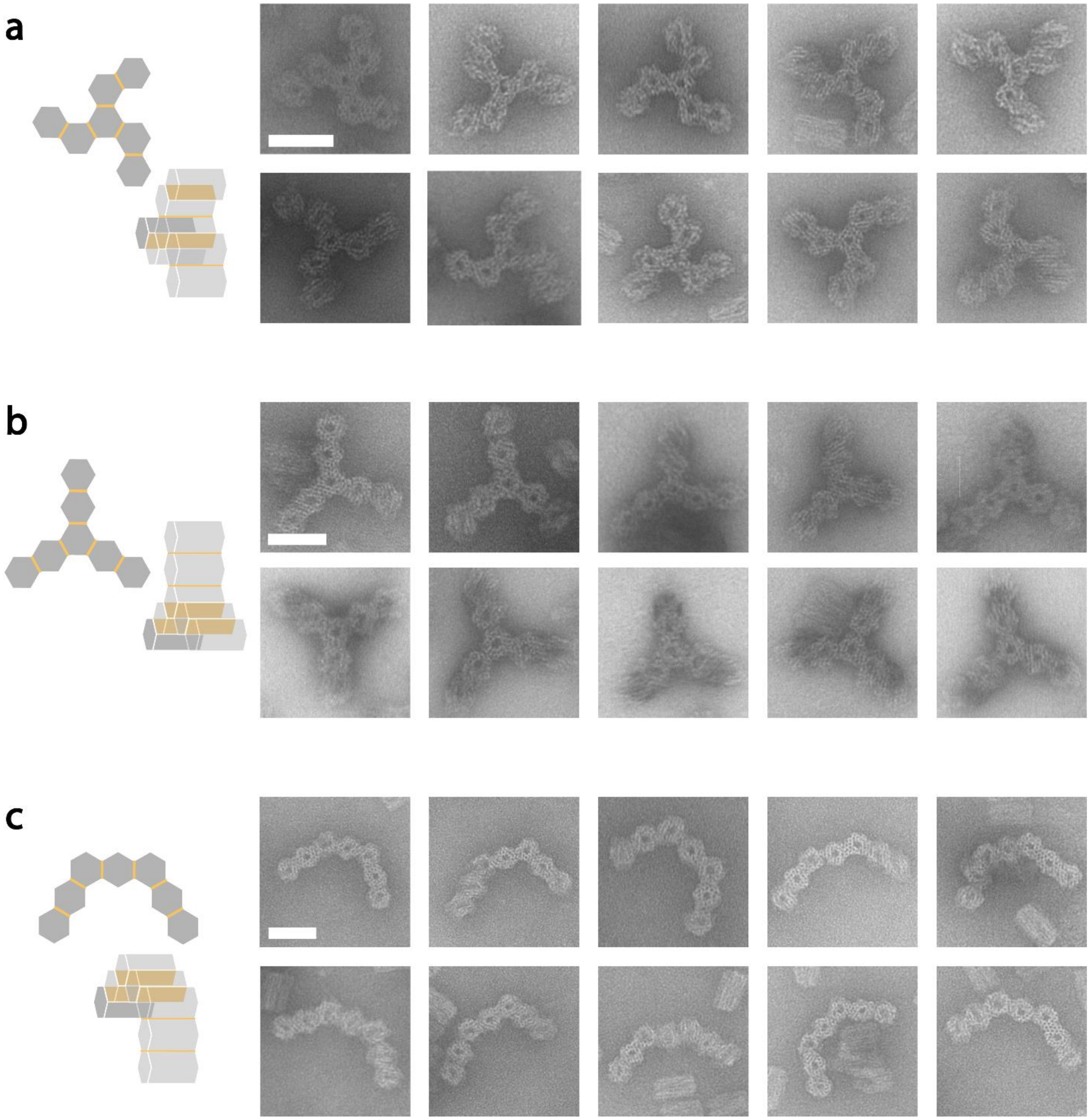

**Figure S17: Gallery of xy-structures 4** showing TEM micrographs of (**a**), (**b**), (**c**) different heptamers. Scale bars are 50 nm and hold for all micrographs of the respective structure.

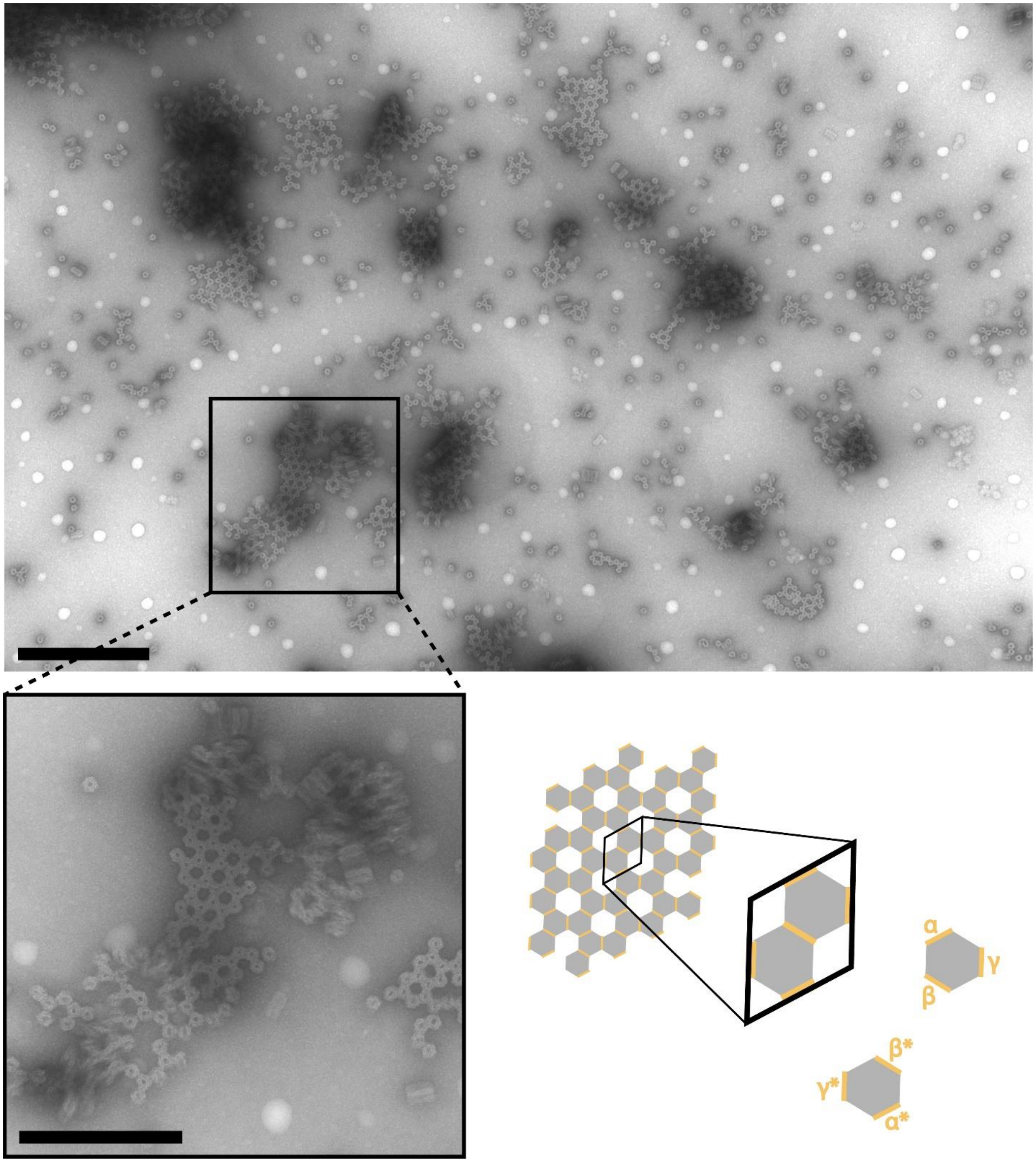

**Figure S18: Infinite xy-structures** were constructed by combining αβγ and α*β*γ* monomers. Together they form a unit cell of more than 1 500 $nm^2$. Enlarged is a piece of ~ 46 monomers spanning an area larger than 36 000 $nm^2$. Scale bar of the overview picture is 500 nm and 250 nm for the enlarged part.

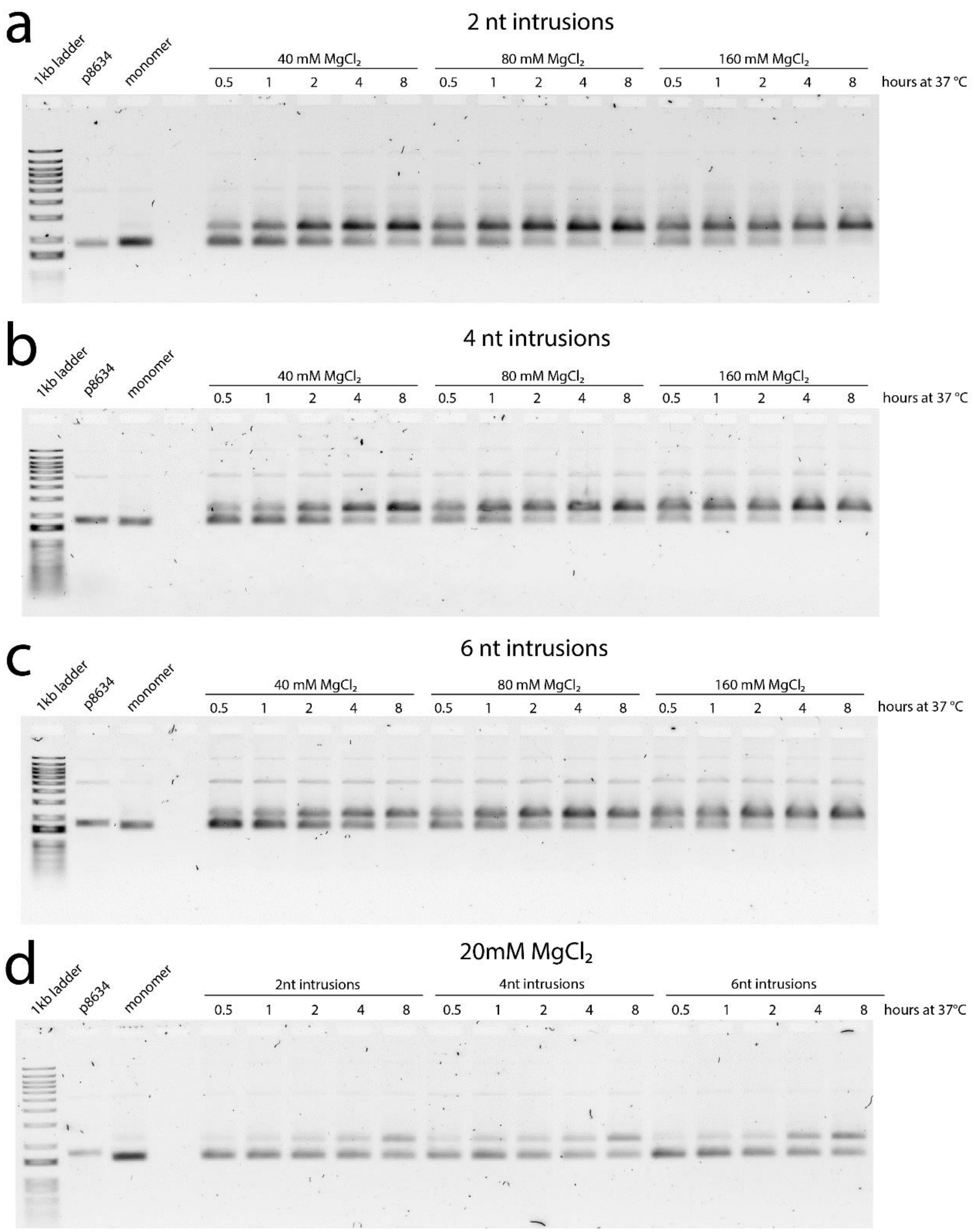

**Figure S19: AGE shift assays of xy-assembly 1:** complementary monomers with (**a**) 2 nt, (**b**) 4 nt, and (**c**) 6 nt staple intrusion were added to buffers containing a total $MgCl_2$ concentration of 40 mM, 80 mM, or 160 mM each, 8, 4, 2, 1, or 0.5 h before AGE was started. (**d**) Complementary monomers with 2 nt, 4 nt and 6 nt were added in a buffer containing 20 mM $MgCl_2$, at time points 8, 4, 2, 1, or 0.5 h before the samples were transferred to the gel. Data also used in Figure 2.

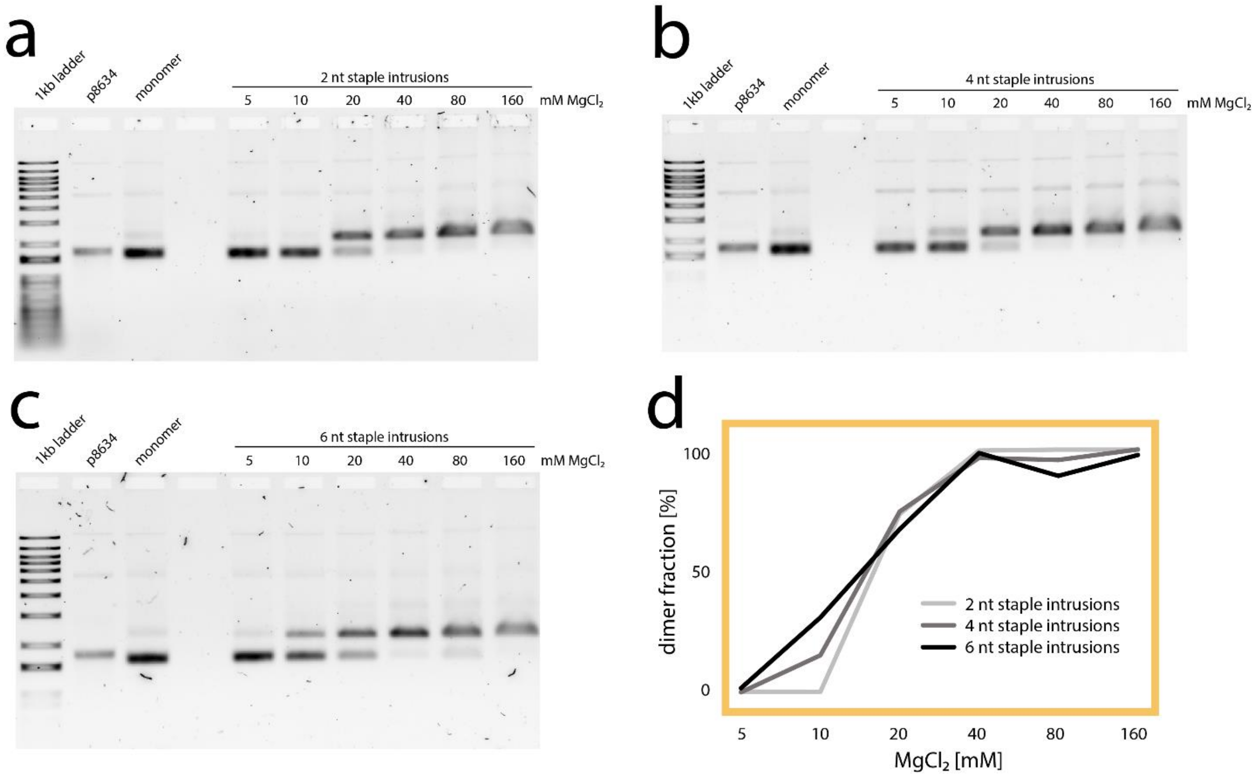

**Figure S20: AGE shift assays of xy-assembly 2**: Analyzing the dimerization after 24 h: complementary monomers with (**a**) 2 nt, (**b**) 4 nt, and (**c**) 6 nt staple intrusion were added to buffers containing a total $MgCl_2$ concentration of 5 mM, 10 mM, 20 mM, 40 mM, 80 mM, or 160 mM each, 24 h before AGE was started. Samples were incubated at 37 °C. Data also shown in Figure 2. (**d**) Fraction of dimers at different $MgCL_2$ concentrations, as fraction of dimer band intensity to the sum of dimer and monomer band intensity. 100 % dimerization normalized to monomer band.

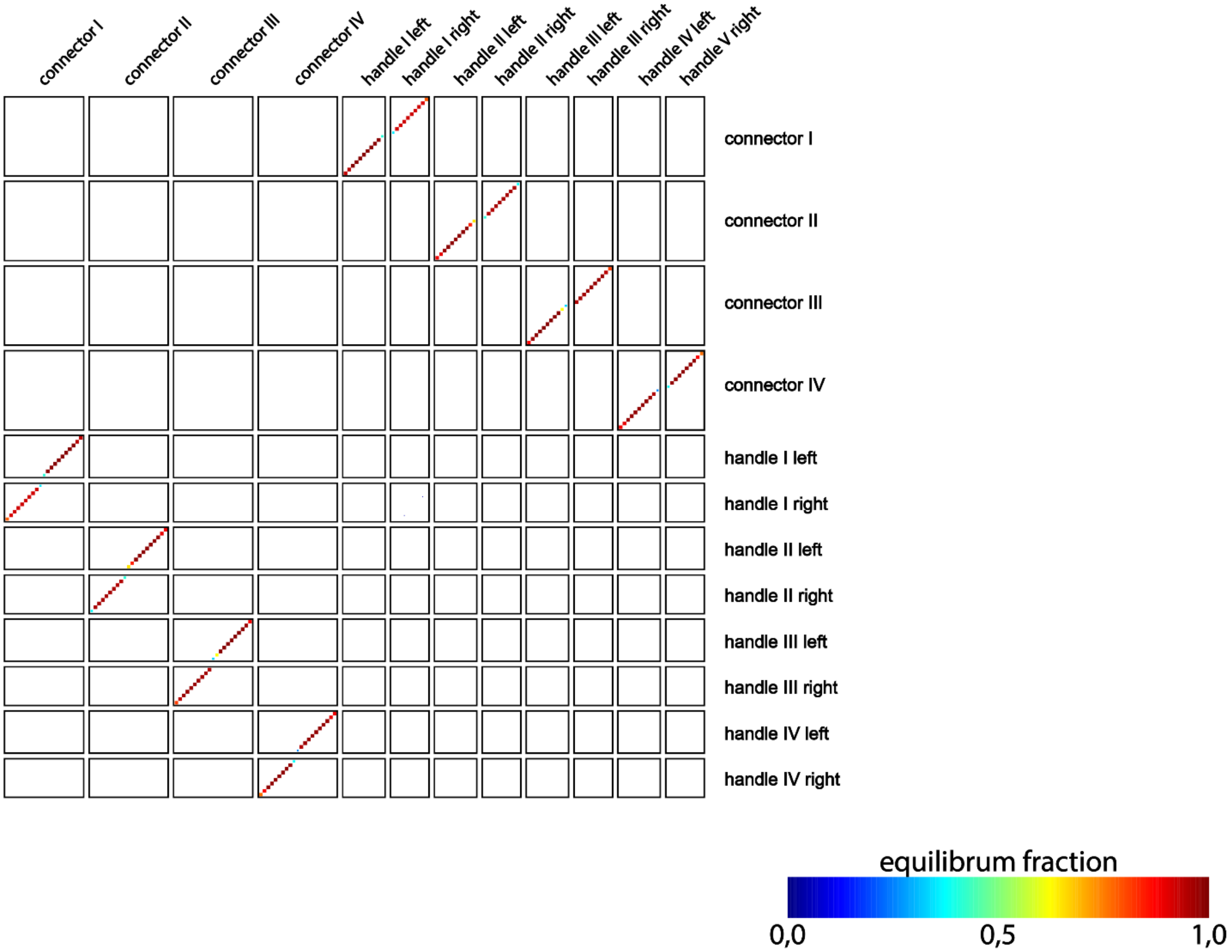

**Figure S21: NUPACK analysis of z-connector orthogonality:** Color scheme indicates complementary connectors and handles bind strongly and selectively with each other even in presence of the other handles and connectors. This shows mutual orthogonality. The simulation was conducted with NUPACK version 2.2 and the following options: 20°C, max. 3 strand complexes, 1 µM per strand, “Serra and Turner 1995”, 1 M NaCl.

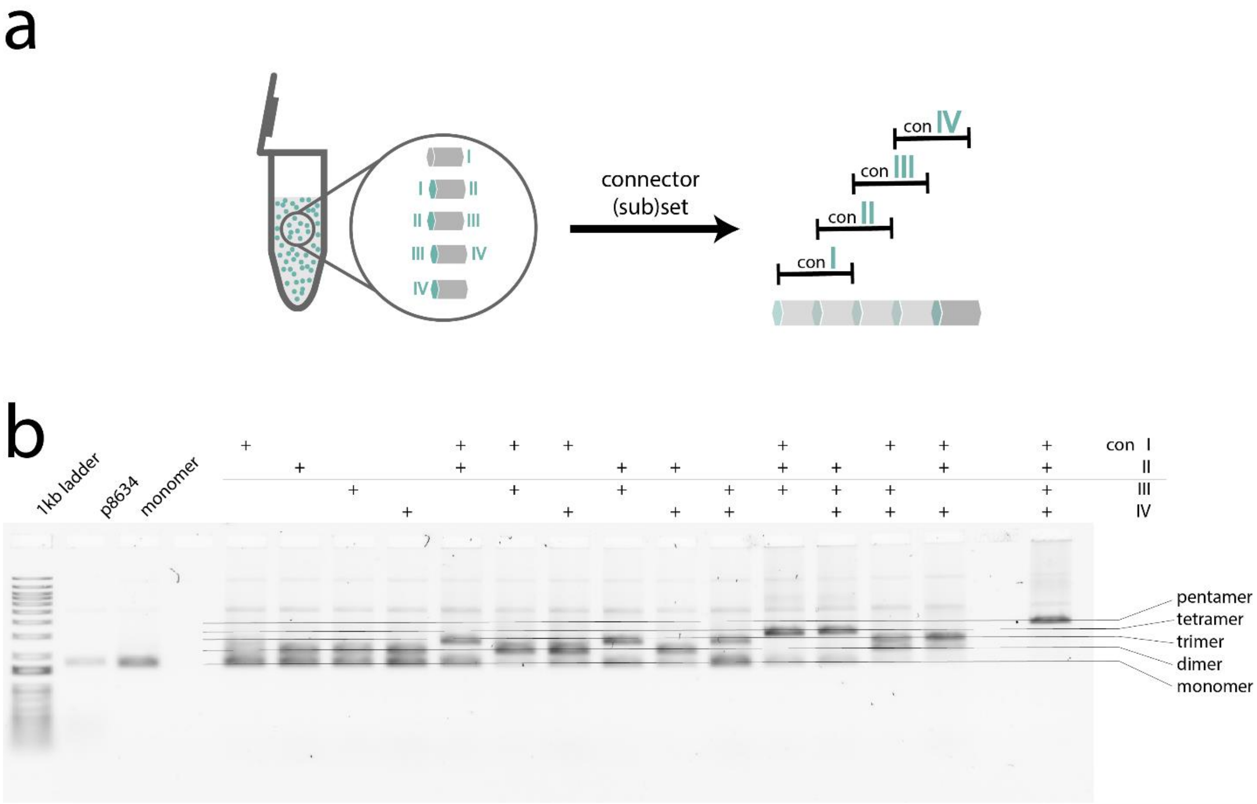

**Figure S22: z-assembly permutations:** (**a**) Schematic of the experiment: To an equimolar mixture of five moDON monomers (zI-right, zI-left/zII-right, zII-left/zIII-right, zIII-left/zIV-right, and zIV-left) subsets of connectors were added and a multitude of moDON multimers in z-direction were created. (**b**) AGE shift assay of z-assembly permutations showed the designed connections of subsets of the moDONs exceptionally well. Addition of single connectors resulted in formation of one part dimers with three parts monomer left unconnected (as only two types of monomers are able to form dimers). Addition of two connectors resulted either in one part trimer and two parts monomers (conI+conII, conII+conIII, or conIII+conIV) or in two parts dimer and one part monomer (conI+conIII, conI+conIV, or conII+conIV). Addition of three connectors resulted either in formation of one part tetramer and one part monomer (conI+conII+conIII or conII+conIII+conIV) or in one part trimer and one part dimer (conI+conIII+conIV or conI+conII+conIV). Addition of all four connector strands yields pentamers. Data was also used in Figure 3.

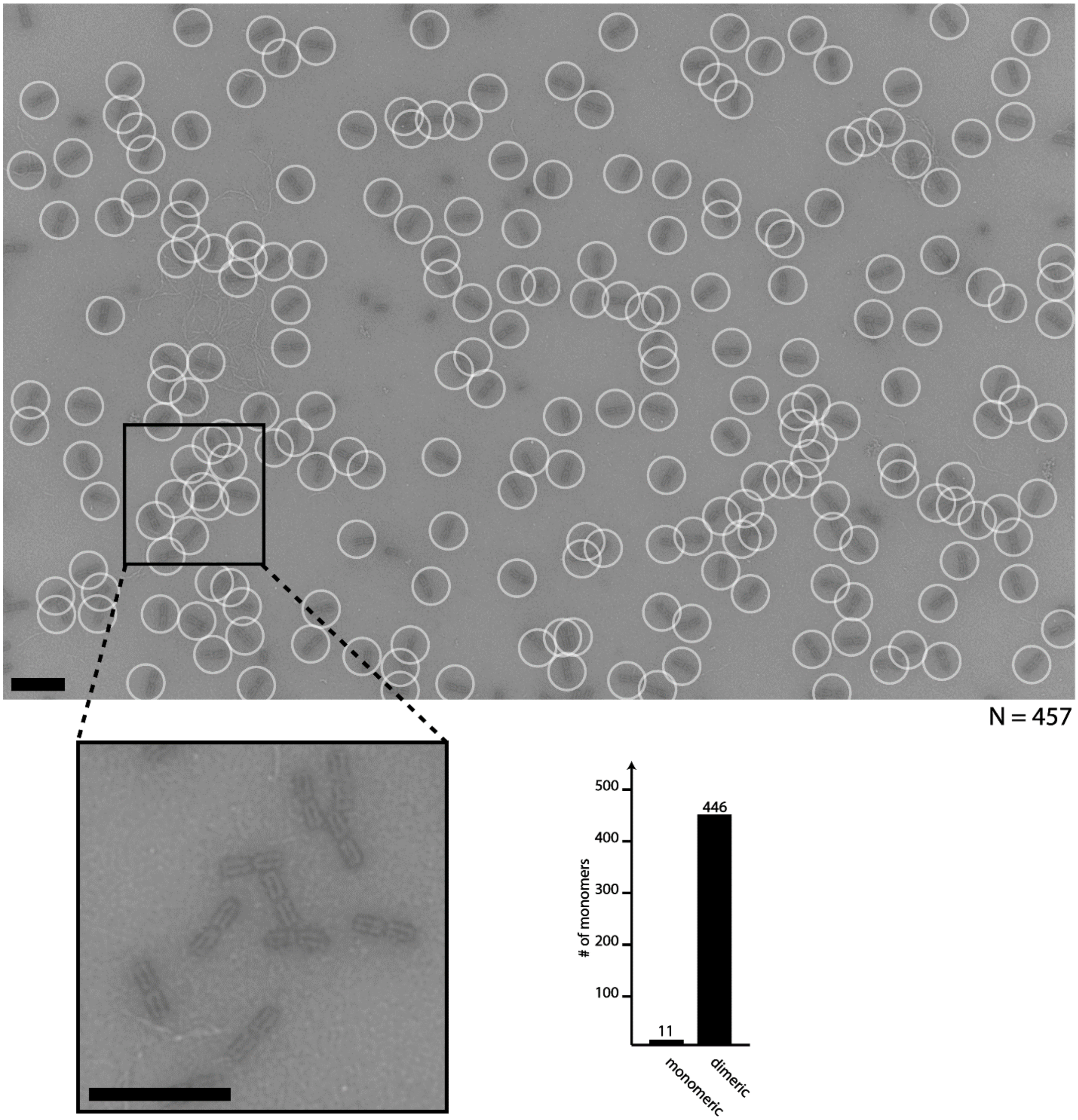

**Figure S23: Overview and statistic of z-dimer formation**. Analysis of the TEM micrograph shows a yield of 97.6 % of dimers, as calculated by the fraction of monomers in the designed structure ($N_{dimer}$ = 446) to the total amount of monomers ($N_{\Sigma}$ = 457). Scale bars are 200 nm.

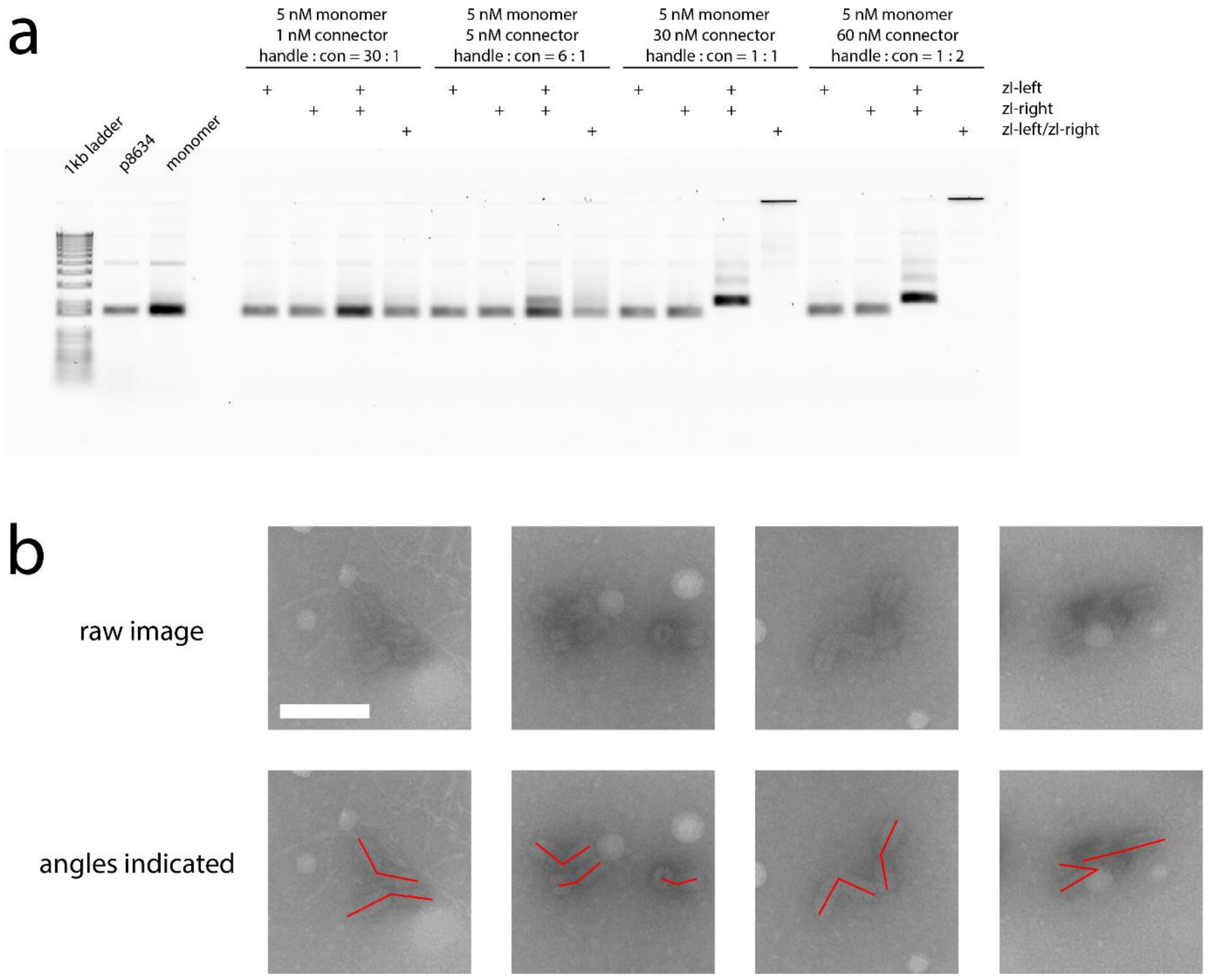

**Figure S24: z-connections with various connector concentrations** (**a**) were constructed from moDONs with handles at the left, the right or both sides. Ratio of handles to connectors was varied from 30:1 to 1:2. Full dimerization, or multimerization (for moDONs with both connection sites) was reached with 1:1 handle : connector ratio. We also noted, that for the 1:1 ratio the infinitely multimerized superstructures did not migrate through the gel, indicating a very high degree of multimerization. The lower, 6:1 ratio of handles to connectors, yielded a low fraction of moDONs dimerized, and further division of connector concentration by a factor of five, did not yield dimers at all. (**b**) TEM micrographs of z-dimers with equimolar connector-handle ratio. Connections are mostly bent along the z-axis. Below the raw micrograph, the same image is duplicated, with the angles between both monomers indicated in red. Scale bar is 100 nm and holds for all micrographs.

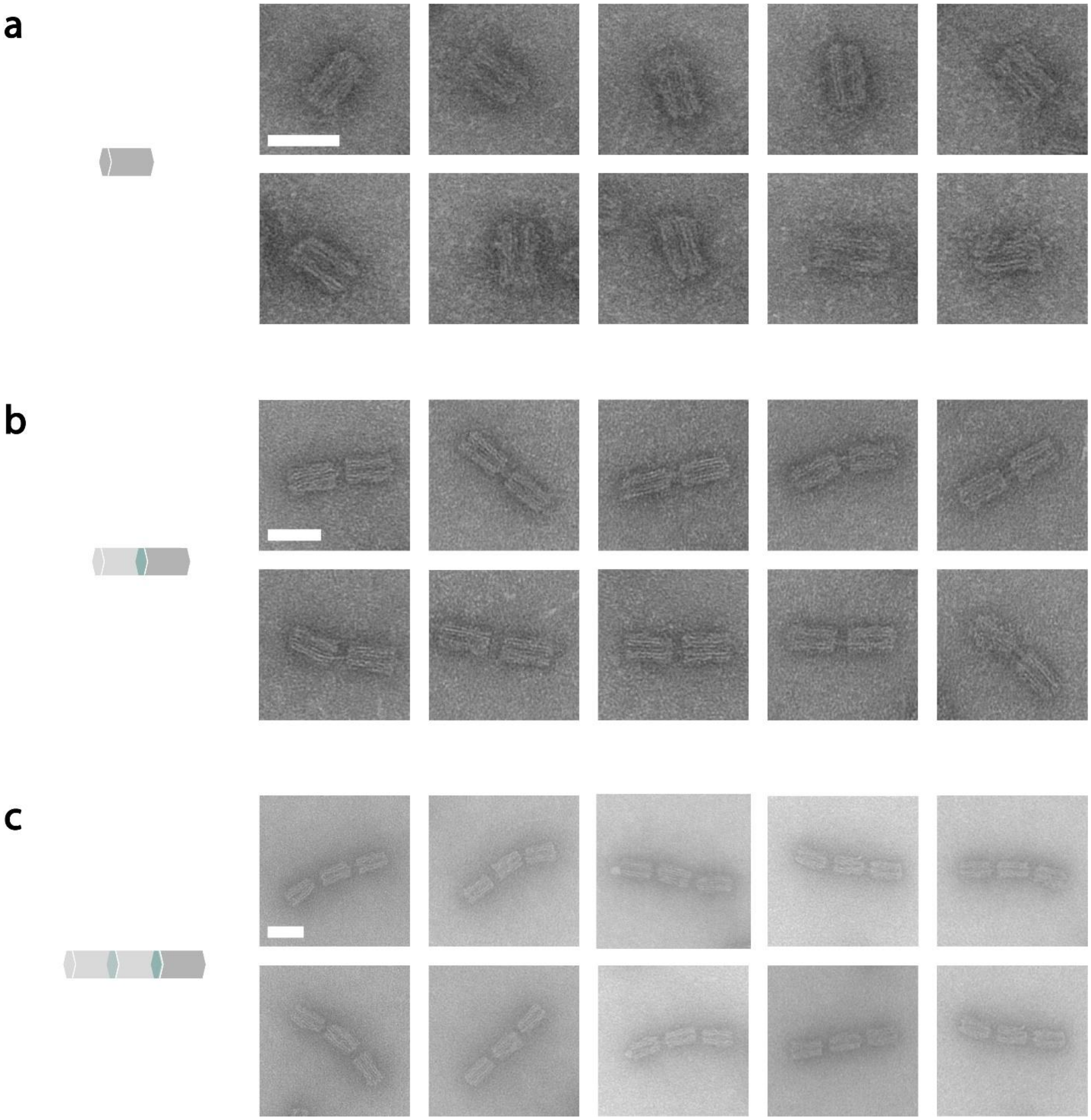

**Figure S25: Gallery of z-structures 1** showing TEM micrographs of (**a**) monomers (**b**) dimers and (**c**) trimers formed by addition of connectors in 5-fold excess to the moDON monomers. Scale bars are 50 nm and hold for all micrographs of the respective structure.

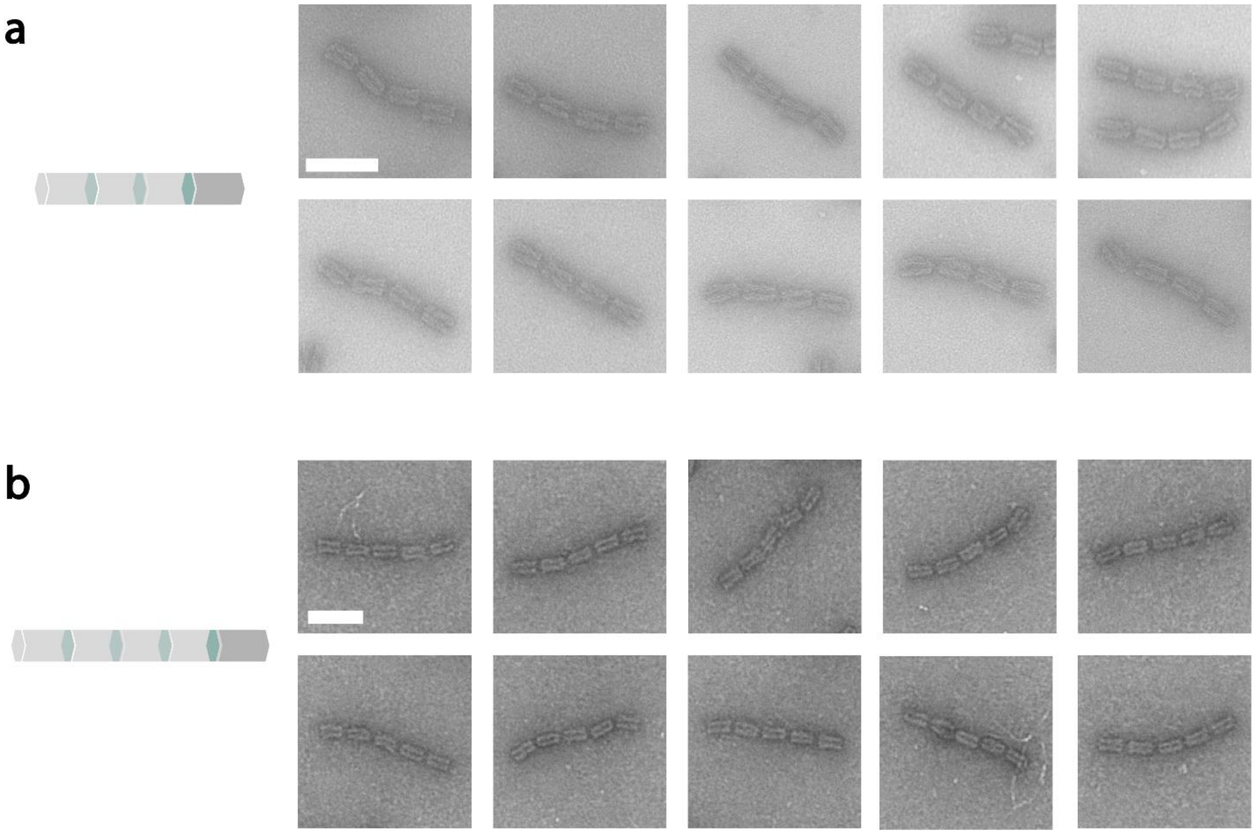

**Figure S26: Gallery of z-structures 2** showing TEM micrographs of (**a**) tetramers and (**b**) pentamers and formed by addition of connectors in 5-fold excess to the moDON monomers. Scale bars are 100 nm and hold for all micrographs of the respective structure.

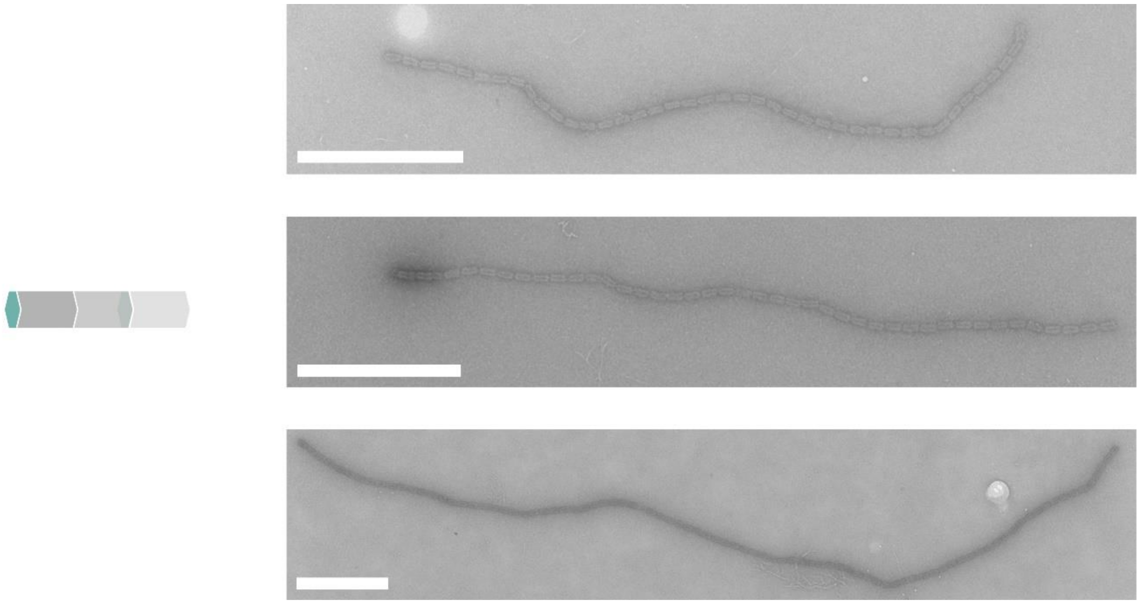

**Figure S27: Infinite tube with monomeric subunit** is constructed with the moDON zI-left/zI-right. z-connections are indicated in turquois. Number of subunits (= monomers) is 41, 42, and 88 (from top to bottom), corresponding to approximately 2 µm, 2 µm, and 4.4 µM length, and 0.23 GDa, 0.23 GDa, and 0.49 GDa weight, respectively. Data of the same experiment is shown in Figure 3. Scale bars are 500 nm.

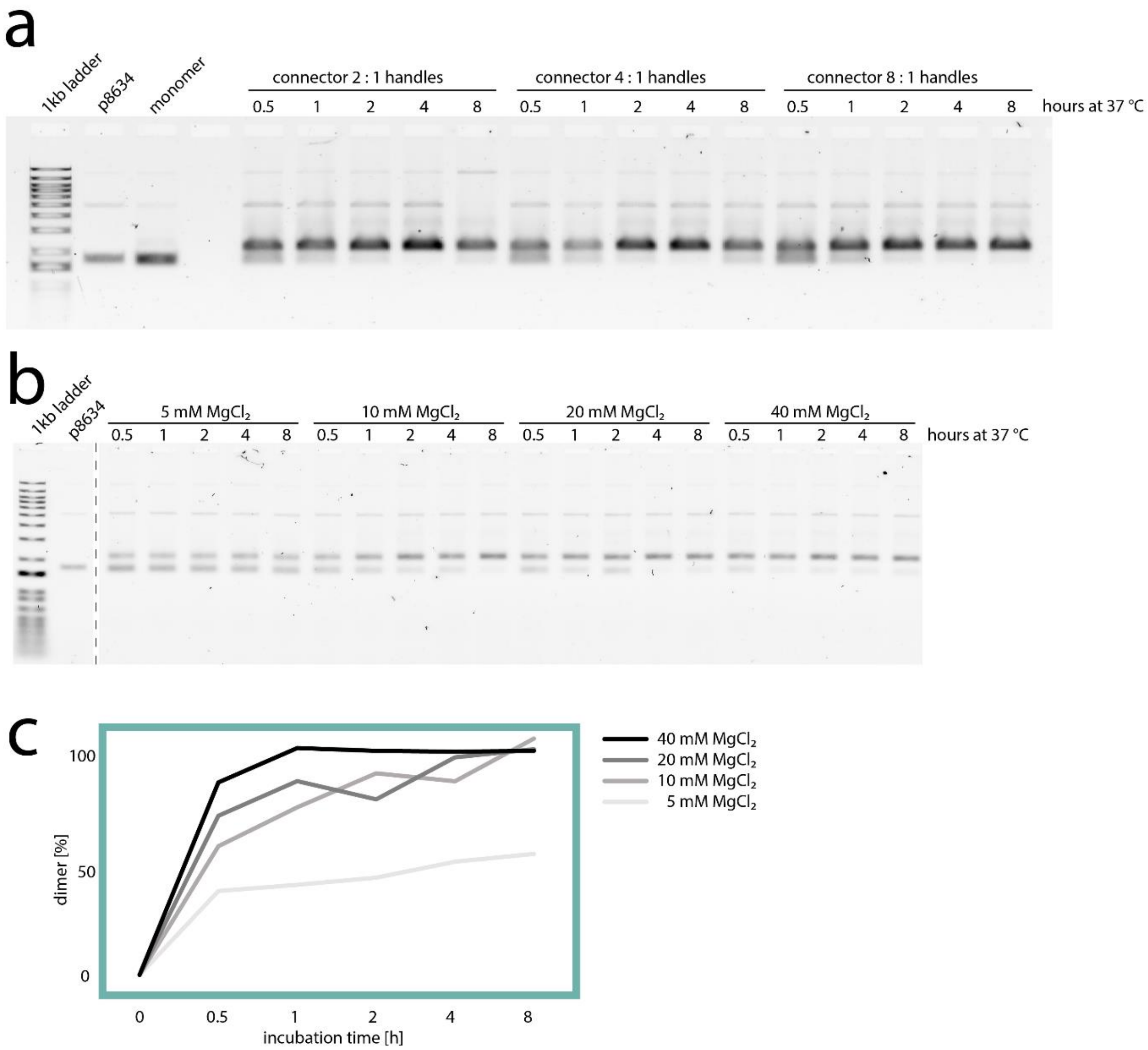

**Figure S28: AGE shift assays of z-assembly:** (**a**) Influence of connector excess: Dimers in z-direction were formed by addition of connector strands in 2-, 4-, or 8-fold excess to handles, each 8, 4, 2, 1, or 0.5 h before the samples were transferred to the gel. Data was also used in Figure 3. (**b**) Influence of $MgCl_2$ concentration: z-directional dimers were formed by addition of connector strands in 5-fold excess to handles in TAE buffer containing 5, 10, 20, or 40 mM $MgCl_2$, each 8, 4, 2, 1, or 0.5 h before the samples were transferred to the gel and the AGE was started. (**c**) $MgCl_2$ dependency of dimerization over time (from analysis of gel bands: comparison of fraction of dimer band intensity to the sum of dimer and monomer band intensity). 100% dimerization normalized to monomer band. 0 h incubation time was set to 0 % dimerization manually.

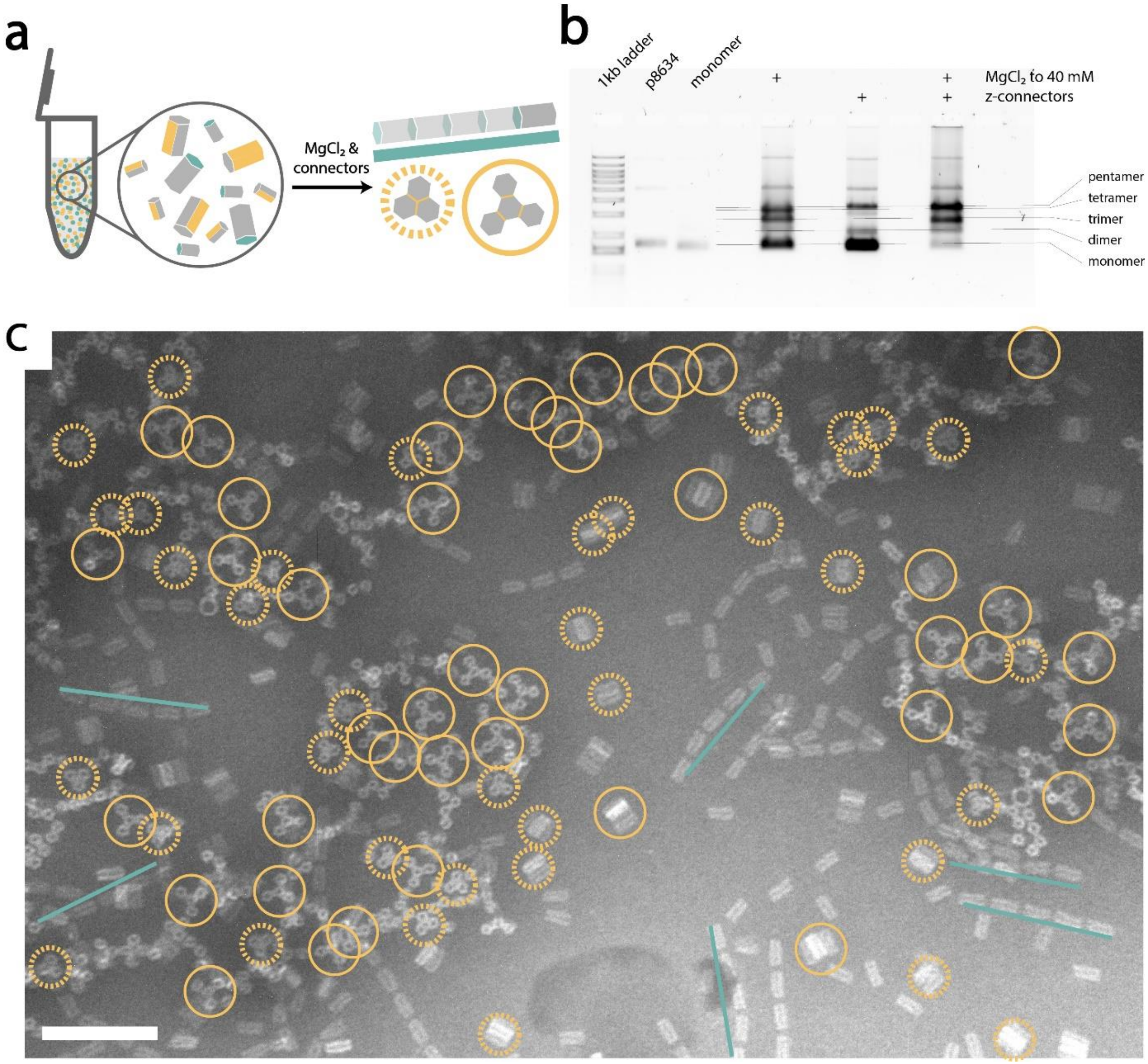

**Figure S29: Parallel assembly:** (**a**) Schematic of experimental setup: $MgCl_2$ and z-connectors were added to the same mixture of 12 moDON monomers, and three distinctly different structures formed parallelly. The mutual orthogonality of xy- and z-connections, as well as the connection sites themselves with respect to each other allows assembly of several different structures in the same reaction vessel, without crosstalk. (**b**) AGE shift assay of parallel formation: Sole addition of $MgCl_2$ leads to formation of xy-trimers and xy-tetramers, sole addition of connectors with low $MgCl_2$ leads to formation of z-pentamers, each trigger leaving the uninvolved moDONs monomeric. Addition of both leads to parallel assembly of all three structures. 20 fmol of the following monomers were used: αβ*, βγ*, and α*γ for the xy-trimer, δεζ, δ*, ε*, and ζ* for the xy-tetramer, and zI-right, zI-left/zII-right, zII-left/zIII-right, zIII-left/zIV-right, and zIV-left for the z-pentamer. (**c**) TEM micrograph of parallelly assembled moDON superstructures: xy-connections are indicated in yellow, z-connections in turquois. In the TEM micrograph, several xy-trimers are indicated with a dotted yellow circle, xy-tetramers with a large yellow circle, and z-pentamers are indicated with a turquoise line. Data of the same experiment is shown in Figure 4. Scale bar is 200 nm.

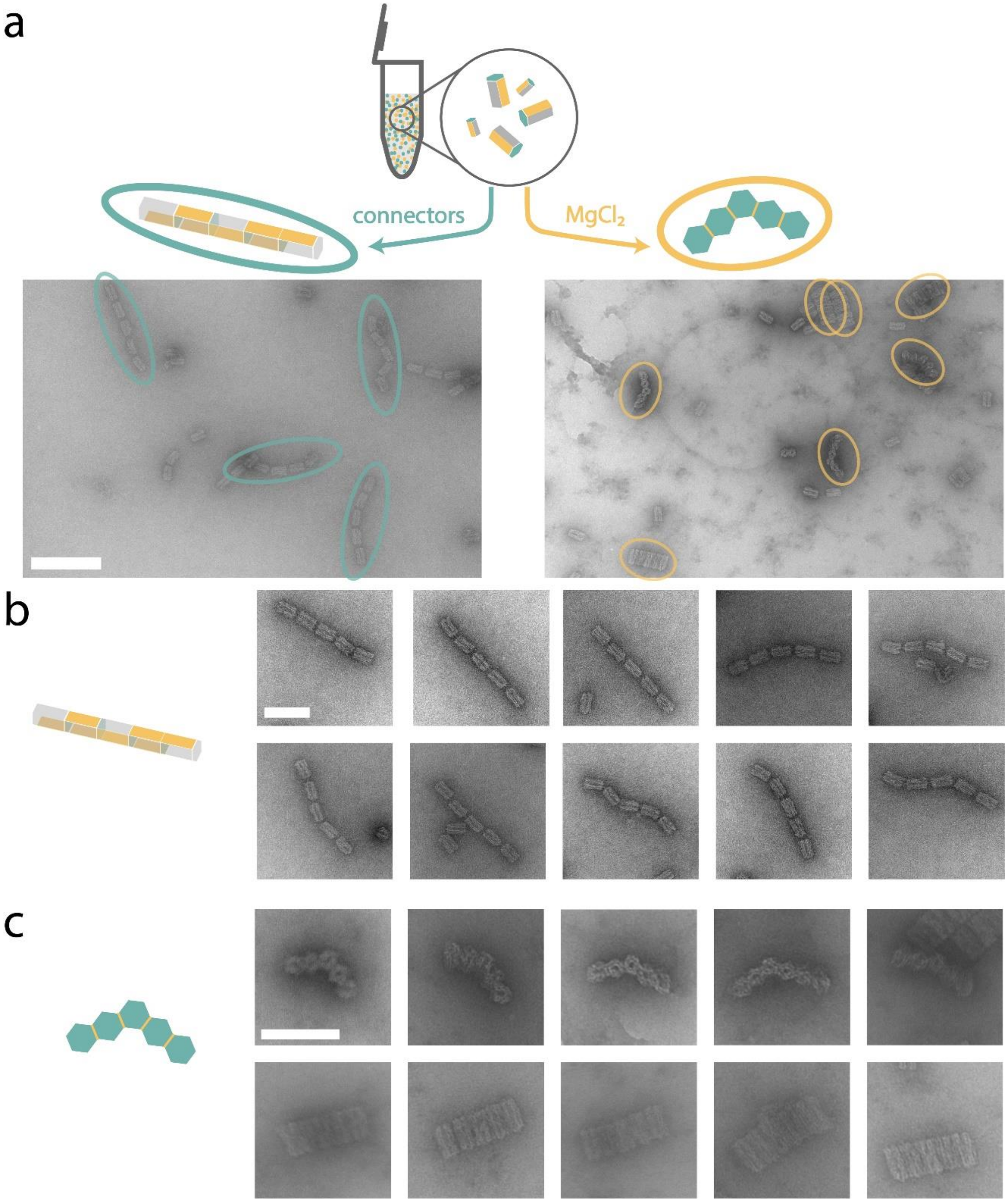

**Figure S30: Selective assembly** of different structures from the same set of moDONs: Schematic (**a**) shows that the resulting structure depends on whether z-connectors or $MgCl_2$ of 40 mM were added. xy-connections are indicated in yellow, z-connections in turquois. The moDONs used in this experiment are β*/zI-left/zII-right, βε*/zI-right, γε/zII-left/zIII-right, γ*ζ/zIV-left, and ζ/zIII-left/zIV-right. Data of the same experiment is shown in Figure 4. Addition of only 40 mM $MgCl_2$ yields a pentamer in the xy-direction. Addition of only z-connectors yields a pentamer in z-direction. Gallery of (**b**) z-assembled pentamers, and (**c**) xy-assembled pentamers. Scale bar for (**a**) is 200 nm and holds for both micrographs, scale bar for (**b**) and (**c**) is 100 nm and holds for all micrographs of the respective structure.

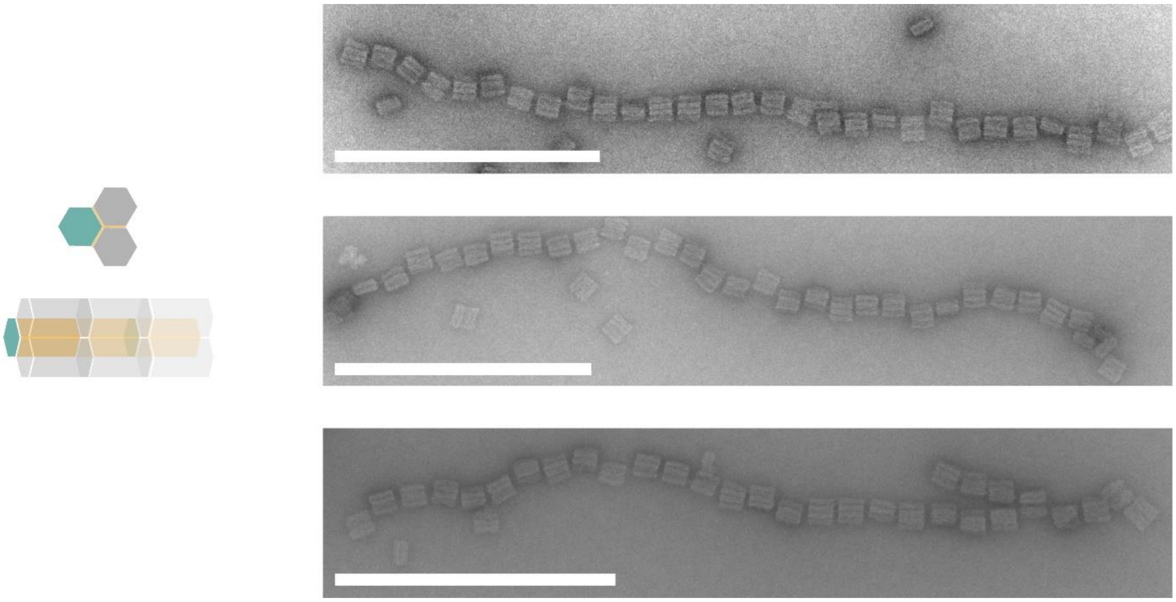

**Figure S31: Infinite tubes with trimeric subunits** are constructed with three moDONs: αβ*, βγ*, and α*γ/zI-left/zI-right. xy-connections are indicated in yellow, z-connections in turquois. The number of subunits is 30, 31, and 28 (from top to bottom), corresponding to ~1,5 µm length and ~0.5 GDa weight, respectively. Scale bars are 500 nm.

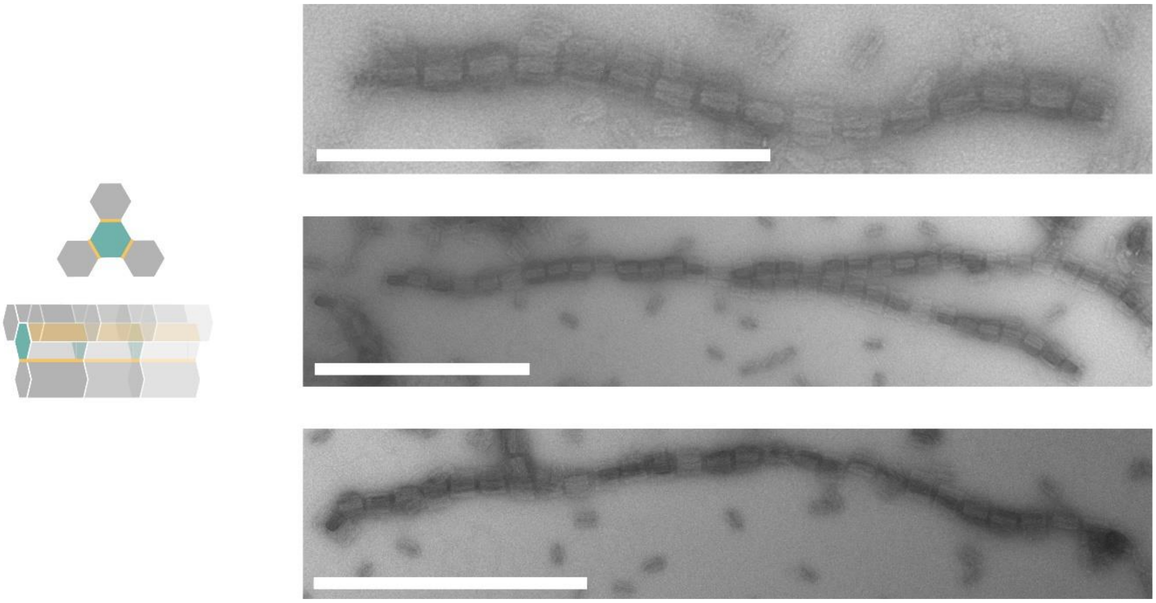

**Figure S32: Infinite tubes with tetrameric subunits** are constructed with four moDONs: δεζ/zII-left/zII-right, δ*, ε*, and ζ*. xy-connections are indicated in yellow, z-connections in turquois. Number of subunits is 16, 34, and 28 (from top to bottom), corresponding to ~0.8 µm, ~1.6 µm, and ~1.4 µm length and approximately 0.35 GDa, 0.76 GDa, and 0.63 GDa weight, respectively. Data of the same experiment is shown in Figure 4. Scale bars are 500 nm.

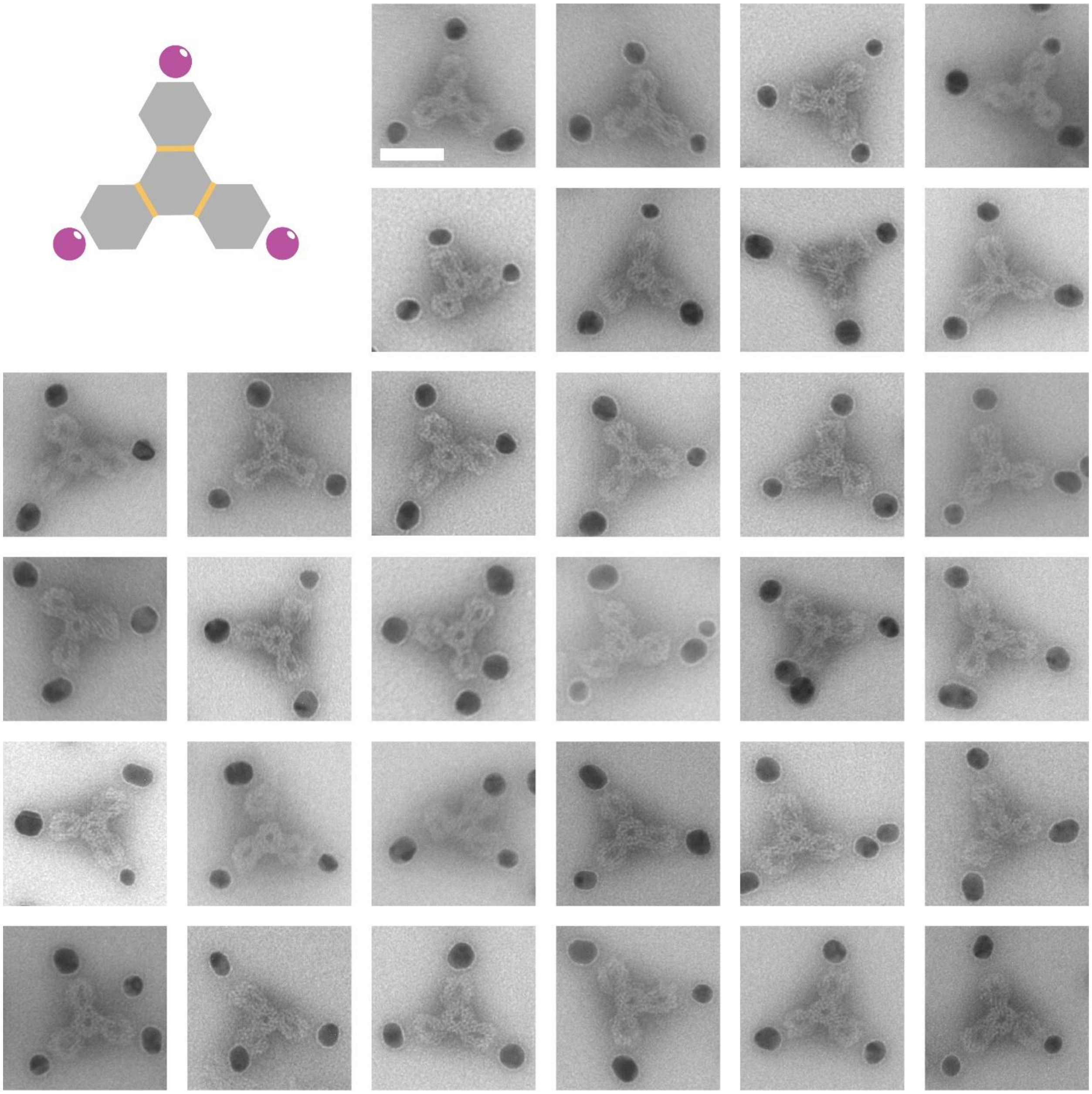

**Figure S33: xy-tetramer with selectively placed Au NPs** at the vertices. The central moDON has the connection sites δεζ, connecting to monomers δ* (Au NP handles in α direction), ε* (Au NP handles in β direction), and ζ* (Au NP handles in γ direction). Data of the same experiment is shown in Figure 4. Scale bar is 50 nm and holds for all micrographs.

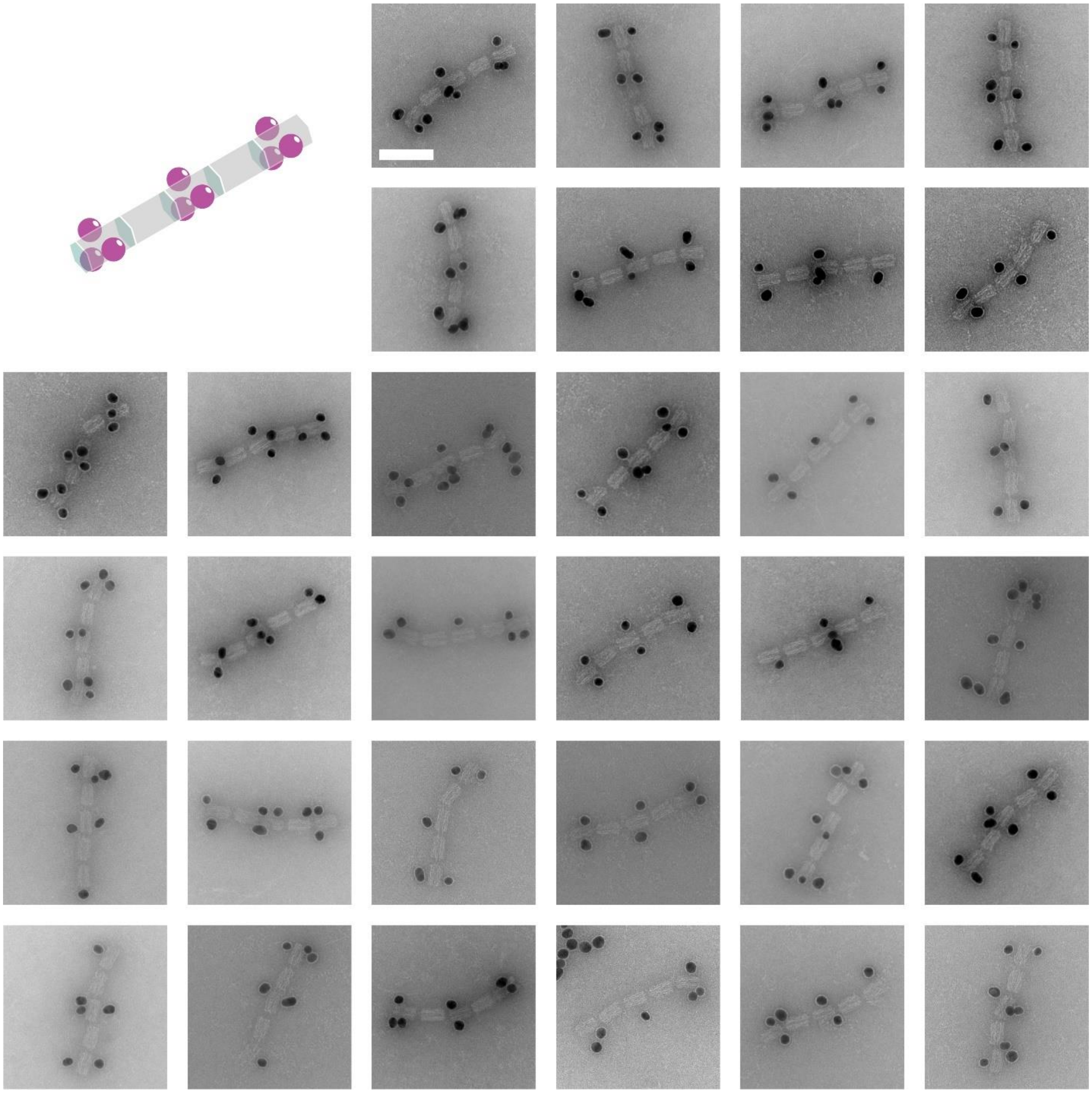

**Figure S34: z-pentamer with selectively placed Au NPs** on monomers one (zI-right), three (zII-left/zIII-right), and five (zIV-left) with three Au NP connection sites (modified αβγ connection sites). Data of the same experiment is also found in Figure 4. Scale bar is 100 nm and holds or all micrographs.

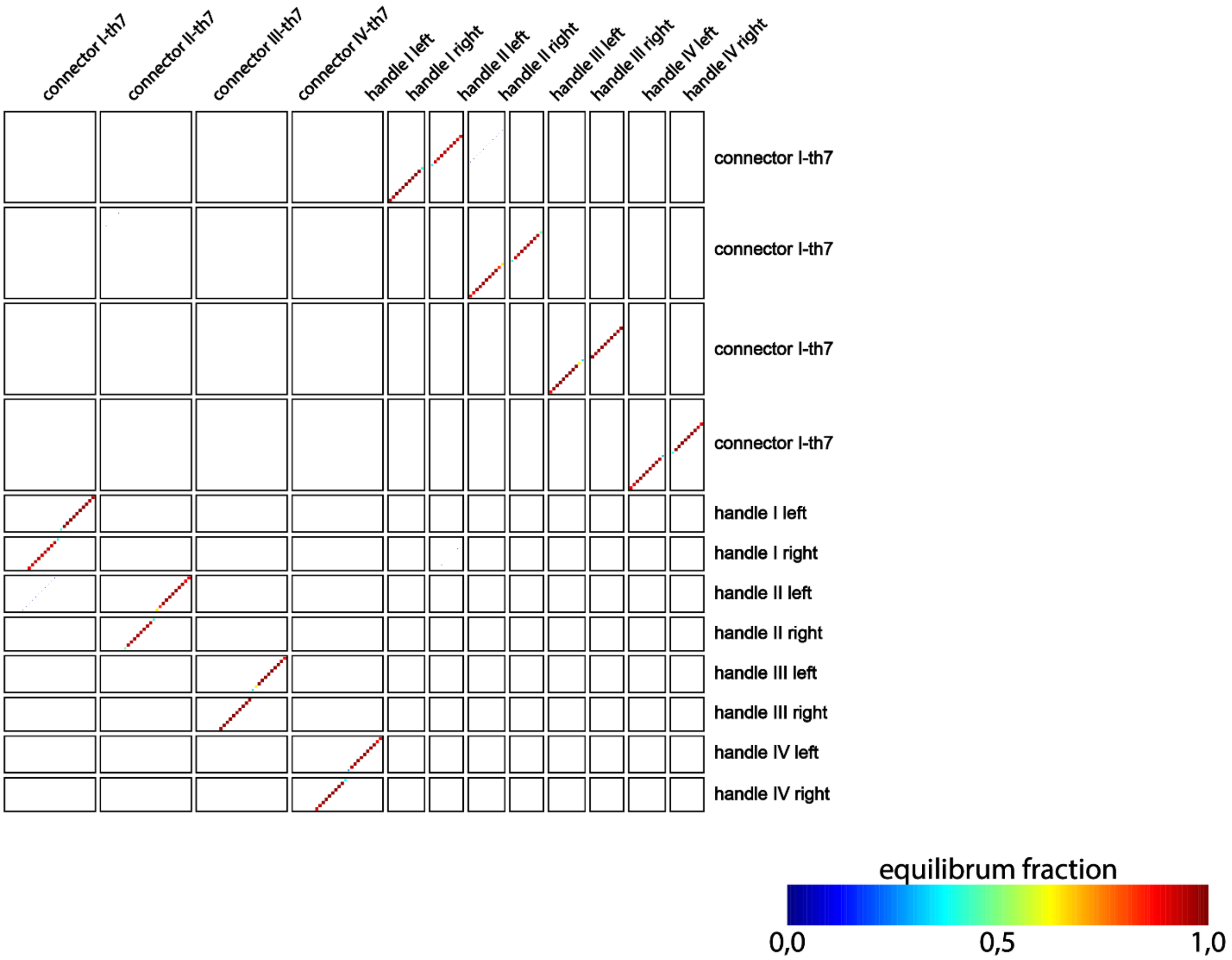

**Figure S35: NUPACK analysis of z-connector orthogonality with toeholds:** color scheme indicates complementary connectors and handles bind strongly and selectively with each other even in presence of the other handles and connectors. This shows mutual orthogonality. Connectors elongated with a toehold region were denoted with the respective Roman letter, indicating their sequence, and "-th7", indicating the addition of a 7 nt long toehold. The toehold does not seem to have a major influence on binding behavior (*cf*. Figure S21). The simulation was conducted with NUPACK version 2.2 and the following options: 20°C, max. 3 strand complexes, 1 µM per strand, "Serra and Turner 1995", 1 M NaCl. For respective sequences see Table S15.

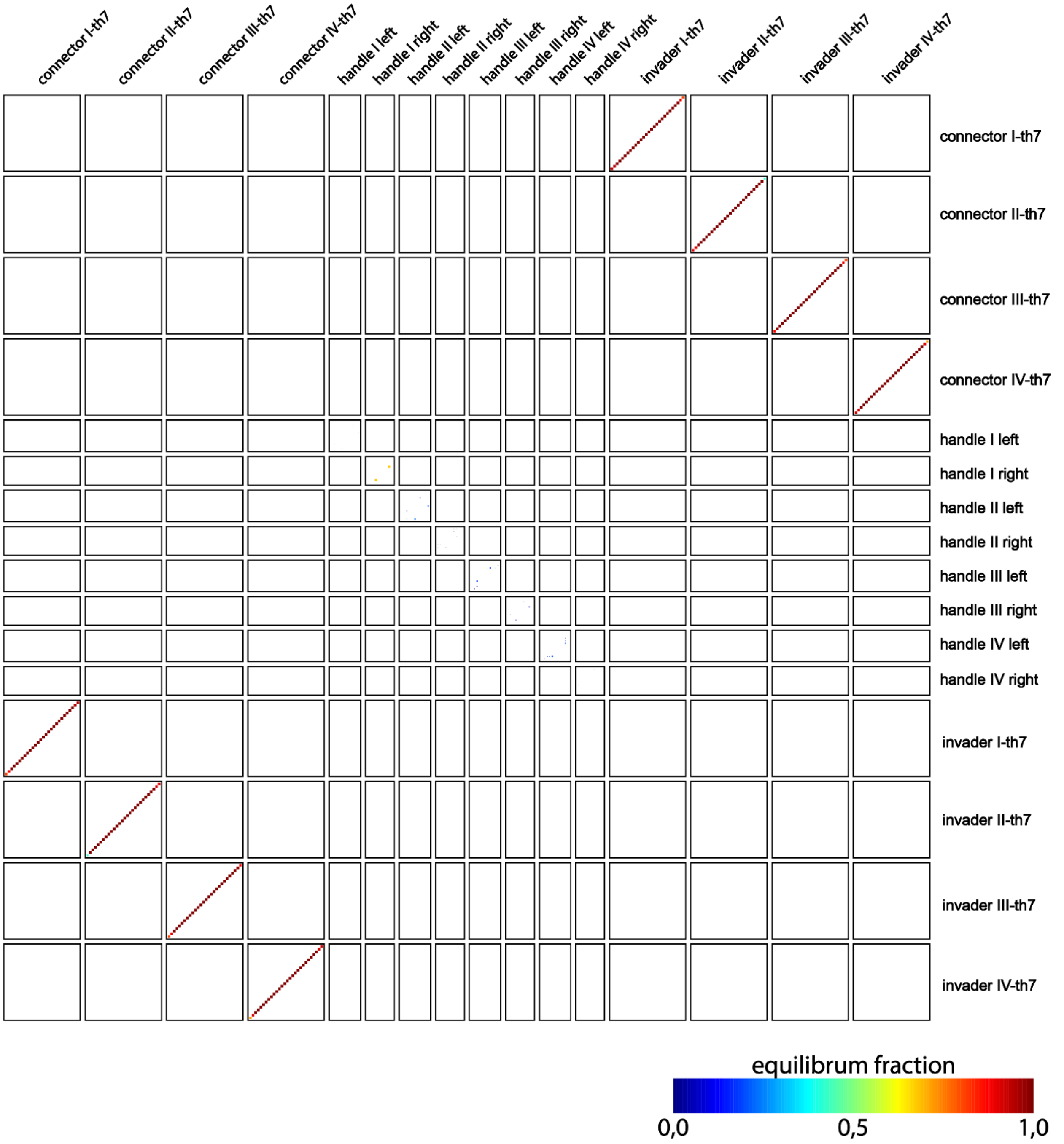

**Figure S36: NUPACK analysis of z-connectors in presence of invader strands:** color scheme indicates the binding of complementary connectors and invaders is favored over the binding of connectors to handles (*cf.* Figure S35). Further, crosstalk between non-complementary handles, connectors and invaders is vanishingly low, indicating retained orthogonality in the strand displacement process. Connectors and invaders elongated with a toehold region were denoted with their respective Roman letter, indicating their sequence, and "-th7", indicating a 7 nt long toehold. The simulation was conducted with NUPACK version 2.2 and the following options: 20°C, max. 3 strand complexes, 1 µM per strand, "Serra and Turner 1995", 1 M NaCl. For respective sequences see Table S15.

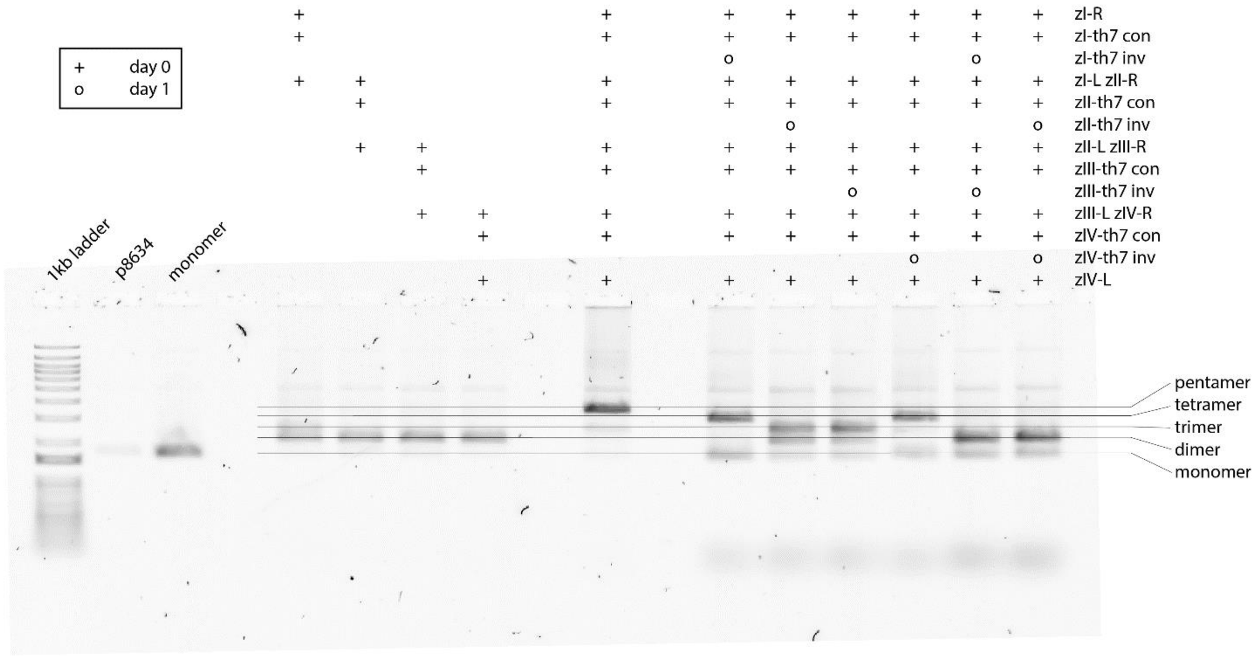

**Figure S37: z-disassembly permutations** of a pentamer, which was formed with 5-fold connector excess over handles over-night. Connectors and invaders elongated with a toehold region were denoted with their respective Roman letter, indicating their sequence, and "-th7" indicating the addition of a 7 nt long toehold. For sequences see Table S15. Invaders were added in 5-fold excess over connectors and again incubated over-night. Addition of invaders against connection zI or zIV resulted in disassembly to a tetramer and a monomer (see Figure S38), addition of zII or zIII invaders resulted in a dimer and a trimer (see Figure S39). Combination of zI and zIII invaders or zII and zIV invaders resulted in 2 dimers and one monomer (see Figure S40). Used moDON monomers: zI-right, zI-left/zII-right, zII-left/zIII-right, zIII-left/zIV-right, and zIV-left.

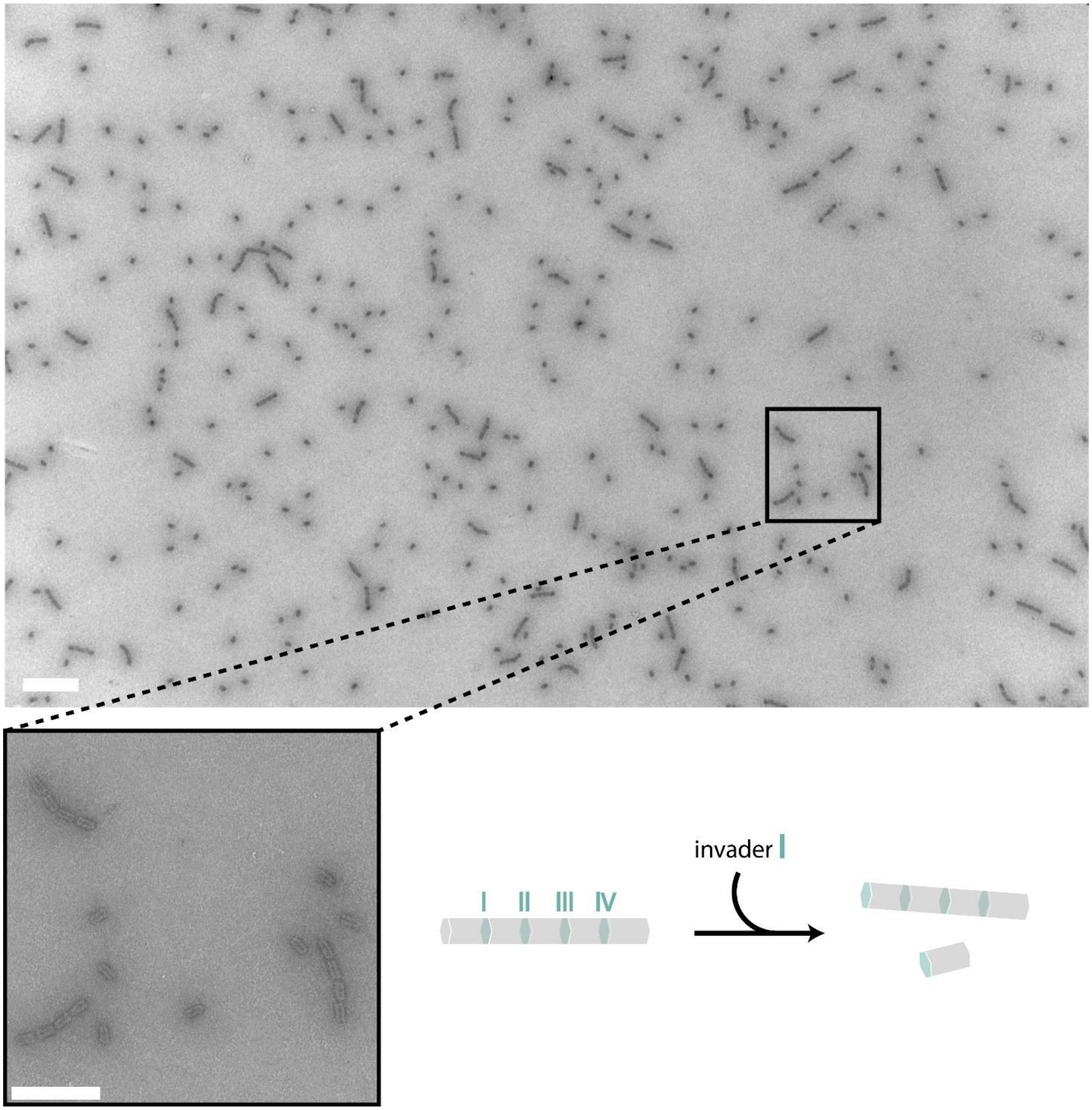

**Figure S38: TEM micrographs of selective z-disassembly 1:** To a fully formed pentamer of z-connections, invader strand I was added. Invader strand I disassembled the connections between the first and second moDON (from left to right) which resulted in a tetramer and a monomer. Scale bar is 500 nm for the overview picture and 200 nm for the enlarged part.

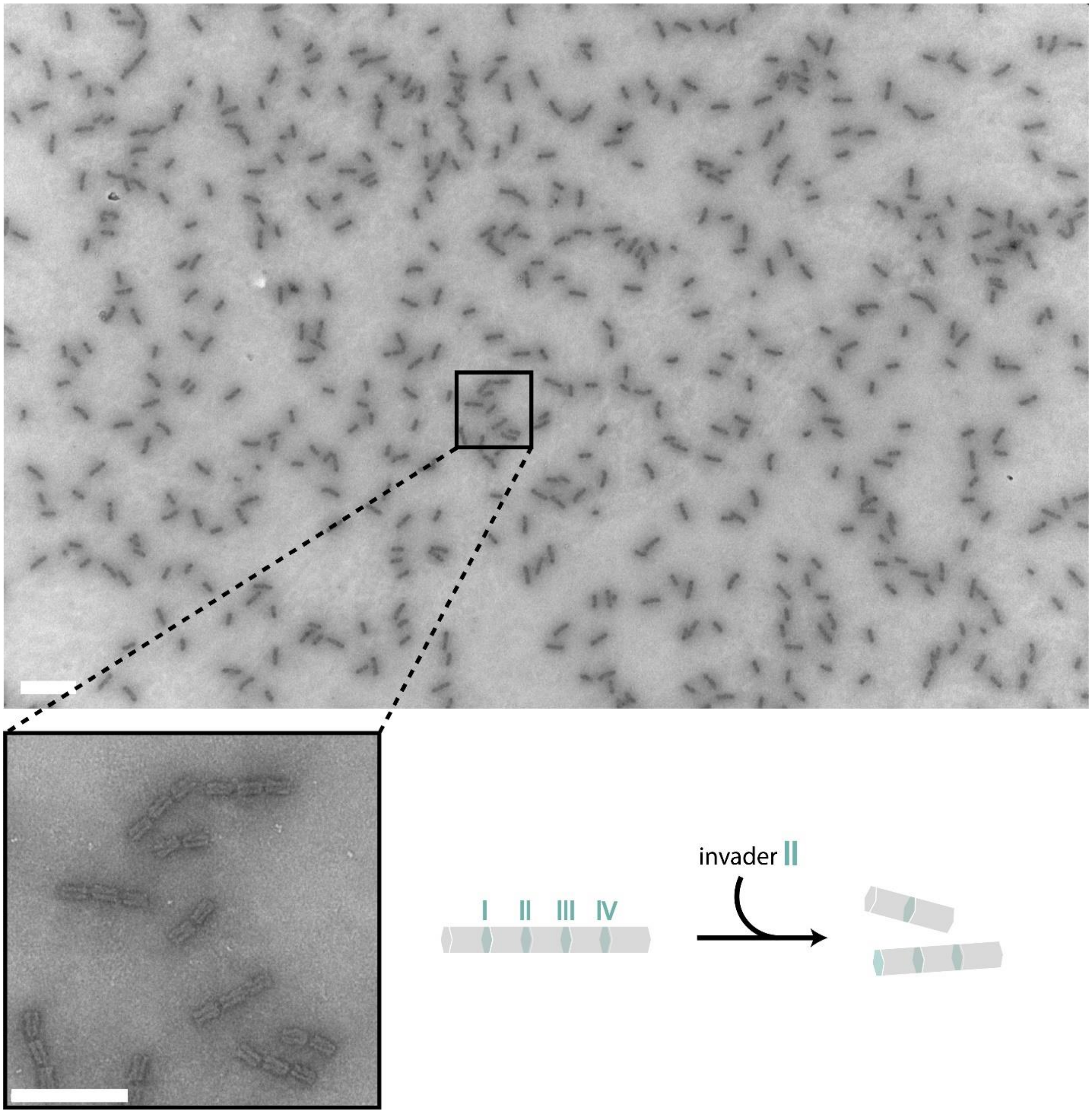

**Figure S39*:* TEM micrographs of selective z-disassembly 2:** To a fully formed pentamer of z-connections invader strand II was added. Invader strand II disassembled the connections between the second and third (from left to right) moDON, which resulted in dimers and trimers. Scale bars are 500 nm for the overview picture and 200 nm for the enlarged part.

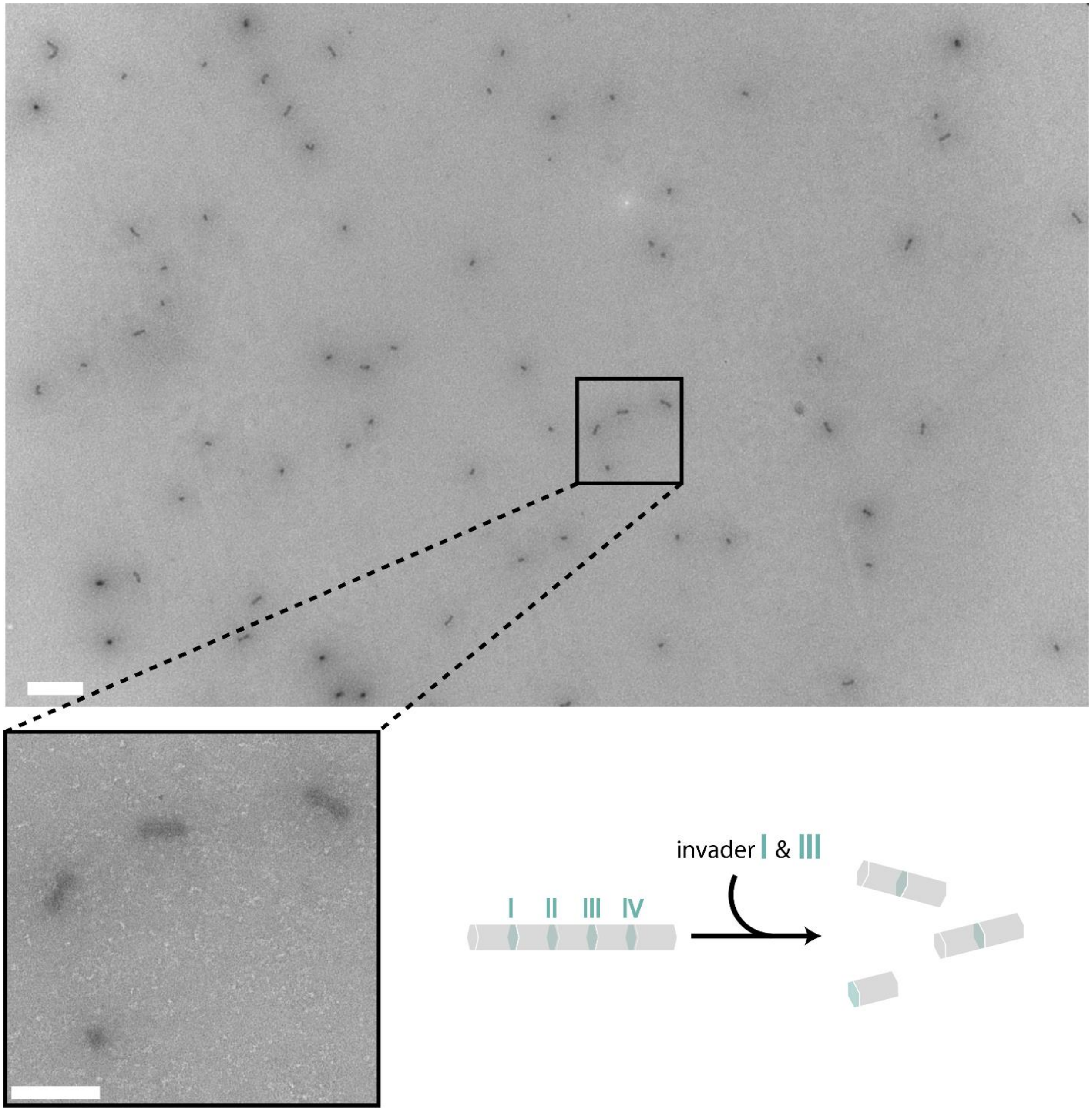

**Figure S40: TEM micrographs of selective z-disassembly 3:** To a fully formed pentamer of z-connections invader strands I and III were added. Invader strand I and III disassembled the connections between the first and second, as well as the third and fourth moDON (from left to right) which resulted in two dimers and one monomer, per pentamer. Scale bars are 500 nm for the overview picture and 200 nm for the enlarged part.

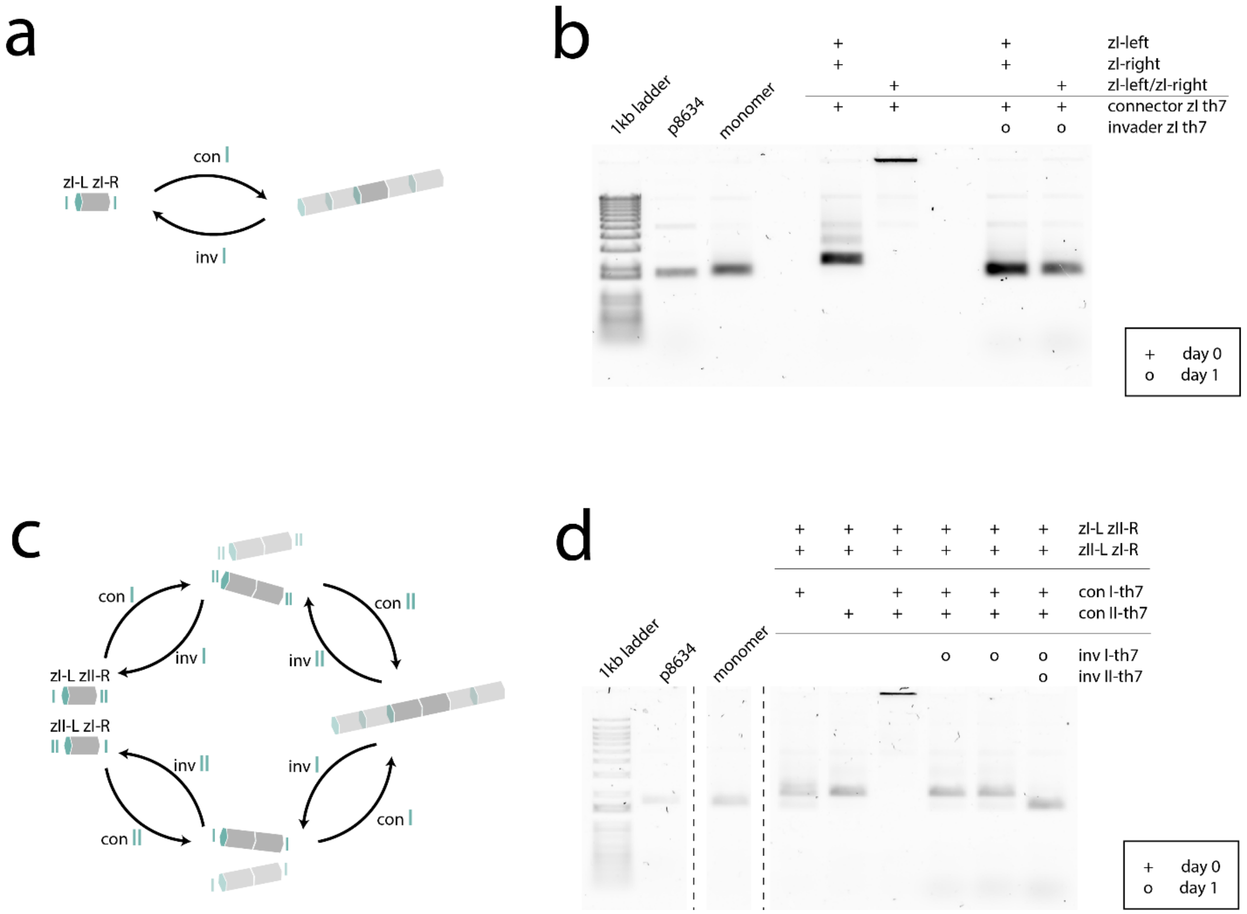

**Figure S41: Assembly & disassembly of periodic z-directional superstructures**: (**a**), (**b**) Homomultimeric or (**c**), (**d**) heteromultimeric. (**a**) Schematic showing assembly and disassembly pathway of the homomultimeric superstructure. (**b**) AGE shift assay of the assembly and disassembly: Addition of connector strand assembles the monomers into infinite superstructures, addition of the respective invader disassembles them again into monomers. (**c**) Schematic showing assembly and disassembly pathways of the heteromultimeric superstructure. (**d**) AGE shift assay shows assembly of both possible subunits through addition of the respective connector strand, and assembly of infinite tubes, when both connectors are added. The tubes can be disassembled into dimers or monomers, depending on whether only one or both invaders are added. Gels were imaged on day 2.

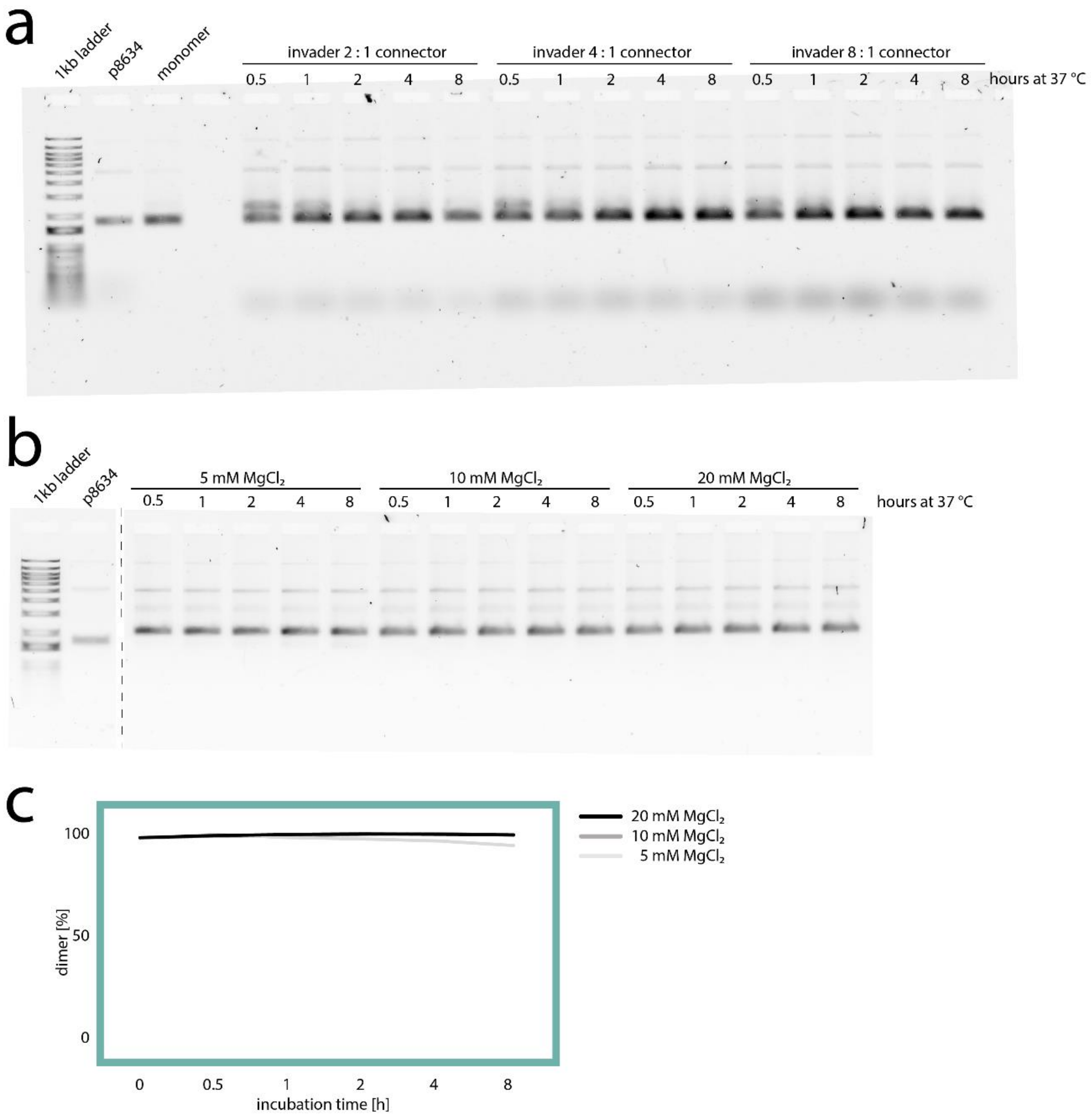

**Figure S42: AGE shift assay of z-disassembly:** (**a**) Influence of invader excess: z-directional dimers were formed through overnight, the next day invaders were added in 2-, 4-, or 8-fold excess over connector strands, each 8, 4, 2, 1, or 0.5 h before the samples were transferred to the gel. (**b**) Influence of $MgCl_2$ concentration: z-directional dimers were formed over-night. They were diluted in buffers with a total $MgCl_2$ concentration of 5 mM, 10 mM, or 20 mM each, 8, 4, 2, 1, or 0.5 h before the samples were transferred to the gel. (**c**) $MgCl_2$ dependency of dimer disassembly (based on analysis of gel band intensities: fraction of dimer band intensity to the sum of dimer and monomer band intensity). 100% dimerization normalized to monomer band.

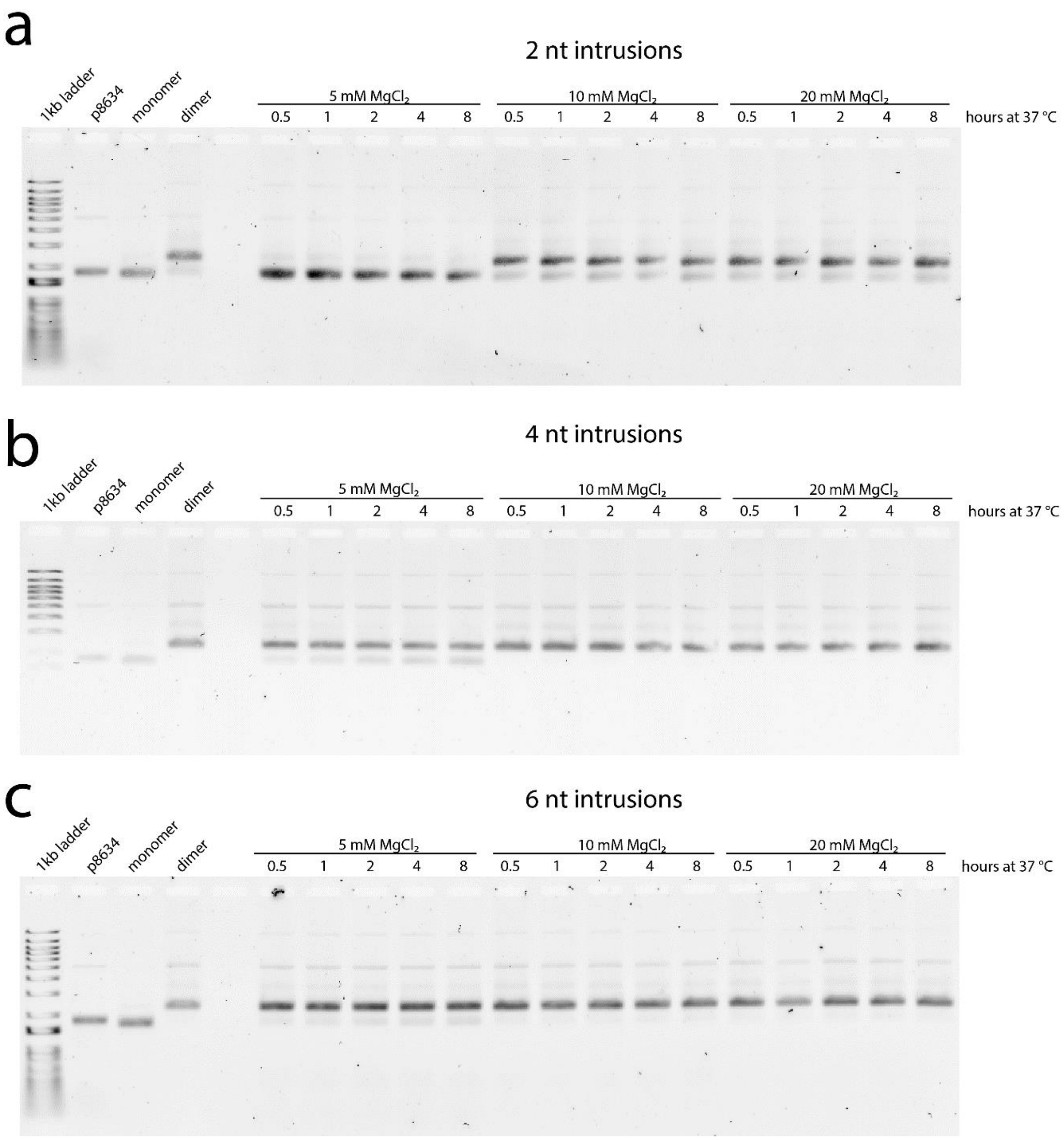

**Figure S43: AGE shift assay of xy-disassembly:** Dimers from xy-connections with (**a**) 2 nt, (**b**) 4 nt, and (**c**) 6 nt staple intrusions were formed over-night. They were diluted in buffers with a total $MgCl_2$ concentration of 5 mM, 10 mM, or 20 mM each, 8, 4, 2, 1, or 0.5 h before the samples were transferred to the gel. Data also used in figure 5.

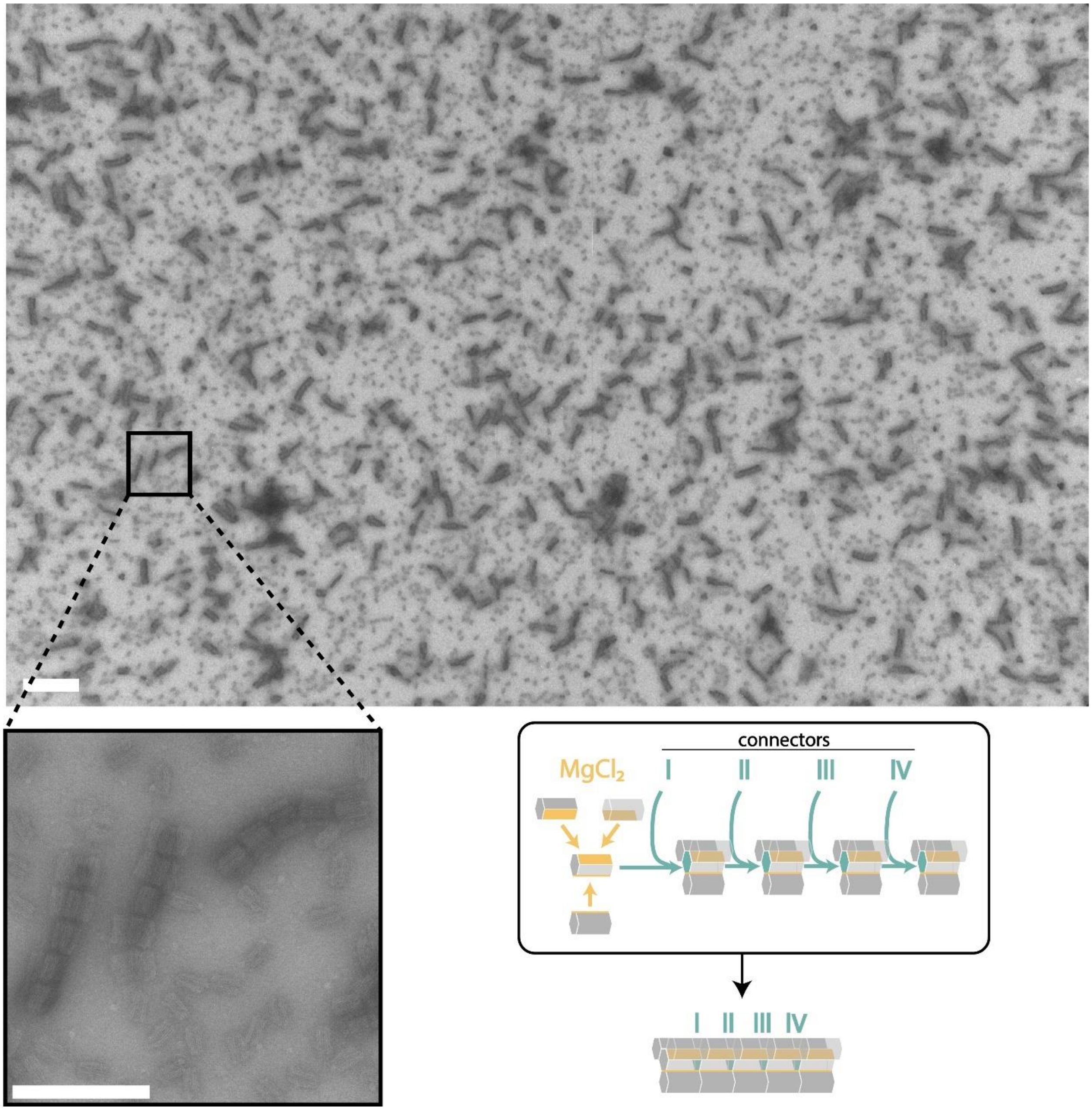

**Figure S44: TEM micrographs of tetrameric pentamer assembly** from xy- and z-connections. The central moDON monomers carry the same xy-connections: δεζ, connecting to moDONs with connection sites δ*, ε*, or ζ*, respectively. At the same time, the central moDONs have different z-connection handles, namely zI-right, zI-left/zII-right, zII-left/zIII-right, zIII-left/zIV-right, or zIV-left. Addition of respective connectors forms a pentamer of tetramers. Scale bar is 500 nm for the overview picture and 200 nm for the enlarged part.

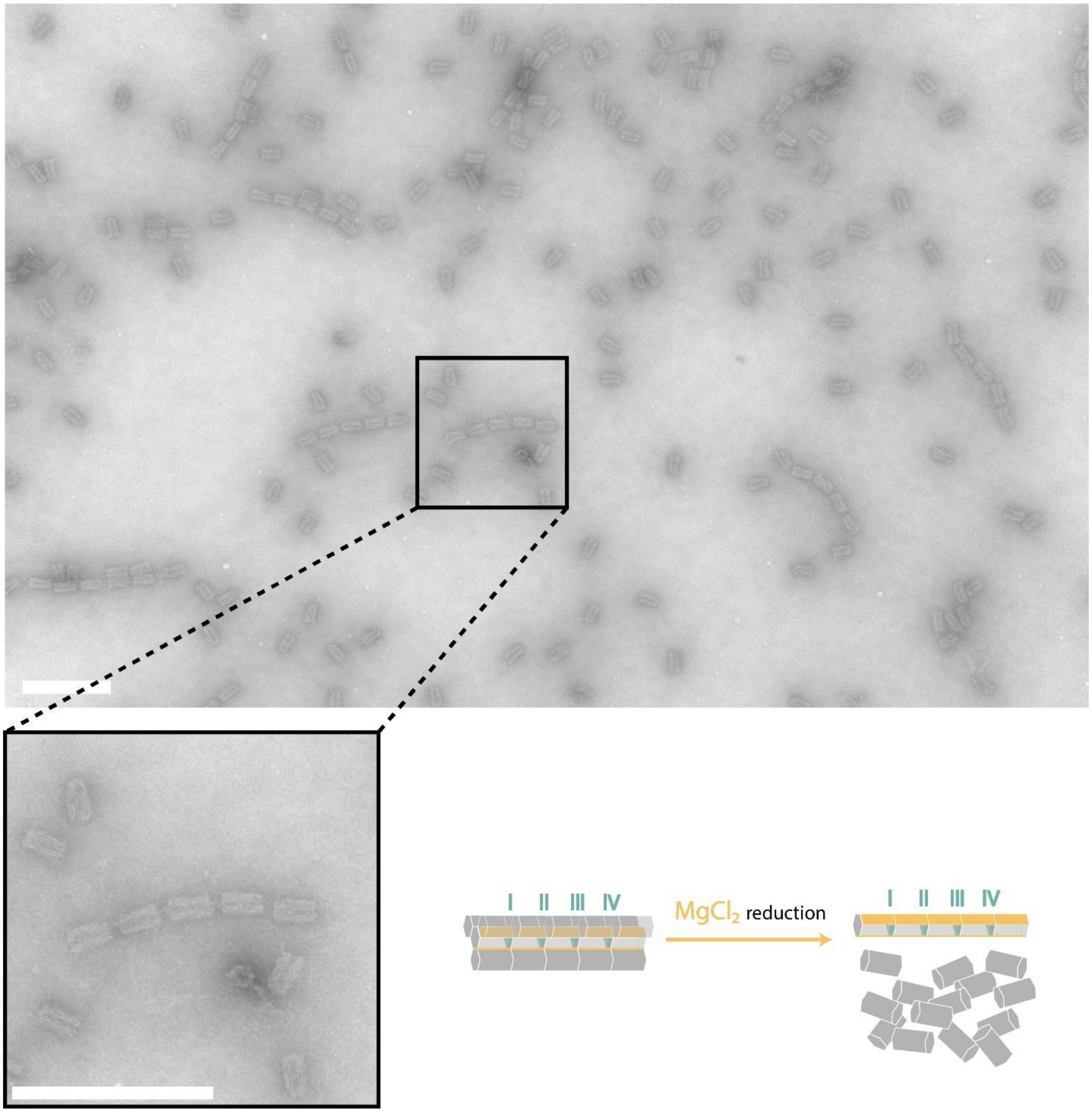

**Figure S45: TEM micrographs of tetrameric pentamer disassembly 1.** Reduction of $MgCl_2$ in the buffer solution leads to disassembly of xy-connections only, resulting in a z-pentamer and monomers. Scale bar is 500 nm for the overview picture and 200 nm for the enlarged part.

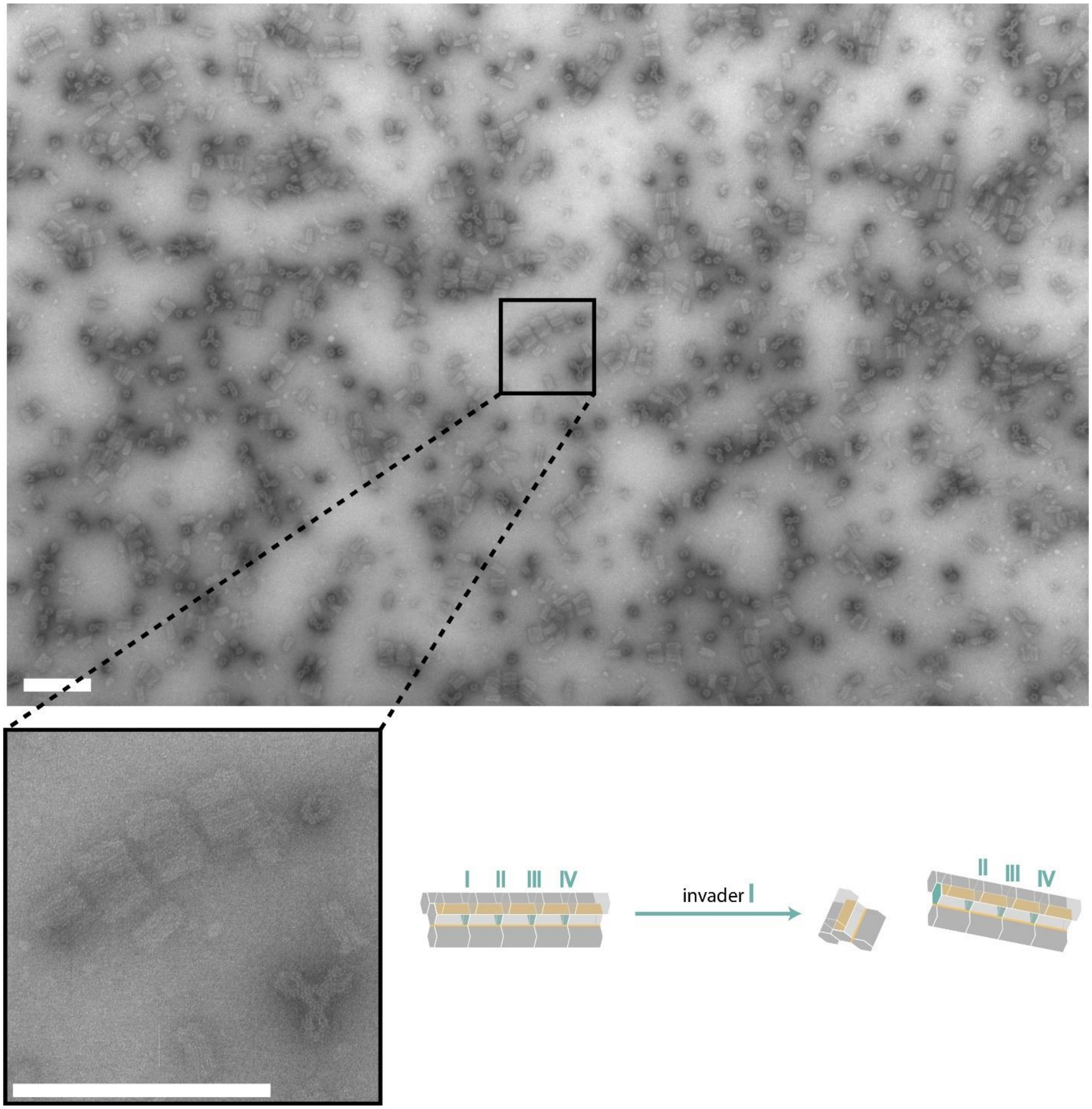

**Figure S46: TEM micrographs of tetrameric pentamer disassembly 2.** The addition of invader I leads to disassembly of the z-connection between the first and the second tetramer, resulting in a tetramer and a tetramer of tetramers. Scale bar is 500 nm for the overview picture and 200 nm for the enlarged part.

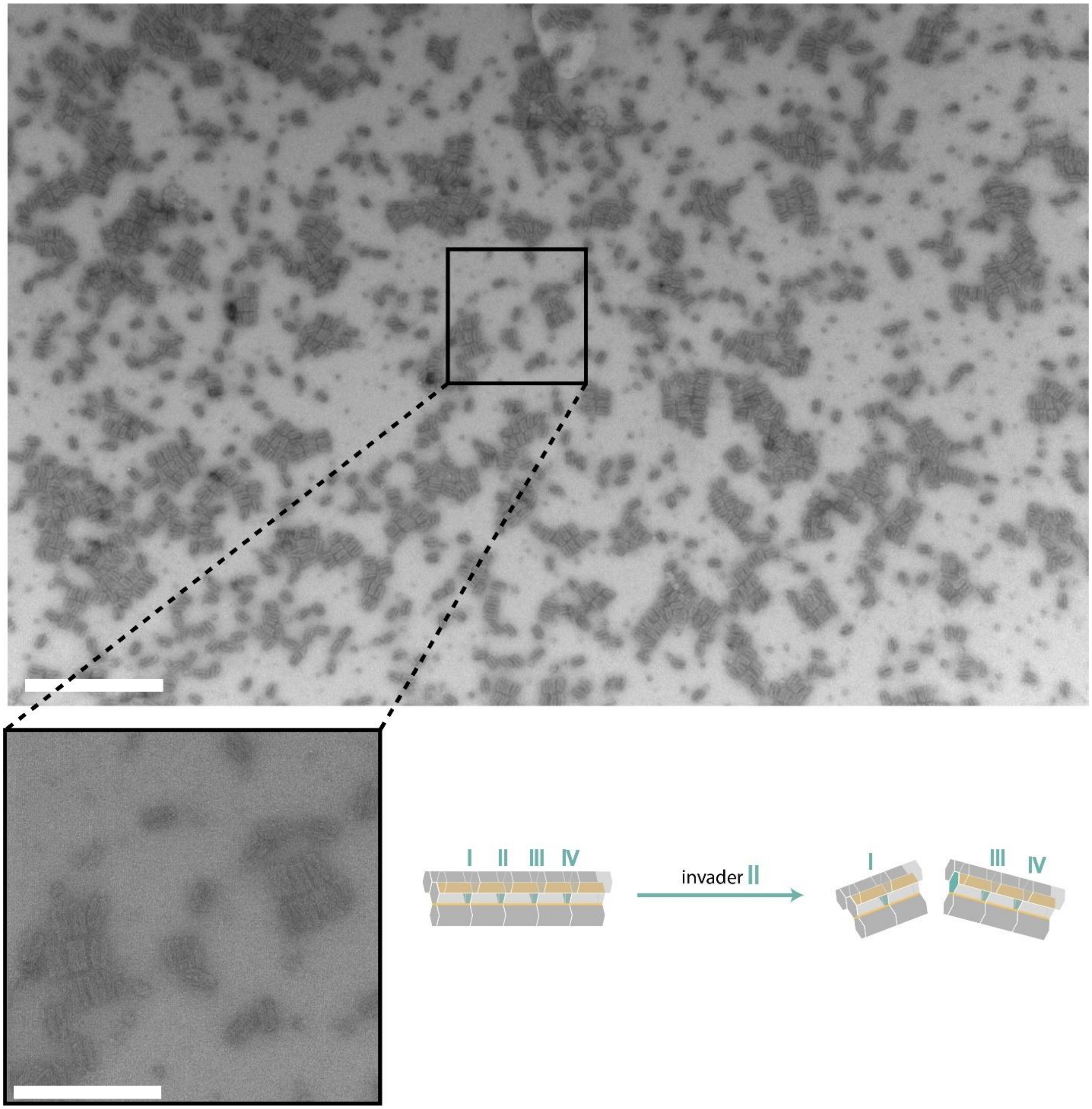

**Figure S47: TEM micrographs of tetrameric pentamer disassembly 3.** The addition of invader II leads to disassembly of the z-connection between the second and third tetramer resulting in a dimer and a trimer of tetramers. Scale bar is 500 nm for the overview picture and 200 nm for the enlarged part.

**Table S1: Staple connection sites and the respective staple mixes**: Configuration 1 and configuration 2 differ only in one half of the total connection site. Each belongs to a different hinge helix. Since the distribution of changed helices is not point-, but mirror-symmetrically (cf. Figure S6), HH50 has four different conformations, HH65, on the opposite side has only one, and the others have two each. This results in the use of the following staples mixes for the modular parts, ordered by hinge helix. Similar colors indicate the same staple mix.

| hinge helix | connection site | staple mix |
|---|---|---|
| 50 | γ* α | γ* α |
| 50 | γ* δ | γ* δ |
| 50 | ζ* α | ζ* α |
| 50 | ζ* δ | ζ* δ |
| 55 | α β* | α β* |
| 55 | α ε* | δ ε* |
| 55 | δ β* | α β* |
| 55 | δ ε* | δ ε* |
| 60 | β* γ | β* γ |
| 60 | β* ζ | ε* ζ |
| 60 | ε* γ | β* γ |
| 60 | ε* ζ | ε* ζ |
| 65 | γ α* | γ α* |
| 65 | γ δ* | γ α* |
| 65 | ζ α* | γ α* |
| 65 | ζ δ* | γ α* |
| 70 | α* β | α* β |
| 70 | α* ε | α* β |
| 70 | δ* β | δ* ε |
| 70 | δ* ε | δ* ε |
| 75 | β γ* | β γ* |
| 75 | β ζ* | β γ* |
| 75 | ε γ* | ε ζ* |
| 75 | ε ζ* | ε ζ* |

**Table S2: Core staples** are constant for all configurations.

| name | sequence 5* -> 3* |
|---|---|
| core_001 | GGAGAATGGATCCCGCCAGTGTGTGCTG |
| core_002 | TAAATGCATCCTCGGAGAAATGACTGATACCGTGAATATTA |
| core_003 | GGTTTTGTATTTTATCTGAACTCTTTTT |
| core_004 | CATAACAACTTCCGTCGCATTCACCCTCATTCAG |
| core_005 | TTTTTCCGCTCACAATTGAGTGAGCTAACTCACATTTTTTT |
| core_006 | TAAACCCCAAAATTATTATCAGGCCAACGGATTTA |
| core_007 | TCGACATTACTTCTAATAACATCACTTGATCTCGG |
| core_008 | GTAACTAGTTTTGTAAAAGATCTTCACAGAGTCTG |
| core_009 | AATCAGATATAATCCAATATTACCGCCATCGTCTG |
| core_010 | AGGATGCAGGTTAGCCTCGTGTATTAA |
| core_011 | AGGTCAAGATGTCTGACGCTGGTAGCGG |
| core_012 | GTAGCAACATTACGCATCGCTATTACGGGCAAATTAGAAGAA |
| core_013 | CTGAGAAGTGTTTTGCGGGAGCTAAACAGGAGGCC |
| core_014 | GTAAAAGCTTGCTGGACAGTCAAATCACATTTGGG |
| core_015 | GGAAGACATTGCTAAACTGGAATACATCGTACCCC |
| core_016 | ATGGGAGAGGAGAACGAGGATATTGCGCAGGTGTTCCTGAGTAACCGTT |
| core_017 | AAATGGATGGCAGAACAAAATAAACAGC |
| core_018 | GAGCAGCTATCGGCAGTCTGTCCATCACGGTTGG |
| core_019 | TTACGATAACAGTATCGATTAGTTGCTATTTTGCGCGAGGCAAAAA |
| core_020 | GCGACCTAAGCGTTCTTAGTTTGACTGTTATCAAGCACTGCATCCTG |
| core_021 | TTTTTAAGAAACCAGCAAAGCAAC |
| core_022 | TGAAACATCTGACCAATACCGAACGAAC |
| core_023 | TTTTTTAAAAGGGACATTCTGGTCACACCGCTCAAGCCATTG |
| core_024 | GCCAACAAGAAGATGAGAGCCTGCTGAA |
| core_025 | CACCAGCGAGATAGAACCCTGCTACATTTATTAACCAGAA |
| core_026 | TGATAGCCACAGACAATATTTTTGAATGTTTAATGCGAAGTGATGAACG |
| core_027 | CTCAGGCACTGCGTGATGCAACTTTTTC |
| core_028 | AACAGTGGCATCTGCCTTTTT |
| core_029 | AAGACGCACTAATAGATTAGAGATAATATTATTATAGTCACA |
| core_030 | TGCCACTCATTGTTGTGAGTGTGGCGATAGAAATA |
| core_031 | ACTTGTGGGAGGATTGGGATAGGTCACGATGAGAA |
| core_032 | GAAGAATATCATTGATGCGTATTAACCATTTAACA |
| core_033 | GCAAATCAACAGTTGAAAGGAATCACCTAGCAGCAAGATGGG |
| core_034 | AAGGTTAAATTCGACAACTCGGGGAAACTTCACCGGTTCCG |
| core_035 | CTAACACTGGTCGTAAACAGAGAGGTTCGCGAAC |
| core_036 | TTTTTCAAATCCCCACCGAACTGTTTTT |
| core_037 | CACAATAGCCGTTCCCGATAGAGCGAAATTAACT |
| core_038 | TTTTTTTAATGAAAGATTAATGAAGATTTTT |
| core_039 | GAAGTATAACGACGCGGGTAC |
| core_040 | TTTTTTTTAAAAGTTTGAGTACCCGAACCTCACTG |
| core_041 | CATTTTGGAAACCACGCCATTAGCCAGCTCGGCCTCAGGAAG |
| core_042 | TTTTTAAATTAATTACAACAGTTCAGGGATTTTT |
| core_043 | CAGTACATCTGTAAAGGTTGGGTTATATAACTATTTTT |
| core_044 | TTTTTTGCCGGAAACCACCTCAAA |
| core_045 | CGCAACTGATTAAGGAGTCAATAGTGAATTTATCAAAATCA |
| core_046 | ATCGTCGCTATTAATTAATTTTACCTTT |
| core_047 | TTTTTAGTGAATAACCTTGCTTAAATCA |
| core_048 | ACGCTCGCACTCCCGCCATTTAAAGCATTGAGG |
| core_049 | TTTTTAGTTTGAGGGGACGACTAACCGTCCACGCTAAAACAG |
| core_050 | TTTTTTATGTAAATGCTGAAGCGGGACGCAGACCTTTTAACTGCT |
| core_051 | CGCATCGGACAGTATTTCCGGCACCGCTTCTGGTTTTT |
| core_052 | TTGGTGTAATGAAAAATGCCACAAGTTCCAGGCTG |
| core_053 | CGGATTGACCGTAACTGGAACTTATCAGTTCAGGGAGGGCGACAAGGC |

| | |
|---|---|
| core_054 | AAATATATTTTAGTTATTTTT |
| core_055 | TTTTTCTTATCATTCCATTTATTTTCATCGTAGGAATTTTT |
| core_056 | CCAGCTTTAATTCGGTAAGAATACGTGGCCTAAAACATCGCCATTAAA |
| core_057 | AAATCAGATCGATTGTGCTGGCCATGAA |
| core_058 | TTTTTAAAATTCGCGGATGAACGGGATTTTT |
| core_059 | AGTTAAAATATGTTTTGAAGCCTTAAATCCCGACTTTAACCGAGTAAC |
| core_060 | CCAATAGGAGCGTCTTTCCTTTTT |
| core_061 | GTCCATGATATTATTTGTGCACATAAACATTGCTAAGAAAG |
| core_062 | ATAAGTCGGCAGACTACAGCGCAACACA |
| core_063 | TTTTTAATTGCGTTGCGGTTATTAATTTTTT |
| core_064 | TGCCAGCTACAAACTCTAAAATATCTTTAGGAGCA |
| core_065 | TGCATTAGACGGGCAACAGCTGATTGCCCCTGTCG |
| core_066 | GCGCGGGGTTTTTCAAAGATTGGGCGTTATCAATGTGGGCGC |
| core_067 | TTTTTAGCCCCAAAAACAGGAGGTTGATAATCAGAAATTTTT |
| core_068 | TTTATCCTGAATCTTACCAA |
| core_069 | AATCGTAAAACTAGCATGTCAGAGCCG |
| core_070 | TTTTTAGAGCCTAATTTGCCAGTTTTCACCA |
| core_071 | TTTTTTCATATGGTAACCGATTGAGGTTTTT |
| core_072 | CAAAGGCCAAGAGAAGGGAAACTGCGTG |
| core_073 | TTAATCATATTCATATAGCAGCACCGTGCGTCAG |
| core_074 | TTTTTAGTAATGTGTAGGTAATTAAATGCAATGCCTGTTTTT |
| core_075 | GGCCGGAGTAATATAATCAAAACTCAACTTGAGCT |
| core_076 | ATGATATTCAACCGCACCGTCACGTCAC |
| core_077 | TTTTTGAGGGAAGGTAAATATTGACGGAAATAAAGGGC |
| core_078 | TTTTTGCGGGAGAAGCCTTTATTTCAACGCAAAACATTATGACCC |
| core_079 | AGGATAAAACCCTCATATATTAGATTCAAAAGGGTTCCAAAT |
| core_080 | CACCAGTAGCTAAACCACCGA |
| core_081 | GCCGGAAACCGACTTGAGCCCATCAATCTCAAACGTGAGTA |
| core_082 | TTTTTATCAAGTTTCGGCATTTTCGGTTTTT |
| core_083 | TAGCGCGTCGCAAATGGTCAATAACCTGTTTAGC |
| core_084 | TTTTTGCCCGTATATGTAATACTTTTTTTTT |
| core_085 | TTTTTTTAGTTTGACCATTAGCTGCGAACGAGTAGATTTTTT |
| core_086 | GGGCGCGAGCTGAACGAGAGG |
| core_087 | CCCAATTATACATTTTTTCATGCCTTTAAATCAGTAGCGACAGATTTTT |
| core_088 | TAAGTATTCATTTGCTAATAGTAGTAGCATTAACATCCAAT |
| core_089 | CAGGGTGGAGAGGCGGTTTGC |
| core_090 | AGATATAGAGTCGGCATACAAAAGGTTT |
| core_091 | GAGTTGCGCGAAAAAATAGCCCGAGATATCCACTA |
| core_092 | CAGTGAATGAATCCGAGTACACATATAGATGAT |
| core_093 | CTCAGAGGCTCAGTGAGGCTGAGACTCCGTATAACAATGCGC |
| core_094 | GAGCTTAATTGCTGAGGAGCGGAGATCG |
| core_095 | TTTTTAGCAAGCCCGGGCGCGTACTATTTTT |
| core_096 | AACTACAGAAGCAAGACCATATTGAATCCCCCTCAGATAGCG |
| core_097 | TTTTTCATAATGCCTCGCCTGATAAATTTAGCCGGGTGTCTT |
| core_098 | TTTTTACCCTGACTATTATAGTCAACGCCTGTAGCATTTTT |
| core_099 | TTTTTAAACGGGTAAAATACGTGAGGAACTTACTGTAGTGTC |
| core_100 | ACACTAAAACACTCTAAGAGGAAGCCCGGATTAGA |
| core_101 | TTTTTCATAAATATTCAAATCAAAAATCAGGTCTTTTTTTT |
| core_102 | TTTTTCCTGCTCCATGTTACTGTGTCGAAATCCGCGATTTTT |
| core_103 | TAGACTGAATGCTCAGAAAACGAGAATAGCGGATTTATAAACTCCAAC |
| core_104 | AAAAACCAAACGGCGCAGAATGTATCAACTACG |
| core_105 | TTTTTTTCAACTAATGCAGATACACTGCGGAATCGTTTTTT |
| core_106 | AAAAGGACTGGGGTTCCAGTCTATTAAATCCTTTGACATTAT |
| core_107 | GTCTCCAGTATTATGTTTCCATTTTTTT |
| core_108 | TGATGGTGCCTGGCCCTGAGACCCGCTT |
| core_109 | AATGAGTTGCAAGGAGTTTATAAGGCAAAAATCATAAATGTT |
| core_110 | AATCCCTGCATCAAAAAGAT |

| | |
|---|---|
| core_111 | CGTGCCAGCGAAAAATATAATGCTGTCTTTGAG |
| core_112 | TTAAAGAACGTGGATCAAAAGTCCTGTT |
| core_113 | GTCAAAGCACCCGCCGCGCTTGTGCTTT |
| core_114 | CGAGAAAGGGCGCTTGGCTTAAGGTTTAGTACCGC |
| core_115 | GCGTCGTAACGCTTAACAAGACCCGTTA |
| core_116 | TTTTTCTGAATTTCCATGTTTTAAATTTTTT |
| core_117 | GCGCGTAACCACCAGGCGAAAAACCGTCTATCAAGCCGGCGA |
| core_118 | TCACGCTATAAGAGGTCATTTAGGTCAGAAAGACT |
| core_119 | GGCAAGTGCATTCGCATCAAATATATTTAGCCCGGAATAGGTACTCAGG |
| core_120 | GGAGAGTGGCGCTAGGAAGGGGATACCGTTTAGCTGAAACGAACGTGG |
| core_121 | TTTTTTGGTTGCTTTGACGAGCACTCAAGAGAAGGATTTTT |
| core_122 | GAGTATCTGCATATTGGGTTC |
| core_123 | CAGAATCCAACAGGAAAAACGCTCATTTTTT |
| core_124 | AGGTGAGGCGGTCAGTATTTTTTT |
| core_125 | TTTTTTTTCTGCGGCAGTTAATCGGTGAAAATGTTTTT |
| core_126 | TTTTTGAAACCAGTTTCTTGTGAAGTCCGTGAAGACGTTTTT |
| core_127 | TTTTTTTTATGTAGATGAAGGTAT |

**Table S3: Staples for HH50 modular part in configuration 1:** Staples also used in other configurations are printed in red, and on the right the respective other configuration is marked with a "y".

| name | sequence 5* -> 3* | δεζ | γ*δ | ζ*α |
|---|---|---|---|---|
| **hh50_Gamma*Alpha_shell_01** | **CGAGCTCGTACAAAGGTGGAAACGATACTTAAAGTAGCATGC** | | **y** | |
| **hh50_Gamma*Alpha_shell_02** | **CCTCTCTGAATTCGTAATCATGGTCATAGCCGGAGTAAAGC** | | **y** | |
| **hh50_Gamma*Alpha_shell_03** | **GCCTAATCCACACATAACGGAACAACATTATTATTTTT** | **y** | **y** | **y** |
| **hh50_Gamma*Alpha_shell_04** | **ATCAGTTTTAAACTTTGACCCAATAGTAGAGTATC** | | | **y** |
| hh50_Gamma*Alpha_shell_05 | TGCAAAAGAAGTTTTGAGCAATTTTCAC | | | |
| **hh50_Gamma*Alpha_shell_06** | **TCCAATATAACGCCTTCAGTTTTCATATACCAGTC** | | **y** | |
| hh50_Gamma*Alpha_shell_07 | TCAATCACAAATCATGACAAGAACCGGA | | | |
| **hh50_Gamma*Alpha_shell_08** | **AGGCGCAGGGGATTTTTTATGGAGATGA** | **y** | **y** | **y** |
| hh50_Gamma*Alpha_shell_09 | TTTTTTGAACGGTGTACAGACCTATTGAAAGAGGACAGATTTTT | | | |
| **hh50_Gamma*Alpha_shell_10** | **AACGAACACATACGAGCTGTTTCCTGTG** | | **y** | **y** |
| **hh50_Gamma*Alpha_shell_11** | **TTTTTCAGGTAGAAAGAGAGATTTAGGAATACCACATTTTT** | **y** | | |
| hh50_Gamma*Alpha_shell_12 | ACCAACTTTCATTACCTAAGGGAATTCTGC | | | |
| hh50_Gamma*Alpha_shell_13 | CCTTATGTAAGAGCAACACTAAGGGGGTCCAGCGA | | | |
| hh50_Gamma*Alpha_shell_14 | TTTCTTTAGATCCGAACGAGG | | | |
| hh50_Gamma*Alpha_shell_15 | CATACATTAGAGTCTGCCAGTCATAACATCATTGTGAATTA | | | |
| hh50_Gamma*Alpha_passive_Gamma*_01 | AATGTGCCACTCGCTAGGCTGGCTGACCTTCATTTTTT | | | |
| **hh50_Gamma*Alpha_passive_Gamma*_02** | **ATTACGAGGCATAGCGATTTTTGGGAAGAATTTTT** | | **y** | |
| **hh50_Gamma*Alpha_passive_Gamma*_03** | **TTTTTAAATCTACGTTAATAAAGGACGTAAGAACTGGCTCA** | | **y** | |
| hh50_Gamma*Alpha_passive_Gamma*_04 | TTTTTCAAGAGTAATCTACGTAACAAAGCTGCTCATTC | | | |
| hh50_Gamma*Alpha_2nt_intrusion_Gamma*_01 | AATGTGCCACTCGCTAGGCTGGCTGACCTTCATGC | | | |
| **hh50_Gamma*Alpha_2nt_intrusion_Gamma*_02** | **ATTACGAGGCATAGCGATTTTTGGGAAGAAGC** | | **y** | |
| **hh50_Gamma*Alpha_2nt_intrusion_Gamma*_03** | **CCCAAGAGTAATCTACGTAACAAAGCTGCTCATTC** | | **y** | |
| hh50_Gamma*Alpha_2nt_intrusion_Gamma*_04 | TGAAATCTACGTTAATAAAGGACGTAAGAACTGGCTCA | | | |
| hh50_Gamma*Alpha_passive_Alpha_01 | GTACAACACCAGAAAATAAGGCTTGCCCTGTTTTT | | | |
| hh50_Gamma*Alpha_passive_Alpha_02 | AGATTTCGTTTATGCGAGTAGTAAATTGGGTTTTT | | | |
| hh50_Gamma*Alpha_passive_Alpha_03 | TTTTTCTTGAGATGAACTTTACCTCGTTTACCAGA | | | |
| hh50_Gamma*Alpha_passive_Alpha_04 | TTTTTACGAGAAACGGAGATTTAGCGAGAGGCTTTCGACGAT | | | |
| hh50_Gamma*Alpha_2nt_intrusion_Alpha_01 | GTACAACACCAGAAAATAAGGCTTGCCCTGTA | | | |
| hh50_Gamma*Alpha_2nt_intrusion_Alpha_02 | AGATTTCGTTTATGCGAGTAGTAAATTGGGTG | | | |
| hh50_Gamma*Alpha_2nt_intrusion_Alpha_03 | AAACGAGAAACGGAGATTTAGCGAGAGGCTTTCGACGAT | | | |
| hh50_Gamma*Alpha_2nt_intrusion_Alpha_04 | CCCTTGAGATGAACTTTACCTCGTTTACCAGA | | | |

**Table S4: Staples for HH55 modular part in configuration 1:** Staples also used in other configurations are printed in red, and to the right the other configuration is marked with a “y”.

| name | sequence 5* -> 3* | δεζ |
|---|---|---|
| hh55_AlphaBeta*_shell_01 | ACCAGGCGGATAATCAGAACGTTTGCTTTTAATT | |
| **hh55_AlphaBeta*_shell_02** | **TTTTTATGCAACTAAAGAGGCCGCTTTTTTT** | **y** |
| hh55_AlphaBeta*_shell_03 | ACACTGACGCCACCTTCTGTATGGGATT | |
| **hh55_AlphaBeta*_shell_04** | **TTTTTGAGGGTAGCAACGGCTAAGACAGCATCGGAACTTTTT** | **y** |
| hh55_AlphaBeta*_shell_05 | CGTCACGAATAATAGAAAGGAACAACTATGAATT | |
| hh55_AlphaBeta*_shell_06 | ATCGCGTAAGCAAACCGACAA | |
| hh55_AlphaBeta*_shell_07 | AAAGAATTTATACCAAGCGCGAAACAAAAAACGAAAGCTTGC | |
| hh55_AlphaBeta*_shell_08 | CGATAGTTGGGCTCAAAACGCTCCAACTTGCGGA | |
| **hh55_AlphaBeta*_shell_09** | **GCAGCGAACAGAGGAGCTCAA** | **y** |
| **hh55_AlphaBeta*_shell_10** | **TTTTTTTCCACAGACAGTAGCGTAACGATCTAAAGTTTTTT** | **y** |
| **hh55_AlphaBeta*_shell_11** | **TTTTTTTGCGGGATCGTCACCGAGTTAATACGGTG** | **y** |
| hh55_AlphaBeta*_shell_12 | AAATAGTCCCTCAGGAATTGCCAGTACATACCGTA | |
| hh55_AlphaBeta*_shell_13 | TGAGAATAATTTTTTCACGTTGAGTACCCCTTTTG | |
| **hh55_AlphaBeta*_shell_14** | **TTTTTTTTGTCGTCTTTTTCAGGGATTTTTT** | **y** |
| **hh55_AlphaBeta*_shell_15** | **CGCTACAAATAGGACCTCATTCCAGACGTTAGTAA** | **y** |
| hh55_AlphaBeta*_passive_Alpha_01 | TTTTTTTAATTGTATCGGTTTATCAGAGGCATCAAAT | |
| hh55_AlphaBeta*_passive_Alpha_02 | TTTTTTTAAACAGCAACCATCGCGAACCAGACCGGTTTAATTCAACCTA | |
| hh55_AlphaBeta*_passive_Alpha_03 | TGACAACTTGATACTTTCGAGGTGAATTTCTTTTT | |
| hh55_AlphaBeta*_passive_Alpha_04 | GAAAATCTCCAAAACAAAAGGAGCCTTTTTT | |
| hh55_AlphaBeta*_2nt_intrusion_Alpha_01 | TGACAACTTGATACTTTCGAGGTGAATTTCCA | |
| hh55_AlphaBeta*_2nt_intrusion_Alpha_02 | GAAAATCTCCAAAACAAAAGGAGCCTCG | |
| hh55_AlphaBeta*_2nt_intrusion_Alpha_03 | AATTAAACAGCAACCATCGCGAACCAGACCGGTTTAATTCAACCTA | |
| hh55_AlphaBeta*_2nt_intrusion_Alpha_04 | AGTTAATTGTATCGGTTTATCAGAGGCATCAAAT | |
| hh55_AlphaBeta*_passive_Beta_01 | GTTGATACACCCTCAGAACCACAACTTTCTTTTT | |
| hh55_AlphaBeta*_passive_Beta_02 | AAGGCACCGAGCTTCAAAGACGACTAAAGCCCACGATATTCGGTTTTTT | |
| hh55_AlphaBeta*_passive_Beta_03 | TTTTTAACAGTTTCAGCGGAGTTGCTAAGCCACCCGTGCCGT | |
| hh55_AlphaBeta*_passive_Beta_04 | TTTTTCGCTGAGGCTTGCAGGCTCCGATCATAACA | |
| hh55_AlphaBeta*_2nt_intrusion_Beta_01 | GTTGATACACCCTCAGAACCACAACTT | |
| hh55_AlphaBeta*_2nt_intrusion_Beta_02 | AAGGCACCGAGCTTCAAAGACGACTAAAGCCCACGATATTCGGTTC | |
| hh55_AlphaBeta*_2nt_intrusion_Beta_03 | CACGCTGAGGCTTGCAGGCTCCGATCATAACA | |
| hh55_AlphaBeta*_2nt_intrusion_Beta_04 | TCAACAGTTTCAGCGGAGTTGCTAAGCCACCCGTGCCGT | |

**Table S5: Staples for HH60 modular part in configuration 1:** Staples also used in other configurations are printed in red, and to the right the other configuration is marked with a "y".

| **name** | **sequence 5* -> 3*** | **δεζ** |
|---|---|---|
| **hh60_Beta*Gamma_shell_01** | **AGTGAGGTCGGTTATTTCGGTCTGAATTTACCGT** | **y** |
| **hh60_Beta*Gamma_shell_02** | **ACATACAAAATCTGTCAGAGGCCGCCACCCTCAG** | **y** |
| hh60_Beta*Gamma_shell_03 | AATGAAATATTCGGTGGCATCGCCAGAA | |
| **hh60_Beta*Gamma_shell_04** | **TTTTTTTAGGATTAGCGGGAGTGTACTGGTAATAAGTTTTT** | **y** |
| **hh60_Beta*Gamma_shell_05** | **CCGGAACGTTGATTTCTGGAAGTTTCATTCACCGTGTATCCA** | **y** |
| **hh60_Beta*Gamma_shell_06** | **TTTTTTTCATAATCAAAATCAAAGCGTTTATTACCGCCACCC** | **y** |
| **hh60_Beta*Gamma_shell_07** | **TTTTTTTTTAACGGGGTAGTAACAGTTTTTT** | **y** |
| hh60_Beta*Gamma_shell_08 | GCCGCCACCATCCTAATAAAGGGAGGTT | |
| **hh60_Beta*Gamma_shell_09** | **TTTTTTCATAGCCCCCTTGCCATCTTTTTTT** | **y** |
| **hh60_Beta*Gamma_shell_10** | **TCCAGTAAATGCCCCCTGCCTGTACCAA** | **y** |
| hh60_Beta*Gamma_shell_11 | TGGAAAGCGCAGTCAACCTATCATGAAA | |
| hh60_Beta*Gamma_shell_12 | GAGGCAGTAGCAAGCAATAAAGCCTCAGAGCATAA | |
| **hh60_Beta*Gamma_passive_Beta*_01** | **AACAGTTAGCCTTGCAGTGCGTCATACATGGCTTTTTT** | **y** |
| **hh60_Beta*Gamma_passive_Beta*_02** | **TATAACACAGAGCCACCACCGGAACCTTTTT** | **y** |
| **hh60_Beta*Gamma_passive_Beta*_03** | **TTTTTTTTGATGATACAGGGTTTTCCACCACACCCATG** | **y** |
| hh60_Beta*Gamma_passive_Beta*_04 | TTTTTGCCTCCCTCAGACCACCACCCTCAGA | |
| **hh60_Beta*Gamma_2nt_intrusion_Beta*_01** | **AACAGTTAGCCTTGCAGTGCGTCATACATGG** | **y** |
| **hh60_Beta*Gamma_2nt_intrusion_Beta*_02** | **TATAACACAGAGCCACCACCGGAA** | **y** |
| **hh60_Beta*Gamma_2nt_intrusion_Beta*_03** | **TGATGATACAGGGTTTTCCACCACACCCATG** | **y** |
| hh60_Beta*Gamma_2nt_intrusion_Beta*_04 | CTCCCTCAGACCACCACCCTCAGA | |
| hh60_Beta*Gamma_passive_Gamma_01 | TTACCATGTCAGACGATTGTTTTT | |
| hh60_Beta*Gamma_passive_Gamma_02 | TTTTTGCCTTGATATTCACAAACACATTAAAAATTCTA | |
| hh60_Beta*Gamma_passive_Gamma_03 | ATTGACAAACCACCACCAGAGCCTTTTT | |
| hh60_Beta*Gamma_passive_Gamma_04 | TTTTTGCCGCCAGCCAATGAACAAAGAATTAGCAAAATTAAG | |
| hh60_Beta*Gamma_2nt_intrusion_Gamma_01 | TTACCATGTCAGACGAT | |
| hh60_Beta*Gamma_2nt_intrusion_Gamma_02 | CTTGATATTCACAAACACATTAAAAATTCTA | |
| hh60_Beta*Gamma_2nt_intrusion_Gamma_03 | ATTGACAAACCACCACCAGAG | |
| hh60_Beta*Gamma_2nt_intrusion_Gamma_04 | CGCCAGCCAATGAACAAAGAATTAGCAAAATTAAG | |

**Table S6: Staples for HH65 modular part in configuration 1:** This part is the same for both configurations and not changed. All staples are printed in red, and to the right the other configuration is marked with a "y".

| name | sequence 5* -> 3* | δεζ |
|---|---|---|
| hh65_GammaAlpha*_shell_01 | AGATTGTACGCAAAGACACCACGGAATTTTT | y |
| hh65_GammaAlpha*_shell_02 | GAGCAAATATCAGGTCATTGCAATAAGA | y |
| hh65_GammaAlpha*_shell_03 | TTTTGAGAGATCTAAATAGCA | y |
| hh65_GammaAlpha*_shell_04 | GAATTATTTCTAGCTGATAAAGAGAGGGTTAGCAAA | y |
| hh65_GammaAlpha*_shell_05 | GAGGAAACTATCTTACCGAAGACAATGA | y |
| hh65_GammaAlpha*_shell_06 | CGTAGAAAATACACGCCAAAATCATATAAACGCC | y |
| hh65_GammaAlpha*_shell_07 | TTTTTAAAATGAAAATAGGGAAGCGCTTTTT | y |
| hh65_GammaAlpha*_shell_08 | TTACAGAAAGAAACGATTTTTTGTTTAACGTCATTTTT | y |
| hh65_GammaAlpha*_shell_09 | GAAATTGACAAGGAATAACATAACTGAACGCTAACCAAGCAAAGCCGTT | y |
| hh65_GammaAlpha*_shell_10 | GCCCAATCTTATTTATCCCAAGAGAAA | y |
| hh65_GammaAlpha*_shell_11 | GCAAGAACCCTTTTACCAGAAGGAAACC | y |
| hh65_GammaAlpha*_shell_12 | ATTTACGCCTTAAGCGCAATAATAACGG | y |
| hh65_GammaAlpha*_shell_13 | TTTTTTAAGTTTATTTTATAGAAAATTTTTT | y |
| hh65_GammaAlpha*_shell_14 | TTTTTATTAGACGGGAGAATTAAAAACAGCAGCCT | y |
| hh65_GammaAlpha*_shell_15 | CACCCTGAACAAAGATAACCCAGTTAA | y |
| hh65_GammaAlpha*_passive_Gamma_01 | AGCACCAAATTAGACAAAGTTTAAGAAAAGTAAGCAGTTTTT | y |
| hh65_GammaAlpha*_passive_Gamma_02 | GGCAAGGACCATCGTAAAGGTAATACCCAAAAGTTTTT | y |
| hh65_GammaAlpha*_passive_Gamma_03 | TTTTTATAGCCGAAGCCAGCAAATTTAGAAATTAT | y |
| hh65_GammaAlpha*_passive_Gamma_04 | TTTTTAACTGGCATGATTAAGACTCAGTATGTAGCTAT | y |
| hh65_GammaAlpha*_2nt_intrusion_Gamma_01 | AGCACCAAATTAGACAAAGTTTAAGAAAAGTAAGCAGCT | y |
| hh65_GammaAlpha*_2nt_intrusion_Gamma_02 | GGCAAGGACCATCGTAAAGGTAATACCCAAAAGAG | y |
| hh65_GammaAlpha*_2nt_intrusion_Gamma_03 | AGAACTGGCATGATTAAGACTCAGTATGTAGCTAT | y |
| hh65_GammaAlpha*_2nt_intrusion_Gamma_04 | TTATAGCCGAAGCCAGCAAATTTAGAAATTAT | y |
| hh65_GammaAlpha*_passive_Alpha*_01 | CATATGAGAGTCTGCTACAATGTAATTGAGTTTTT | y |
| hh65_GammaAlpha*_passive_Alpha*_02 | GACATTCTTACCAGTAAATCAGTCACCATAAAGGTGGCAATTTTT | y |
| hh65_GammaAlpha*_passive_Alpha*_03 | TTTTTCGCTAATATCAGAGAGTCAGAGG | y |
| hh65_GammaAlpha*_passive_Alpha*_04 | TTTTTCATATAAAAGAAATAAGCAATTGTAATTTTGTT | y |
| hh65_GammaAlpha*_2nt_intrusion_Alpha*_01 | CATATGAGAGTCTGCTACAATGTAATTG | y |
| hh65_GammaAlpha*_2nt_intrusion_Alpha*_02 | GACATTCTTACCAGTAAATCAGTCACCATAAAGGTGGC | y |
| hh65_GammaAlpha*_2nt_intrusion_Alpha*_03 | CTAATATCAGAGAGTCAGAGG | y |
| hh65_GammaAlpha*_2nt_intrusion_Alpha*_04 | TATAAAAGAAATAAGCAATTGTAATTTTGTT | y |

**Table S7: Staples for HH70 modular part in configuration 1:** Staples also used in other configurations are printed in red, and to the right the other configuration is marked with a "y".

| name | sequence 5* -> 3* | δεζ |
|---|---|---|
| hh70_Alpha*Beta_shell_01 | TGAAAGCCGTCTGGCGGTATTATAGATAAGTCCTGCATGTTC | |
| hh70_Alpha*Beta_shell_02 | TTGGTAACATAGTCGCTATCCCTCATTTTTGCGGG | |
| hh70_Alpha*Beta_shell_03 | TTAACCTCGCAAGACGTCGGAACCCAAGCCTGTT | |
| hh70_Alpha*Beta_shell_04 | AGAACGGGTATTAAGTAATTCACGACAA | |
| hh70_Alpha*Beta_shell_05 | TCATCAACATTAAATTATACA | |
| hh70_Alpha*Beta_shell_06 | ATCGAGAACAAGCAATCAGATAAAATAA | |
| hh70_Alpha*Beta_shell_07 | ACGCCATCGTTTTAGCGAACCTCAAGATGAACGGT | |
| **hh70_Alpha*Beta_shell_08** | **TTTTTTCATTACCGCGCTTACGAGCATTTTT** | **y** |
| hh70_Alpha*Beta_shell_09 | TTTTTAATACCGACCGTGTATTTTGTTTTTTT | |
| hh70_Alpha*Beta_shell_10 | TATCCCATAAGACGTGTCCATCGGCTGTCTTTCTTTTT | |
| **hh70_Alpha*Beta_shell_11** | **TTTTTTGTAGAAACCAATCAATCCTAAT** | **y** |
| hh70_Alpha*Beta_shell_12 | TAAACAAAACAAGAATAGAAGGCTTATCCCCACTC | |
| hh70_Alpha*Beta_shell_13 | TAGTATCATATGCGTGTGAGCAATAGGA | |
| hh70_Alpha*Beta_shell_14 | AGAAAAAAGAACGCGAGAAAAATCCAATCCGGCTT | |
| hh70_Alpha*Beta_shell_15 | ATTTCATCTTCTGACCTAATCATCCGGAAATTTAATGGTTTGATTTTT | |
| hh70_Alpha*Beta_shell_16 | AATAAGAGTAATTGGCTTAATTGAGAA | |
| hh70_Alpha*Beta_shell_17 | TCGCCATGGCATTTTCGAGCCAGAATAT | |
| hh70_Alpha*Beta_passive_Alpha*_01 | CCTGTAGAAAGTACAGCTAATGCAGAACGCGCCTTTTT | |
| hh70_Alpha*Beta_passive_Alpha*_02 | ATTAAATACGTTAATGATAAATAAGGCGTTAAATTTTT | |
| hh70_Alpha*Beta_passive_Alpha*_03 | TTTTTTGTTTATCAACACTAAGAACACCCAG | |
| hh70_Alpha*Beta_passive_Alpha*_04 | TTTTTTAAGAATAAACAAATTACT | |
| hh70_Alpha*Beta_2nt_intrusion_Alpha*_01 | CCTGTAGAAAGTACAGCTAATGCAGAACGCG | |
| hh70_Alpha*Beta_2nt_intrusion_Alpha*_02 | ATTAAATACGTTAATGATAAATAAGGCGTTA | |
| hh70_Alpha*Beta_2nt_intrusion_Alpha*_03 | TTTATCAACACTAAGAACACCCAG | |
| hh70_Alpha*Beta_2nt_intrusion_Alpha*_04 | AGAATAAACAAATTACT | |
| hh70_Alpha*Beta_passive_Beta_01 | CGACAAAAGGCAGAATTTAACAACGCCAACTTTTT | |
| hh70_Alpha*Beta_passive_Beta_02 | TTTTTATGTAATTTAGGTAAAACCAAGTACCGTT | |
| hh70_Alpha*Beta_passive_Beta_03 | TTTTTCAACGCTCATAGGTCTGTGGGAACAAACGG | |
| hh70_Alpha*Beta_passive_Beta_04 | ACAGTAGCTTACCAGTATAAAGCTTTTT | |
| hh70_Alpha*Beta_2nt_intrusion_Beta_01 | CCCAACGCTCATAGGTCTGTGGGAACAAACGG | |
| hh70_Alpha*Beta_2nt_intrusion_Beta_02 | CTATGTAATTTAGGTAAAACCAAGTACCGTT | |
| hh70_Alpha*Beta_2nt_intrusion_Beta_03 | CGACAAAAGGCAGAATTTAACAACGCCAACTT | |
| hh70_Alpha*Beta_2nt_intrusion_Beta_04 | ACAGTAGCTTACCAGTATAAAGCGC | |

**Table S8: Staples for HH75 modular part in configuration 1:** Staples also used in other configurations are printed in red, and to the right the other configuration is marked with a "y".

| name | sequence 5* -> 3* | δεζ |
|---|---|---|
| **hh75_BetaGamma*_shell_01** | **TTTTTGCTACGGCGCCTGAGCAATTTTT** | **y** |
| **hh75_BetaGamma*_shell_02** | **TCACGACTTGGGTAACGCCAGATTATTT** | **y** |
| **hh75_BetaGamma*_shell_03** | **TTCGCTATGGCGAACGGATTCGCCTGATTGCTTTTTGAATTACCCAGC** | **y** |
| **hh75_BetaGamma*_shell_04** | **GGCAAAGCCAGAAGGATAGAAAGGGTTGATGGCAATTCATTTTTT** | **y** |
| hh75_BetaGamma*_shell_05 | TCGGTGCTGAAAACATAGCGATAGCTTAGTTGGGATTGTCGG | |
| **hh75_BetaGamma*_shell_06** | **TTTTTATATTCCTGATTATCAGAGCGGAATTATCATCTTTTT** | **y** |
| **hh75_BetaGamma*_shell_07** | **TATCAAAGGACAAACGGATTTTCCCAG** | **y** |
| hh75_BetaGamma*_shell_08 | GCACGTAAAACAGAGATTAAGGTTGTAATAGACTT | |
| **hh75_BetaGamma*_shell_09** | **TTTTTAAGAAGATGATGAAACTCAATTA** | **y** |
| **hh75_BetaGamma*_shell_10** | **TTTTTCAATATAATCCTTGAAATTGTTATTTTTT** | **y** |
| hh75_BetaGamma*_passive_Beta_01 | ATTCTCCGAGAGACTCCCTTAGTACCTTTTACATTTTT | |
| hh75_BetaGamma*_passive_Beta_02 | TTTTTAGATGAATATACAGTAACAGAATCCTGGGCCTC | |
| hh75_BetaGamma*_passive_Beta_03 | TTTTTTCGGGAGAAACAATTTTTCGTAGAAAAGGGGGACCAAGCTCATTTGA | |
| hh75_BetaGamma*_passive_Beta_04 | AATAAAGAAATTGCAGGTTTAACGTCTTTTT | |
| hh75_BetaGamma*_2nt_intrusion_Beta_01 | ATTCTCCGAGAGACTCCCTTAGTACCTTTTA | |
| hh75_BetaGamma*_2nt_intrusion_Beta_02 | TCAGATGAATATACAGTAACAGAATCCTGGGCCTC | |
| hh75_BetaGamma*_2nt_intrusion_Beta_03 | GGGAGAAACAATTTTTCGTAGAAAAGGGGGACCAAGCTCATTTGA | |
| hh75_BetaGamma*_2nt_intrusion_Beta_04 | AATAAAGAAATTGCAGGTTTAACG | |
| **hh75_BetaGamma*_passive_Gamma*_01** | **TTACGCTTTTTTAATGGAAAATTTCATGAATACCATCGCGCAGTTTTT** | **y** |
| **hh75_BetaGamma*_passive_Gamma*_02** | **TTTTTCTGAATAATGGACCTACCA** | **y** |
| **hh75_BetaGamma*_passive_Gamma*_03** | **TTTTTAGGCGAATTATTCATTAAACAAAAAGTTACATCAAGAAAACATTTTT** | **y** |
| **hh75_BetaGamma*_passive_Gamma*_04** | **GATTGTTTGGATTATACTTTTTTT** | **y** |
| **hh75_BetaGamma*_2nt_intrusion_Gamma*_01** | **TTACGCTTTTTTAATGGAAAATTTCATGAATACCATCGCGC** | **y** |
| **hh75_BetaGamma*_2nt_intrusion_Gamma*_02** | **GAATAATGGACCTACCA** | **y** |
| **hh75_BetaGamma*_2nt_intrusion_Gamma*_03** | **GCGAATTATTCATTAAACAAAAAGTTACATCAAGAAAACATTTTT** | **y** |
| **hh75_BetaGamma*_2nt_intrusion_Gamma*_04** | **GATTGTTTGGATTATAC** | **y** |

**Table S9: Staples for HH50 modular part in configuration 2:** Staples also used in other configurations are printed in red, and to the right the other configuration is marked with a "y". Some staples from configuration 1 are also needed, *cf.* Table S3

| name | sequence 5* -> 3* | γ*δ | ζ*α |
|---|---|---|---|
| hh50_Zeta*Delta_shell_01 | TAATCATGGTCATAGCCGGAGTAAAGC | | |
| **hh50_Zeta*Delta_shell_02** | **CGAGCTCGTACAAAGGTGGAAACGATACTTAAAGTAGCATGCATCTACG** | | **y** |
| hh50_Zeta*Delta_shell_03 | GTACAACGGAGATTTAGCGAGAGGCTTTCGACGAT | | |
| hh50_Zeta*Delta_shell_04 | TCCAATATAACGCCCGCAGTTTTCATATTTTAAGA | | |
| hh50_Zeta*Delta_shell_05 | TACCAGATGCAAAAGAAGTTTTGAGCAATTTTCAC | | |
| hh50_Zeta*Delta_shell_06 | TTTTTGAACGGTGTACAGACCAGATTTGAAAGAGGACAGATTTTTT | | |
| **hh50_Zeta*Delta_shell_07** | **TTAATAAACTGGCTGAATTACCTTATG** | | **y** |
| hh50_Zeta*Delta_shell_08 | TATTCATTACCCAAATCAACGTAACAAAGCTGCTCCCTCGTT | | |
| hh50_Zeta*Delta_shell_09 | TTTTTACCAACTCTTGACAAGTAAGGGAATTCTGC | | |
| hh50_Zeta*Delta_shell_10 | GCTTGAATAAGAGCAACACTAAGGGGGTCCAGCGA | | |
| hh50_Zeta*Delta_shell_11 | TTTCTTTAGATCCGAACGAGGTCAATCAAACCGGA | | |
| hh50_Zeta*Delta_shell_12 | CATACATTAGAGTCTGCCAGTCATAACATTCAGTGAATAAG | | |
| hh50_Zeta*Delta_passive_Zeta*_01 | AATGTGCCACTCGCGGCTGGCTGTTTTT | | |
| hh50_Zeta*Delta_passive_Zeta*_02 | ATTACGAGGCATAGTCATTGTCATTATACCAGTCAGGACGTTTTT | | |
| hh50_Zeta*Delta_passive_Zeta*_03 | TTTTTTTGGGAAGAAAACCTCTCTGAATTCG | | |
| hh50_Zeta*Delta_passive_Zeta*_04 | TTTTTACCTTCATCAAGAGTAGCGCATAGGGGATTTTTTATGGAGATGA | | |
| hh50_Zeta*Delta_2nt_intrusion_Zeta*_01 | AATGTGCCACTCGCGGCTGGCTGAC | | |
| hh50_Zeta*Delta_2nt_intrusion_Zeta*_02 | ATTACGAGGCATAGTCATTGTCATTATACCAGTCAGGACGGT | | |
| hh50_Zeta*Delta_2nt_intrusion_Zeta*_03 | CCACCTTCATCAAGAGTAGCGCATAGGGGATTTTTTATGGAGATGA | | |
| hh50_Zeta*Delta_2nt_intrusion_Zeta*_04 | AGTTGGGAAGAAAACCTCTCTGAATTCG | | |
| hh50_Zeta*Delta_passive_Delta_01 | ATCAGTTTTAAACTGCTTGAGATGGTTTTTT | | |
| hh50_Zeta*Delta_passive_Delta_02 | TTTTTTTAATTTCAACTTTCCCTGACGAGAAACACCTTTTT | | |
| hh50_Zeta*Delta_passive_Delta_03 | TTTTTAGAACGAGTAGTAAATTGGTTGACCCAATAGTAGAGTATC | | |
| hh50_Zeta*Delta_2nt_intrusion_Delta_01 | ATCAGTTTTAAACTGCTTGAGATGGTGA | | |
| hh50_Zeta*Delta_2nt_intrusion_Delta_02 | TAAGAACGAGTAGTAAATTGGTTGACCCAATAGTAGAGTATC | | |
| hh50_Zeta*Delta_2nt_intrusion_Delta_03 | AATTAATTTCAACTTTCCCTGACGAGAAACACCGA | | |

**Table S10: Staples for HH55 modular part in configuration 2:** Some staples from configuration 1 are also needed, *cf.* Table S4

| **name** | **sequence 5* -> 3*** |
|---|---|
| hh55_DeltaEpsilon*_shell_01 | ACCAGGCGGATAATCAGAAC |
| hh55_DeltaEpsilon*_shell_02 | GTTGATACACCCTCAGAACCGCCACCCGTGCCGT |
| hh55_DeltaEpsilon*_shell_03 | GTTTGCTTTTAATTCGTCACTTCAGCG |
| hh55_DeltaEpsilon*_shell_04 | AAGGCACCGAGCTTCAAAGACGACTAAAAGCTTGATGCGCCG |
| hh55_DeltaEpsilon*_shell_05 | AAAGAATTTATACCAAGCGCGAAACAAAAAACGAAAAAAAAG |
| hh55_DeltaEpsilon*_shell_06 | ATCAGCTTGTGCGAGAATCTCTCCAACTTGCGGA |
| hh55_DeltaEpsilon*_shell_07 | GAGTGAGAATAGAAGAGTACCCCTTTTG |
| hh55_DeltaEpsilon*_shell_08 | ACAATGATCGGTCGCTGAGGCTTGCAGGCTATAGTTACCGCA |
| hh55_DeltaEpsilon*_shell_09 | CTATAGTCCCTCTTCAACAGTCAGTACATACCGTA |
| hh55_DeltaEpsilon*_passive_Epsilon_01 | TTTTTTCACGTTGAAAATCTCCAAAGAGGCATCAAAT |
| hh55_DeltaEpsilon*_passive_Epsilon_02 | TTTTTTTAATTGTACTTAAACGCGAACCAGACCGGTTTAATTCAACCTA |
| hh55_DeltaEpsilon*_passive_Epsilon_03 | TGAATTTTCGGTTTGCTCCAAAAGGAGCCTTTTTT |
| hh55_DeltaEpsilon*_passive_Epsilon_04 | AGGAACAACTAAAGATAATAATTTTTTTTTT |
| hh55_DeltaEpsilon*_2nt_intrusion_Epsilon_01 | TGAATTTTCGGTTTGCTCCAAAAGGAGCCTCA |
| hh55_DeltaEpsilon*_2nt_intrusion_Epsilon_02 | AGGAACAACTAAAGATAATAATTTTTCG |
| hh55_DeltaEpsilon*_2nt_intrusion_Epsilon_03 | AATTAATTGTACTTAAACGCGAACCAGACCGGTTTAATTCAACCTA |
| hh55_DeltaEpsilon*_2nt_intrusion_Epsilon_04 | AGTCACGTTGAAAATCTCCAAAGAGGCATCAAAT |
| hh55_DeltaEpsilon*_passive_Delta*_02 | ATCGCGTAAGCAAATTCGAGGCATCGCCCATTTTT |
| hh55_DeltaEpsilon*_passive_Delta*_03 | TTTTTGGATTTTGCTAAACAAATGAATT |
| hh55_DeltaEpsilon*_passive_Delta*_04 | TTTTTCGCATAACCGATATATCAACAAC |
| hh55_DeltaEpsilon*_2nt_intrusion_Delta*_01 | ACACTGACGCCACCTTCTGTATGTA |
| hh55_DeltaEpsilon*_2nt_intrusion_Delta*_02 | ATCGCGTAAGCAAATTCGAGGCATCGCC |
| hh55_DeltaEpsilon*_2nt_intrusion_Delta*_03 | ATTTTGCTAAACAAATGAATT |
| hh55_DeltaEpsilon*_2nt_intrusion_Delta*_04 | ATCGCATAACCGATATATCAACAAC |

**Table S11: Staples for HH60 modular part in configuration 2:** Some staples from configuration 1 are also needed, cf. Table S5

| name | sequence 5* -> 3* |
|---|---|
| hh60_Epsilon*Zeta_shell_01 | TTACCATTAGCAAGCAATAAAGCCTCAGAGCATAA |
| hh60_Epsilon*Zeta_shell_02 | AATGAAATATTCGGTGGCATC |
| hh60_Epsilon*Zeta_shell_03 | GCCAGAATGGAAAGCGCAGTCAACCTATCATGAAA |
| hh60_Epsilon*Zeta_passive_Epsilon*_04 | TTTTTGCCTCCCTCAGACCACCACCCTCACA |
| hh60_Epsilon*Zeta_2nt_intrusion_Epsilon*_04 | CTCCCTCAGACCACCACCCTCACA |
| hh60_Epsilon*Zeta_passive_Zeta_01 | AACAAATAAATCCTTTGGCCTCAGGAGGTTGAGGCAGTTTTT |
| hh60_Epsilon*Zeta_passive_Zeta_02 | TTTTTGTCAGACGACATTAAAAATTCTA |
| hh60_Epsilon*Zeta_passive_Zeta_03 | GCATTGATGATATTCAGAGCCGCCACCAGAACCTTTTT |
| hh60_Epsilon*Zeta_passive_Zeta_04 | TTTTTACCACCAGAGCCGCCGCCACAATGAACAAAGAATTAGCAAAATTAAG |
| hh60_Epsilon*Zeta_2nt_intrusion_Zeta_01 | AACAAATAAATCCTTTGGCCTCAGGAGGTTGAGGC |
| hh60_Epsilon*Zeta_2nt_intrusion_Zeta_02 | CAGACGACATTAAAAATTCTA |
| hh60_Epsilon*Zeta_2nt_intrusion_Zeta_03 | GCATTGATGATATTCAGAGCCGCCACCAGAA |
| hh60_Epsilon*Zeta_2nt_intrusion_Zeta_04 | CACCAGAGCCGCCGCCACAATGAACAAAGAATTAGCAAAATTAAG |

**Table S12: Staples for HH70 modular part in configuration 2:** Some staples from configuration 1 are also needed, *cf.* Table 7

| **name** | **sequence 5* -> 3*** |
|---|---|
| hh70_Delta*Epsilon_shell_01 | TGAAAGCCGTCTGGCGGTATTCTAAGAACACCCAG |
| hh70_Delta*Epsilon_shell_02 | TTGGTAACATAGTCGCTATCCCTCATTTTTGCGGGTAAGAAT |
| hh70_Delta*Epsilon_shell_03 | TTAACCTCGCAAGACGTCGGAACCCAAAGGGCTT |
| hh70_Delta*Epsilon_shell_04 | TCATCAACATTAAAATATTTA |
| hh70_Delta*Epsilon_shell_05 | ATTAAATACGTTAATGTGATAAATAAGGCGTTAAA |
| hh70_Delta*Epsilon_shell_06 | ACAAGCAATCAGATATAGAAGGCTTATCCCCACTCATCGAGA |
| hh70_Delta*Epsilon_shell_07 | AAACACCTATCATAAATTGAGAATCGCCTGTGAGCAATAGGA |
| hh70_Delta*Epsilon_shell_08 | TTTTTAAATACCGACCGTATTTTGTTTTTTT |
| hh70_Delta*Epsilon_shell_09 | TGCGTTATACAAATTCTTACCCAACGCT |
| hh70_Delta*Epsilon_shell_10 | CAACAGTAGAACGCGAGAAAAATCCAATCCGGCTT |
| hh70_Delta*Epsilon_shell_11 | TTTTTATTTCATCTTCTGACCTAAAAGCAGTATAATTTAATGGTTTGTTTTT |
| hh70_Delta*Epsilon_shell_12 | ATAGAATCGGCTGTCTTTCTTTTT |
| hh70_Delta*Epsilon_shell_13 | ACACATAAACAAAGAGCCAGTAATAAG |
| hh70_Delta*Epsilon_shell_14 | AGAATATCAGACGACGACAATGTTCAGC |
| hh70_Delta*Epsilon_passive_Delta*_01 | AGAACGGGTATTAAATCAACACCTGAACAATTTTT |
| hh70_Delta*Epsilon_passive_Delta*_02 | ACGCCATCGTTTTATAATTACTATTTTT |
| hh70_Delta*Epsilon_passive_Delta*_03 | TTTTTGAAAAATAATATCCCATATAAGT |
| hh70_Delta*Epsilon_passive_Delta*_04 | TTTTTGAAAAAGCCTGTTTAGGGAATCAGCGAACCTCAAGATGAACGGT |
| hh70_Delta*Epsilon_2nt_intrusion_Delta*_01 | AGAACGGGTATTAAATCAACACCTGAAC |
| hh70_Delta*Epsilon_2nt_intrusion_Delta*_02 | ACGCCATCGTTTTATAATTAC |
| hh70_Delta*Epsilon_2nt_intrusion_Delta*_03 | AAAATAATATCCCATATAAGT |
| hh70_Delta*Epsilon_2nt_intrusion_Delta*_04 | AAAAGCCTGTTTAGGGAATCAGCGAACCTCAAGATGAACGGT |
| hh70_Delta*Epsilon_passive_Epsilon_01 | GAACGCGTTCTGTCAAAGTACCGACAAAAGTTTTT |
| hh70_Delta*Epsilon_passive_Epsilon_02 | TTTTTGTAAAGTAACCTGTTTACCAAGTACCGTTCCTGTAGTAATGCA |
| hh70_Delta*Epsilon_passive_Epsilon_03 | TTTTTAGGCAGAGGTAGGTCTGTGGGAACAAACGG |
| hh70_Delta*Epsilon_passive_Epsilon_04 | CATTTTCCGCCAACATGTAATTTTTTTT |
| hh70_Delta*Epsilon_2nt_intrusion_Epsilon_01 | CCAGGCAGAGGTAGGTCTGTGGGAACAAACGG |
| hh70_Delta*Epsilon_2nt_intrusion_Epsilon_02 | CTGTAAAGTAACCTGTTTACCAAGTACCGTTCCTGTAGTAATGCA |
| hh70_Delta*Epsilon_2nt_intrusion_Epsilon_03 | GAACGCGTTCTGTCAAAGTACCGACAAAAGTT |
| hh70_Delta*Epsilon_2nt_intrusion_Epsilon_04 | CATTTTCCGCCAACATGTAATTTGC |

**Table S13: Staples for HH75 modular part in configuration 2:** Some staples from configuration 1 are also needed, *cf.* Table S8

| name | sequence 5* -> 3* |
|---|---|
| hh75_EpsilonZeta*_shell_01 | ATTCTCCGAGAGACTCCCTTAGAATCCTGGGCCTC |
| hh75_EpsilonZeta*_shell_02 | TTACATCGGGAGAAACAATAAAGGGGGACCAAGCTCATTTGA |
| hh75_EpsilonZeta*_passive_Epsilon_01 | TCGGTGCTGAAAACATAGCGATCAGATGAATATTTTTT |
| hh75_EpsilonZeta*_passive_Epsilon_02 | TTTTTTAGATTTTCAGGTTTAACGTAGCTTAGTTGGGATTGTCGG |
| hh75_EpsilonZeta*_passive_Epsilon_03 | TTTTTACAGTAACAGTACCGAAATAAACATTGATTAAGGTTGTAATAGACTT |
| hh75_EpsilonZeta*_passive_Epsilon_04 | GCACGTAAAAGAAATTGCGTTTTT |
| hh75_EpsilonZeta*_2nt_intrusion_Epsilon_01 | TCGGTGCTGAAAACATAGCGATCAGATGAAT |
| hh75_EpsilonZeta*_2nt_intrusion_Epsilon_02 | CAACAGTAACAGTACCGAAATAAACATTGATTAAGGTTGTAATAGACTT |
| hh75_EpsilonZeta*_2nt_intrusion_Epsilon_03 | GATTTTCAGGTTTAACGTAGCTTAGTTGGGATTGTCGG |
| hh75_EpsilonZeta*_2nt_intrusion_Epsilon_04 | GCACGTAAAAGAAATTGCGGG |

**Table S14: Staples for cross-configurations of HH50:** Some staples from configuration 1 and 2 are also needed, *cf.* Table S3 and Table S9

| name | sequence 5* -> 3* |
|---|---|
| hh50_Gamma*Delta_shell_01 | TACCAGATGCAAAAGAAGTTTGAGCAATTTTCA |
| hh50_Gamma*Delta_shell_02 | TTTTTGAACGGTGTACAGACCAGTTTTGAAAGAGGACAGATTTTTT |
| hh50_Gamma*Delta_shell_03 | CCGGATAGCGCATAGGGGATTTTTTATGGAGATGA |
| hh50_Gamma*Delta_shell_04 | AGCTGCTCATTCAGTGAATAAGGCTTGCCCTCGTT |
| hh50_Gamma*Delta_shell_05 | TTTTTACCAACTCATTACCCATAAGGGAATTCTGC |
| hh50_Gamma*Delta_shell_06 | CCAGATGTAAGAGCAACACTAAGGGGGTCCAGCGA |
| hh50_Gamma*Delta_shell_07 | TTTCTTTAGATCCGAACGAGGTCAATCAAATCAACACAAGAA |
| hh50_Gamma*Delta_shell_08 | CATACATTAGAGTCTTGCCAGTCATAACCCTGACGAGAAACA |
| hh50_Gamma*Delta_passive_Gamma*_01 | AATGTGCCACTCGCGGCTGGCTGACCTTCATCATTTTT |
| hh50_Gamma*Delta_passive_Gamma*_04 | TTTTTAGAGTAATCTTGGTAACAA |
| hh50_Gamma*Delta_2nt_intrusion_Gamma*_01 | AATGTGCCACTCGCGGCTGGCTGACCTTCATCAGC |
| hh50_Gamma*Delta_2nt_intrusion_Gamma*_02 | ATTACGAGGCATAGCGATTTTTGGGAAGAAGC |
| hh50_Gamma*Delta_2nt_intrusion_Gamma*_03 | TGAAATCTACGTTAATAAAGGACGTAAGAACTGGCTCA |
| hh50_Gamma*Delta_2nt_intrusion_Gamma*_04 | CCAGAGTAATCTTGGTAACAA |
| hh50_Gamma*Delta_passive_Delta_01 | ATCAGTTTTAAACTAACTTTAATCATTTTTT |
| hh50_Gamma*Delta_passive_Delta_02 | TTTTTTGTGAATTACCTTAACGAGTAGTAAATTGGGTTTTT |
| hh50_Gamma*Delta_passive_Delta_03 | TTTTTCTTGAGATGGTTTAATTTCTTGACCCAATAGTAGAGTATC |
| hh50_Gamma*Delta_2nt_intrusion_Delta_01 | ATCAGTTTTAAACTAACTTTAATCATGA |
| hh50_Gamma*Delta_2nt_intrusion_Delta_02 | AATGTGAATTACCTTAACGAGTAGTAAATTGGGGA |
| hh50_Gamma*Delta_2nt_intrusion_Delta_03 | TACTTGAGATGGTTTAATTTCTTGACCCAATAGTAGAGTATC |
| hh50_Zeta*Alpha_shell_01 | TACCAGATGCAAAAGAAGTTTGAGCAATTTTCA |
| hh50_Zeta*Alpha_shell_02 | TTTTTGAACGGTGTACAGACCTATTGAAAGAGGACAGATTTTTT |
| hh50_Zeta*Alpha_shell_03 | GATATTCATTACCCAAATCAACGATTGGGCTTGAGCCTCGTT |
| hh50_Zeta*Alpha_shell_04 | TTTTTACCAACTATCTTGACATAAGGGAATTCTGC |
| hh50_Zeta*Alpha_shell_05 | ACTTTAATAAGAGCAACACTAAGGGGGTCCAGCGA |
| hh50_Zeta*Alpha_shell_06 | TTTCTTTAGATCCGAACGAGGTCAATCAAGAACCG |
| hh50_Zeta*Alpha_shell_07 | CATACATTAGAGTCTTGCCAGTCATAACATGGTTTAATTTCA |
| hh50_Zeta*Alpha_passive_Zeta*_01 | AATGTGCCACTCGCTAGGCTGGCTTTTT |
| hh50_Zeta*Alpha_passive_Zeta*_04 | TTTTTGACCTTCATCAAGAGAGGCGCAGGGGATTTTTTATGGAGATGA |
| hh50_Zeta*Alpha_2nt_intrusion_Zeta*_01 | AATGTGCCACTCGCTAGGCTGGCAC |
| hh50_Zeta*Alpha_2nt_intrusion_Zeta*_02 | ATTACGAGGCATAGTCATTGTCATTATACCAGTCAGGACGGT |
| hh50_Zeta*Alpha_2nt_intrusion_Zeta*_03 | AGTTGGGAAGAAAACCTCTCTGAATTCG |
| hh50_Zeta*Alpha_2nt_intrusion_Zeta*_04 | CCTGACCTTCATCAAGAGAGGCGCAGGGGATTTTTTATGGAGATGA |
| hh50_Zeta*Alpha_passive_Alpha_01 | GTACAACGGCTTGCCCTGACGAGAAACACCTTTTT |
| hh50_Zeta*Alpha_passive_Alpha_02 | TTTTTAGAACGAGTAGTAATAACAAAGCTGCTCATTCTTTTT |
| hh50_Zeta*Alpha_passive_Alpha_03 | TTTTTAGTGAATAAGGAGATTTAGCGAGAGGCTTTCGACGAT |
| hh50_Zeta*Alpha_2nt_intrusion_Alpha_01 | GTACAACGGCTTGCCCTGACGAGAAACACCTG |
| hh50_Zeta*Alpha_2nt_intrusion_Alpha_02 | CCAGAACGAGTAGTAATAACAAAGCTGCTCATTCTA |
| hh50_Zeta*Alpha_2nt_intrusion_Alpha_03 | AAAGTGAATAAGGAGATTTAGCGAGAGGCTTTCGACGAT |

**Table S15: z-connections staples** of all 4 configurations for the left and right attachment sites and the respective connectors with and without toeholds and the invaders

| name | sequence 5' -> 3' |
|---|---|
| left_passive_01 | TTTTTGACAGGAACGGTACGCGATTAAAGGGATTTTATTTT |
| left_passive_02 | TTTTTGGAAATACCTACATTTTGAGACCAGTAATTTTT |
| left_passive_03 | TTTTTAACACCGCCTGCCCCTCAATCTTTTT |
| left_passive_04 | TTTTTAATATCTGGTCAGTTGTATCAAA |
| left_passive_05 | TTTTTTGGTTTGCCCCAGCAGAGCAAGCGGTCCACGCTTTTT |
| left_passive_06 | TTTTTCAGTTTGGAACAAGAGGGGTTGAGTGTTGTTCTTTTT |
| right_passive_01 | TGTTCGCCACTGGTGACCTGGAAGAGTTTTT |
| right_passive_02 | TTTTTTGACGACTGGGGATTTCAGAGCAGGCAATGCATTTTT |
| right_passive_03 | TTTTTTGAACCACCAGGCTATATCATATATGTGTTTTT |
| right_passive_04 | GCAATAAAAATGCGCCGCCTTTTT |
| right_passive_05 | TTTTTACATCGGGTTGATGCAGACATCACGAAGGTGTTTTT |
| right_passive_06 | AGATGATGACCGTACTCAATTTTT |
| zI_left_01 | TGTAGTAATGGACAGGAACGGTACGCGATTAAAGGGATTTTATTT |
| zI_left_02 | TGTAGTAATGGGAAATACCTACATTTTGAGACCAGTAATTTTT |
| zI_left_03 | TGTAGTAATGAACACCGCCTGCCCCTCAATCTTT |
| zI_left_04 | TGTAGTAATGAATATCTGGTCAGTTGTATCAAA |
| zI_left_05 | TGTAGTAATGTGGTTTGCCCCAGCAGAGCAAGCGGTCCACGCTTT |
| zI_left_06 | TGTAGTAATGCAGTTTGGAACAAGAGGGGTTGAGTGTTGTTCTTT |
| zI_right_01 | TGTTCGCCACTGGTGACCTGGAAGAGGTGGTAGTAGA |
| zI_right_02 | TTTGAAACCAGTTTCTTGTGAAGTCCGTGAAGACGGTGGTAGTAGA |
| zI_right_03 | TTTTGACGACTGGGGATTTCAGAGCAGGCAATGCAGTGGTAGTAGA |
| zI_right_04 | GCAATAAAAATGCGCCGCCGTGGTAGTAGA |
| zI_right_05 | TTTACATCGGGTTGATGCAGACATCACGAAGGTGGTGGTAGTAGA |
| zI_right_06 | AGATGATGACCGTACTCAAGTGGTAGTAGA |
| zII_left_01 | AGTAGATTGAGACAGGAACGGTACGCGATTAAAGGGATTTTATTT |
| zII_left_02 | AGTAGATTGAGGAAATACCTACATTTTGAGACCAGTAATTTTT |
| zII_left_03 | AGTAGATTGAAACACCGCCTGCCCCTCAATCTTT |
| zII_left_04 | AGTAGATTGAAATATCTGGTCAGTTGTATCAAA |
| zII_left_05 | AGTAGATTGATGGTTTGCCCCAGCAGAGCAAGCGGTCCACGCTTT |
| zII_left_06 | AGTAGATTGACAGTTTGGAACAAGAGGGGTTGAGTGTTGTTCTTT |
| zII_right_01 | TGTTCGCCACTGGTGACCTGGAAGAGGTAGTAGTGAT |
| zII_right_02 | TTTGAAACCAGTTTCTTGTGAAGTCCGTGAAGACGGTAGTAGTGAT |
| zII_right_03 | TTTTGACGACTGGGGATTTCAGAGCAGGCAATGCAGTAGTAGTGAT |
| zII_right_04 | GCAATAAAAATGCGCCGCCGTAGTAGTGAT |
| zII_right_05 | TTTACATCGGGTTGATGCAGACATCACGAAGGTGGTAGTAGTGAT |
| zII_right_06 | AGATGATGACCGTACTCAAGTAGTAGTGAT |
| zIII_left_01 | GTTAGAAGTGGACAGGAACGGTACGCGATTAAAGGGATTTTATTT |
| zIII_left_02 | GTTAGAAGTGGGAAATACCTACATTTTGAGACCAGTAATTTTT |
| zIII_left_03 | GTTAGAAGTGAACACCGCCTGCCCCTCAATCTTT |
| zIII_left_04 | GTTAGAAGTGAATATCTGGTCAGTTGTATCAAA |
| zIII_left_05 | GTTAGAAGTGTGGTTTGCCCCAGCAGAGCAAGCGGTCCACGCTTT |
| zIII_left_06 | GTTAGAAGTGCAGTTTGGAACAAGAGGGGTTGAGTGTTGTTCTTT |
| zIII_right_01 | TGTTCGCCACTGGTGACCTGGAAGAGGATGGGAAGAT |
| zIII_right_02 | TTTGAAACCAGTTTCTTGTGAAGTCCGTGAAGACGGATGGGAAGAT |
| zIII_right_03 | TTTTGACGACTGGGGATTTCAGAGCAGGCAATGCAGATGGGAAGAT |
| zIII_right_04 | GCAATAAAAATGCGCCGCCGATGGGAAGAT |
| zIII_right_05 | TTTACATCGGGTTGATGCAGACATCACGAAGGTGGATGGGAAGAT |
| zIII_right_06 | AGATGATGACCGTACTCAAGATGGGAAGAT |

| | |
|---|---|
| zIV_left_01 | TGAGGTAGAAGACAGGAACGGTACGCGATTAAAGGGATTTTATTT |
| zIV_left_02 | TGAGGTAGAAGGAAATACCTACATTTTGAGACCAGTAATTTTT |
| zIV_left_03 | TGAGGTAGAAAACACCGCCTGCCCCTCAATCTTT |
| zIV_left_04 | TGAGGTAGAAAATATCTGGTCAGTTGTATCAAA |
| zIV_left_05 | TGAGGTAGAATGGTTTGCCCCAGCAGAGCAAGCGGTCCACGCTTT |
| zIV_left_06 | TGAGGTAGAACAGTTTGGAACAAGAGGGGTTGAGTGTTGTTCTTT |
| zIV_right_01 | TGTTCGCCACTGGTGACCTGGAAGAGGTAAAGAGATA |
| zIV_right_02 | TTTGAAACCAGTTTCTTGTGAAGTCCGTGAAGACGGTAAAGAGATA |
| zIV_right_03 | TTTTGACGACTGGGGATTTCAGAGCAGGCAATGCAGTAAAGAGATA |
| zIV_right_04 | GCAATAAAAATGCGCCGCCGTAAAGAGATA |
| zIV_right_05 | TTTACATCGGGTTGATGCAGACATCACGAAGGTGGTAAAGAGATA |
| zV_right_06 | AGATGATGACCGTACTCAAGTAAAGAGATA |
| z_connector_I | CATTACTACATCTACTACCAC |
| z_connector_II | TCAATCTACTATCACTACTAC |
| z_connector_III | CACTTCTAACATCTTCCCATC |
| z_connector_IV | TTCTACCTCATATCTCTTTAC |
| z_connector_I-th7 | CTACTATCATTACTACATCTACTACCAC |
| z_connector_II-th7 | ACACTGCTCAATCTACTATCACTACTAC |
| z_connector_III-th7 | CTACCAACACTTCTAACATCTTCCCATC |
| z_connector_IV-th7 | AACTCCATTCTACCTCATATCTCTTTAC |
| z_invader_I-th7 | GTGGTAGTAGATGTAGTAATGATAGTAG |
| z_invader_II-th7 | GTAGTAGTGATAGTAGATTGAGCAGTGT |
| z_invader_III-th7 | GATGGGAAGATGTTAGAAGTGTTGGTAG |
| z_invader_IV-th7 | GTAAAGAGATATGAGGTAGAATGGAGTT |

**Table S16: 4 nt and 6 nt overlap connections** for α and α* sites.

| name | sequence 5' -> 3' |
|---|---|
| hh50_Alpha_4nt_intrusion_01 | AGATTTCGTTTATGCGAGTAGTAAATTGGGTGTT |
| hh50_Alpha_4nt_intrusion_02 | CGCCCTTGAGATGAACTTTACCTCGTTTACCAGA |
| hh50_Alpha_4nt_intrusion_03 | TAAAACGAGAAACGGAGATTTAGCGAGAGGCTTTCGACGAT |
| hh50_Alpha_4nt_intrusion_04 | GTACAACACCAGAAAATAAGGCTTGCCCTGTAAG |
| hh55_Alpha_4nt_intrusion_01 | GAAAATCTCCAAAACAAAAGGAGCCTCGCT |
| hh55_Alpha_4nt_intrusion_02 | TGAGTTAATTGTATCGGTTTATCAGAGGCATCAAAT |
| hh55_Alpha_4nt_intrusion_03 | GCAATTAAACAGCAACCATCGCGAACCAGACCGGTTTAATTCAACCTA |
| hh55_Alpha_4nt_intrusion_04 | TGACAACTTGATACTTTCGAGGTGAATTTCCATA |
| hh65_Alpha*_4nt_intrusion_01 | AATATCAGAGAGTCAGAGG |
| hh65_Alpha*_4nt_intrusion_02 | CATATGAGAGTCTGCTACAATGTAAT |
| hh65_Alpha*_4nt_intrusion_03 | GACATTCTTACCAGTAAATCAGTCACCATAAAGGTG |
| hh65_Alpha*_4nt_intrusion_04 | TAAAAGAAATAAGCAATTGTAATTTTGTT |
| hh70_Alpha*_4nt_intrusion_01 | TATCAACACTAAGAACACCCAG |
| hh70_Alpha*_4nt_intrusion_02 | CCTGTAGAAAGTACAGCTAATGCAGAACG |
| hh70_Alpha*_4nt_intrusion_03 | ATTAAATACGTTAATGATAAATAAGGCGT |
| hh70_Alpha*_4nt_intrusion_04 | AATAAACAAATTACT |
| hh50_Alpha_6nt_intrusion_01 | AGATTTCGTTTATGCGAGTAGTAAATTGGGTGTTTA |
| hh50_Alpha_6nt_intrusion_02 | CGCGCCCTTGAGATGAACTTTACCTCGTTTACCAGA |
| hh50_Alpha_6nt_intrusion_03 | GTTAAAACGAGAAACGGAGATTTAGCGAGAGGCTTTCGACGAT |
| hh50_Alpha_6nt_intrusion_04 | GTACAACACCAGAAAATAAGGCTTGCCCTGTAAGAA |
| hh55_Alpha_6nt_intrusion_01 | GAAAATCTCCAAAACAAAAGGAGCCTCGCTAA |
| hh55_Alpha_6nt_intrusion_02 | ATTGAGTTAATTGTATCGGTTTATCAGAGGCATCAAAT |
| hh55_Alpha_6nt_intrusion_03 | TGGCAATTAAACAGCAACCATCGCGAACCAGACCGGTTTAATTCAACCTA |
| hh55_Alpha_6nt_intrusion_04 | TGACAACTTGATACTTTCGAGGTGAATTTCCATATA |
| hh65_Alpha*_6nt_intrusion_01 | TATCAGAGAGTCAGAGG |
| hh65_Alpha*_6nt_intrusion_02 | CATATGAGAGTCTGCTACAATGTA |
| hh65_Alpha*_6nt_intrusion_03 | GACATTCTTACCAGTAAATCAGTCACCATAAAGG |
| hh65_Alpha*_6nt_intrusion_04 | AAAGAAATAAGCAATTGTAATTTTGTT |
| hh70_Alpha*_6nt_intrusion_01 | TCAACACTAAGAACACCCAG |
| hh70_Alpha*_6nt_intrusion_02 | CCTGTAGAAAGTACAGCTAATGCAGAA |
| hh70_Alpha*_6nt_intrusion_03 | ATTAAATACGTTAATGATAAATAAGGC |
| hh70_Alpha*_6nt_intrusion_04 | TAAACAAATTACTAG |

**Table S17: Au NP handle sequences**: Three connection sites with two handles each were designed by 3' end elongation of staples, which were otherwise used as part of connection sites. Two Ts were used as linkers between Au NP and its sequence as well as between staples and handles.

| name | sequence 5' -> 3' |
|---|---|
| AuNP_handle_hh50_Alpha | GTACAACACCAGAAAATAAGGCTTGCCCTG TT ATGTAGGTGGTAGAG |
| AuNP_handle_hh55_Alpha | TGACAACTTGATACTTTCGAGGTGAATTTC TT ATGTAGGTGGTAGAG |
| AuNP_handle_hh60_Gamma | ATTGACAAACCACCACCAGAGCC TT ATGTAGGTGGTAGAG |
| AuNP_handle_hh65_Gamma | GGCAAGGACCATCGTAAAGGTAATACCCAAAAG TT ATGTAGGTGGTAGAG |
| AuNP_handle_hh70_Beta | ACAGTAGCTTACCAGTATAAAGC TT ATGTAGGTGGTAGAG |
| AuNP_handle_hh75_Beta | ATTCTCCGAGAGACTCCCTTAGTACCTTTTACA TT ATGTAGGTGGTAGAG |
| AuNP_sequence | [thiol-C6] - TT CTCTACCACCTACAT |

**Table S18: Number of nt for specific configurations** in absolute numbers and as (*) percentage from a full, passivated structure (9408 nt as sum of core staples, ABC-shell, ABC-passive, and z-passive). Additionally, the absolute number (rounded up) and percentage of nt used per connection site. This divided the number of nt for xy-connections by six and z-connections by eight. The overall number of nt used was 17 875, which is less than the number of nt of staples needed for two whole structures.

| component | # of nt | %* | # nt / con. site | %* / con. site |
|---|---|---|---|---|
| core | 4 437 | 47.16 | | |
| ABC - shell | 2 798 | 29.74 | 467 | 4.96 |
| ABC - passivation | 1 750 | 18.60 | 292 | 3.10 |
| ABC - staple intrusions | 1 510 | 16.05 | 252 | 2.68 |
| DEF - shell | 1 390 | 14.77 | 232 | 2.47 |
| DEF - passivation | 1 118 | 11.88 | 187 | 1.99 |
| DEF - staple intrusions | 1 026 | 10.91 | 171 | 1.82 |
| cross config. - shell | 569 | 6.05 | | |
| cross config. - passivation | 375 | 3.99 | | |
| cross config. - staple intrusions | 479 | 5.09 | | |
| z - passive | 423 | 4.50 | | |
| z - handles | 1 916 | 20.37 | 240 | 2.55 |
| z - connectors | 84 | 0.89 | 21 | 0.22 |
| Σ | 17 875 | 190.00 | | |
